# Distinct mycorrhizal communities in sympatric Lepanthes orchids revealed by long-read sequencing

**DOI:** 10.64898/2026.02.17.706056

**Authors:** Piotr T. Tuczapski, Dorset W. Trapnell

## Abstract

Mycorrhizal fungi form essential mutualisms with the majority of land plants, yet their role on plant community composition and diversification remains poorly understood. Orchids provide a model system for studying these interactions because the orchid life cycle obligatorily depends on mycorrhizae. Historically, studies have emphasized the role of niche partitioning and competition avoidance resulting in distinct mycobiome compositions among coexisting orchids. Since closely related orchid species have been found to associate with similar groups of fungi, it has been speculated that different fungal species are needed for coexistence but not for speciation. However, fungi have often been examined at lower resolution levels (i.e., class, order, family, or genus). Using third-generation long-read sequencing (PacBio) of the full ITS region, we characterized fungal communities at fine scale with high accuracy. We evaluated alpha- and beta-diversity of fungal communities in four closely related, narrowly endemic epiphytic orchid species from the rapidly diversifying genus *Lepanthes* in one of the world’s richest biodiversity hotspots. Our analyses reveal that orchid species have distinct mycobiont assemblages, with differences in composition unevenly distributed across species. These results suggest that shifts in fungal partners may contribute to speciation and rapid diversification in *Lepanthes*. This study highlights the potential evolutionary role of mycorrhizal fungi in orchid diversification and demonstrates the value of high-resolution sequencing in uncovering cryptic fungal diversity.

## INTRODUCTION

A conundrum that has long intrigued the scientific community is what factors have driven the exceptionally high levels of biodiversity in the tropics. Scientists have similarly tried to understand what factors have contributed to the rapid diversification of orchids, the second largest flowering plant family, the majority of which occur in the tropics. One of the most important mutualisms in over 80% of land plants is their interaction with mycorrhizal fungi (Smith & Read 2008). Within the field of plant ecology, there is a growing recognition that this mutualism influences plant community composition, niche partitioning, and plant diversification, warranting further study (Sutherland et al. 2013).

Mycorrhizal symbionts play a vital role in facilitating nutrient uptake and are particularly beneficial in nutrient-poor conditions (Yang et al. 2016). Dust-like orchid seeds lack resources (i.e. endosperm) and thus cannot germinate without nutrients transferred by orchid mycorrhizal fungi (OMF) (Smith & Read 2008). Consequently, orchids have an obligatory relationship with OMF, but little is known about its possible role in structuring orchid communities and in driving orchid diversification. It has been previously argued that presence of distinct mycorrhizal partners in recently diverged orchid species would support the idea that mutualism plays a key role in speciation (Waterman et al. 2011).

Some have speculated that coexistence of orchid species in sympatry might be facilitated by their association with different mycorrhizal fungi, which may result in resource partitioning and reduced competition (Waterman et al. 2011, Jacquemyn et al. 2014). However, most studies of orchid-mycorrhizal relationships target terrestrial orchids from temperate regions, despite the fact that most orchids are tropical species and 80% of those are epiphytes (Givnish et al. 2015). Additionally, studies examining the role of mycorrhizal symbionts in governing coexistence of orchid species to date have focused on distantly related plant taxa that co-occur over broad geographic areas on the scale of hundreds of square meters to hundreds of square kilometers (Otero et al. 2011, Waterman et al. 2011, Jacquemyn et al. 2012a, Jacquemyn et al. 2014, Waud et al. 2016). In contrast, New World epiphytic orchids often occur in highly localized multi-species communities, sometimes occupying the same tree branches. Thus, the co-occurrence of high orchid diversity at small spatial scales in the New World tropics is a fertile testing ground of the mycorrhizal-mediated niche partitioning hypothesis.

The highest diversity of epiphytic orchids in the Neotropics is found in montane cloud forest habitat at elevations of 300-3000 m above sea level (asl) (Dodson and Escobar 1993). Epiphytic orchids are a critical component of the native flora (Dodson 1999). In Ecuador, at the elevational zone (1000–1500 m asl) that hosts the largest number of epiphytes, an estimated 53% of all epiphytic species are orchids (Küper, Kreft, Nieder, K ster, & Barthlott 2004). In the Monteverde region of Costa Rica, orchids constitute a large portion of the local flora: of the approximately 1700 plant species recorded, about 600 species are orchids (Nadkarni and Wheelwright 2000). The drivers responsible for such high orchid diversity in the region are poorly understood. Pérez-Escobar et al. (2017) have examined whether uplift of the Andes may have been a factor in Neotropical orchid diversification, an idea originally proposed by Gentry & Dodson (1987). In one of the two largest Neotropical orchid groups, subtribe Pleurothallidinae (44 genera and 5100 species, Karremans 2016), they found no correlation between paleo-elevation and diversification, probably because the most recent common ancestor of the subtribe had already been adapted to montane cloud forest habitats (*c*. 1200 m asl). In addition, the Andes Mountain Range seems to have acted as a source of new lineages for the rest of the continent, with speciation taking place in situ. Finally, uplift of the Andes probably did not act as a barrier to orchid dispersal (Pérez-Escobar et al. 2017). Taken together, these results suggest that factors other than physical barriers must have contributed to the diversification of the Pleurothallidinae, such as shifts in pollinator partners and/or fungal symbionts across an elevational gradient and heterogenous habitat (Lugo et al. 2008).

Here, we characterize the composition and diversity of mycorrhizal communities associated with four closely related, epiphytic orchid species that grow in sympatry in the New World tropics. If shifts in mycobionts play a role in speciation, differences in fungal partners should be detectable in recently diverged orchid species. Our study species belong to the evolutionarily young genus *Lepanthes* Swartz (Orchidaceae) which has the highest speciation rate in the Pleurothallidinae subtribe (Pérez-Escobar et al., 2017). The four focal species (*L. cribbii*, *L. falx-bellica, L. mentosa,* and *L. monteverdensis*) are endemic to the Monteverde region of Costa Rica and often grow sympatrically (i.e. on the same tree). This serves as a powerful system for addressing questions regarding mycorrhizae-mediated niche partitioning and whether shifts in symbionts may contribute to orchid diversification. The aims of the study are to (i) genetically identify fungal root-associates of the four orchid species; (ii) estimate variability of root-associated fungal communities (alpha diversity) within each orchid species; (iii) estimate variability in mycobiont community composition (beta diversity) among orchid taxa; and (iv) compare alpha- and beta-diversity among orchid individuals within species and among species. This research will advance our understanding of the poorly understood relationship between epiphytic orchids and their symbionts, and address a question that has long mystified biologists, namely what factors contribute to exceptionally high species diversity.

## METHODS

### Study Species

*Lepanthes* Swartz (Orchidaceae) is a Neotropical genus comprised of over 1400 species (Bogarin, Karremans, and Fernandez 2018; POWO 2024), making it one of the largest genera within the family. Members of the genus are distributed from Mexico to Bolivia and northern Brazil with most species restricted to high elevation cloud forests and paramos of the Andes Mountain Range (Pérez Escobar, Kolanowska and Parra-Sanchez 2013). The genus *Lepanthes* is estimated to have evolved over the last 2.5 million years and has the highest speciation rate in the Pleurothallidinae subtribe (Pérez Escobar et al. 2017). Scarce reports on the pollination ecology indicate that male fungus gnats are attracted to flowers via sexual deception (Blanco & Barboza 2005, Karremans et al. 2019, Peakall 2023). Many members of the genus are narrow endemics (Pridgeon 2005) and co-occur, making the genus particularly well-suited for studies addressing the question of the role of mycorrhizal specificity and niche partitioning in the diversification of sympatric species (Bayman et al. 1997). Although not documented for the focal species, some *Lepanthes* spp. flower and set fruit continuously throughout the year (Tremblay & Ackerman 2001). The four study species produce one, or occasionally two, inflorescences on the apex of the stem (called a ramicaul) (P. Tuczapski, pers. obs.). Each inflorescence can bear multiple (15 or more) flowers, typically with only one open at a time (P. Tuczapski, pers. obs.). As with most orchids, the tiny seeds are well adapted for wind dispersal (Acevedo et al. 2015, Karremans et al. 2023). While it is unknown how many seeds are produced per capsule; it is not uncommon for the fruits of other members of the family to produce thousands to millions of dust-like seeds (Ardittii & Ghani 2000). Because the seeds lack endosperm, they cannot germinate without the carbon resources supplied by their mycorrhizal symbionts. *Lepanthes monteverdensis* has been observed to occasionally produce secondary leaf-bearing stems with roots on the apex of ramicauls (P. Tuczapski pers. obs.), although most *Lepanthes* species are thought to not reproduce asexually (Karremans & Vieira-Uribe, 2020). A study of four *Lepanthes* species from Puerto Rico has estimated a highly variable lifespan with an average of 3.4 years (Tremblay 2000). The four focal species (*L. cribbii, L. falx-bellica, L. mentosa, L. monteverdensis*) are epiphytes with pendent habit, endemic to the Monteverde region of Costa Rica, which is characterized as a tropical montane cloud forest. The study species are approx. 6-10 cm in height and flowers are approximately 5 mm long. The four focal species often grow in syntopy (*i.e.*, on the same tree).

### Sampling

Collections were made at the beginning of the rainy season in July 2019 and July 2021 (**Fig. 1**). Root samples were collected from 10 adult orchids per population, where possible, from eight sites within or near the Monteverde Cloud Forest Biological Reserve. A population is defined as all the conspecific orchids growing within a tree. Study sites are situated at 1470-1756 m asl and are separated from one another by at least 1 km and a maximum of ∼6 km. Each site encompasses one to eleven trees (i.e. phorophytes) supporting one to four species of *Lepanthes*. Samples were collected from 50 populations occupying 35 phorophytes belonging to 23 species (**Supp. Table 1**). Orchid root samples were collected from 102 individuals of *L. monteverdensis* from 11 populations (mean 9.3 individuals/population), 183 individuals of *L. falx-bellica* from 22 populations (mean of 8.3 individuals/population), 74 individuals of *L. cribii* from 10 populations (mean of 7.4 individuals/population) and 50 individuals of *L. mentosa* from 7 populations (mean of 7.1 individuals/population) (**Supp. Table 2**). Roots were immediately cleaned, surface sterilized in 10% bleach solution for 5 minutes, stored in 2% CTAB buffer and snap-frozen in liquid nitrogen for transport to the University of Georgia for analysis.

**Figure 1.**
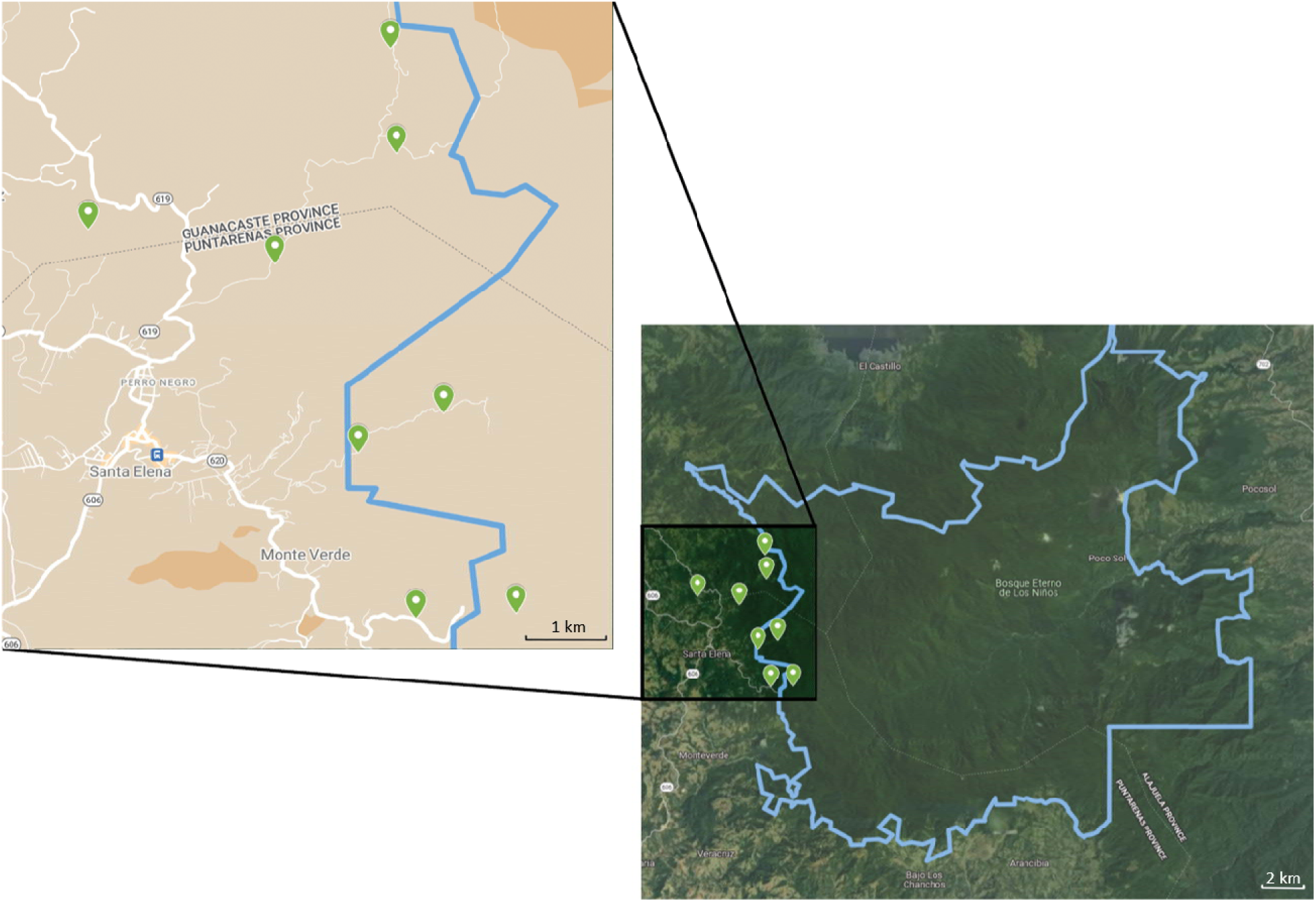
Map of study sites in the Monteverde Cloud Forest Biological Reserve. (outlined in blue). Green pins indicate sampled locations, which span ∼6 km. Additional areas to the north and south of the study sites were searched for the four study species without success. Source: Map data ©2025 Google Imagery ©2025 Airbus, CNES/Airbus, Landsat/Copernicus, Maxar Technologies.

Samples were also collected from tree bark that was in direct contact with sampled orchid roots (**Supp. Table 3**). Bark samples were immediately snap-frozen in liquid nitrogen for transport to the University of Georgia for analysis.

### Sequencing Library Preparation

Genomic DNA (fungal and orchid) was extracted from macerated root tissue using a modified CTAB protocol (Doyle 1991). Genomic DNA (fungal and tree) was also extracted from bark samples (see **Supp. Table 3** for details).

DNA quality and quantity were evaluated using an ND-1000 Nanodrop® spectrophotometer. In order to capture the broadest spectrum of mycorrhizal taxonomic units possible, the entire fungal internal transcribed spacer (fITS) ITS1-5.8S-ITS2 region was PCR amplified with two primer sets in two independent reactions. The first was a universal fungal primer set, ITS1-F_KYO2 (Toju et al. 2012) and ITS4 (White et al. 1990), which allows non-discriminating amplification of the fITS. This primer pair offers improved non-biased coverage across diverse taxonomic groups of fungi while being relatively selective for fungal ITS sequences rather than plant ITS sequences (Toju et al. 2012). The second primer set used is taxon-specific ITS4-Tul which targets orchid-associated *Tulasnella* (Taylor & McCormick 2008) paired with ITS1-F_KYO2. We designed forward and reversed primers from both primer sets to be fused with PacBio barcodes and applied combinatorial dual-indexed barcoding system in the library preparation, following the approach employed previously in TruSeq and NextEra Illumina library preparation methods (Glenn et al. 2019). Thus, each sample was amplified twice - once with each primer set - and assigned a unique barcode combination per primer set, effectively creating two separate libraries. PCR reactions were carried out in 25 μL reactions containing 12.25 μL molecular grade ddH2O, 5 μL 5x Kapa HiFi buffer, 0.75 μL 10 μM Kapa dNTP’s mix, 0.5 μL Kapa HiFi DNA polymerase (Roche, Basel, Switzerland), 2 μL of each forward and reverse 5 μM primers, and 2.5 μL of each sample genomic DNA (10, 5, 2.5. or 1.25 ng/μL dilutions). Most samples required 30 cycles to observe amplicon DNA bands on agarose gels, with some samples needing 35 or rarely 40 cycles. PCR conditions were as follows: initial denaturation (5 min at 95°C); 15 cycles of touchdown (30 s at 94°C, 30 s at 60°C, −1°C/cycle, 30 s at 72°C); followed by 10-25 cycles (30 s at 94°C, 30 s at 50°C, 30 s at 72°C); final extension (5 min at 72°C); hold (12°C). A total of 818 uniquely barcoded root samples were mixed and submitted to Georgia Genomics and Bioinformatics Core (GGBC, UG Athens, GA, RRID:SCR_010994) for sequencing on PacBio Sequel II platform. Note that we have also submitted together with these samples 239 tree bark samples (**Supp. Table 3**), which are included in the report of sequencing data as well as initial denoising and dereplication of reads in the Results section (see below), but otherwise we have not included them in the downstream analyses.

### Sequence Quality Control and Amplicon Processing

All reads were re-oriented into 5’ to 3’ direction by searching for sequences matching the forward primer, with an allowance for ≤ 5 mismatches, using DADA2 (Callahan et al. 2016) package in R version 4.3.2 (R Development Core Team 2023). Next, primers were trimmed using BBDuk with allowance for ≤ 2 mismatches. Reads from both primer sets (universal and taxon-specific) were merged for each sample. The remaining Quality Control steps were performed using DADA2. We inspected read quality profiles and applied read dereplication using an error rate learning algorithm, before implementing a core sample inference algorithm that returned amplicon sequence variants (ASVs), i.e. fungal taxonomic units, for each sample. Lastly, we removed chimeric sequences before constructing the ASV count table. Fungal taxonomy was assigned to ASVs using the General Fasta release file from the UNITE 9.0 ITS reference dataset (Abarenkov et al. 2022). The result was a dataset consisting of all fungal ASVs recovered from root extracts. Data generated by this workflow will be referred to as the general fungal (GF) dataset.

We assigned ecological guilds to all recovered fungal taxa from the GF dataset using FUNGuild in R version 4.3.2 (R Development Core Team 2023, Nguyen et al. 2016). From these, we retrieved only those ASVs belonging to guilds that have been classified as mycorrhizal. Data generated by this workflow will be referred to as the classified mycorrhizal fungi (CMF) dataset. All ASVs found with fewer than six reads in both datasets were removed from subsequent analyses due to recovery of artifact ASVs in the mock community (see Results section below).

### Sequencing/PCR Bias Control

Some commonly employed measures of alpha- and beta-diversity in ecological communities rely on the assumption that read abundance recovered from sequencing platforms corresponds to the abundance of given taxon in the sample. In an effort to test this assumption and control for potential PCR and/or sequencing biases, a mock fungal community (SynMock; Palmer et al. 2017) was included in the sequencing run. To create the SynMock community, we used 12 *Escherichia coli* strains containing pUC57 ampicillin-resistant plasmids with cloned synthetic ITS-like sequences (598 – 668 bp). First, cultures were grown in LB medium with ampicillin at 37°C overnight on a shaker table. Next, plasmid DNA was extracted from bacterial colonies using an alkaline lysis method (Cold Spring Harb Protoc; doi:10.1101/pdb.prot093344).

Concentrations of DNA were measured with an ND-1000 Nanodrop® spectrophotometer and extracts were subsequently equimolarly mixed to a final concentration of 20 ng/μL for each member in the SynMock community. This was then diluted to 10 ng/μL and PCR-amplified according to the protocol described previously.

### Alpha Diversity Analyses

Alpha diversity captures the variability in a single community (i.e. fungal ASVs) and can include richness (number of ASVs) and evenness (relative abundances of ASVs) measures. We estimated the α-diversity of fungal communities present in the roots of individual orchids using QIIME2 (Boylen et al. 2019). We repeated these calculations separately for the GF and CMF datasets.

We used Faith’s phylogenetic diversity (PD) (Faith 1992) to estimate ASV richness. Since we found that the abundance of reads returned after sequencing is not representative of the abundance of community member sequences (and presumably taxa) present in the sample before the PCR (see Results section below on SynMock), we did not rely on quantitative measures such as Shannon-Wiener diversity index (*H*) (Shannon 1948) or Pielou’s evenness (*E)* (Pielou 1966).

We built a phylogenetic tree from ASV sequences using *phylogeny* plugin’s *align-to-tree-mafft-fasttree* pipeline which performs a multiple sequence alignment using MAFFT (Katoh & Standley 2013) followed by default masking in QIIME2. For the purpose of testing for differences in PD, we treated roots of an individual plant as a sample unit and orchid species as a group. Pairs of orchid species were tested for significant differences with the Kruskal-Wallis test (Kruskal & Wallis 1952). We chose non-parametric test (instead of e.g. ANOVA) since group sizes differed and group variances may be heterogenous. To assess whether sampling effort was sufficient to fully capture α-diversity, we plotted rarefaction curves.

### Beta Diversity Analyses

Beta diversity captures the variability in community composition (*i.e*., identity of ASVs) among samples, *i.e.* how similar or different two communities are in terms of shared taxa. In this study, we compared fungal diversity among conspecific orchid species. We used ASVs found in individual plants for qualitative (Jaccard and UniFrac) analyses, where individual plant was treated as a sample unit and orchid species as a group. As with alpha diversity we did not rely on quantitative (e.g. Bray-Curtis, Bray & Curtis 1957) analysis.

The extent of overlap in fungal ASVs associating with different orchid species was visualized with principal coordinate analysis (PCoA) (Gower 1966) and non-metric multidimensional scaling plots using *phyloseq* (McMurdie and Holmes 2013) and *vegan* (Oksanen et al. 2022) packages in R version 4.3.2 (R Development Core Team 2023). The degree of similarity/dissimilarity of ASVs in the four orchid taxa was estimated qualitatively using Jaccard (S_j_) (Jaccard 1912), and UniFrac (U_AB_) (Lozupone & Knight 2005) coefficients. First, we tested for differences in the composition of fungal communities among orchid species. Jaccard Similarity Index S_j_ (converted to a distance) was calculated between each pair of orchid individuals (within- and among-*Lepanthes* species). Next, we tested whether there was a difference between fungal communities of orchid species when fungal phylogenetic relationships were considered. We calculated Unweighted UniFrac Distance among each pair of plants using the same phylogeny generated as described before. Additionally, we created unmasked GF alignment with MAFFT (Katoh & Standley 2013), followed with elimination of poorly aligned positions and divergent regions of an alignment with Gblocks ver. 0.91b (Castresana 2000), masking and phylogenetic analysis as described previously in QIIME2. We then calculated U_AB_ distances using this phylogeny. Finally, Bray-Curtis Dissimilarity Index was calculated between pairs of populations (within and among *Lepanthes* species). The significance of the differences between orchid species was tested with permutational multivariate analysis of variance (PERMANOVA) (Anderson 2014) using 999 permutations. All analyses were performed separately for the GF and CMF datasets in QIIME2 as well as packages *phyloseq*, *Biostrings*, and *vegan* in R version 4.3.2 (R Development Core Team 2023).

## RESULTS

### Sequencing, Amplicon Processing and Mock Community

The average quality score of raw pre-processed reads returned from sequencing was 98.9%. Screening of 3,141,700 reads yielded 2,826,941 demultiplexed circular consensus sequencing (CCS) reads (i.e. 90% success rate) for an average of 2,584 reads per sample. After trimming primer sequences, 98.7% of demultiplexed reads were retained.

Post implementation of the DADA2 dereplication and denoising algorithm returned 2,678,332 reads. From these, DADA2 inferred 17,116 ASVs, of which 1,778 were chimeras. Thus, 10% of all ASVs were chimeras, however these chimeric ASVs translated to only 3.4% (90,732 reads) of the total number of reads. Removal of chimeras yielded 15,338 ASVs, of which 8,058 originated from root samples (GF). After assigning these ASVs to guilds using FUNGuild, 1,714 ASVs were classified as mycorrhizal. Of these, 942 ASVs originated from root samples and constitute our CMF dataset.

To verify whether abundance of amplicons returned after PCR is representative of the abundance of community member sequences present in the sample, we used a mock fungal community, SynMock (Palmer et al. 2017). We recovered all 12 SynMock ASVs with 100% read accuracy for a total of 3,588 sequences. Subsequent to the denoising step in DADA2 and chimera removal, 3,423 sequences remained. After chimera removal there were an unequal number of reads per SynMock ASV (114 to 505 reads per ASV) despite DNA extracts having been mixed equimolarly before PCR reactions (**Supp. Table 4, Supp. Fig. 1**). For this reason, we did not use a quantitative approach that relies on read counts for calculating α-(*i.e.,* Shannon-Wiener Diversity Index or Pielou’s evenness index) and β - (*i.e.,* Bray-Curtis coefficient) diversities. In addition to the 12 true ASVs, we recovered three artifact ASVs at frequencies of 2 to 5 reads each. Thus, we removed from real samples all ASVs with fewer than 6 reads per sample as potentially erroneous, yielding 6,543 features in the GF dataset and 870 features in the CMF.

*L. monteverdensis* was assigned 3,539 GF ASVs (vs. 274 CMF ASVs), *L. falx-bellica* 2,402 GF ASVs (vs. 356 CMF ASVs), *L. cribbii* 1,083 GF ASVs (vs. 176 CMF ASVs), and *L. mentosa* 837 GF ASVs (vs. 187 CMF ASVs).

We examined rarefaction curves generated by QIIME2 and the *phyloseq* package in R to choose appropriate sampling depth for calculating α- and β-diversity metrics. In both datasets, ASV and Faith’s PD accumulation curves reached asymptotes. For the GF dataset, we used a sampling depth of 679 which retains 69% (279) of 404 filtered samples (**Fig. 2A** and **2B**). For the CMF dataset, we used a sampling depth of 445, which retains 62% (236) of 382 filtered samples (**Fig. 2C** and **2D**). When populations were used as sample units and ASV occurrences were summed, we applied a sampling depth of 100 for the CMF dataset which retains 76% (38) of 50 filtered samples (populations). For the CMF dataset, we used a sampling depth of 23 which retains 74% (37) of the filtered 50 samples.

**Figure 2.**
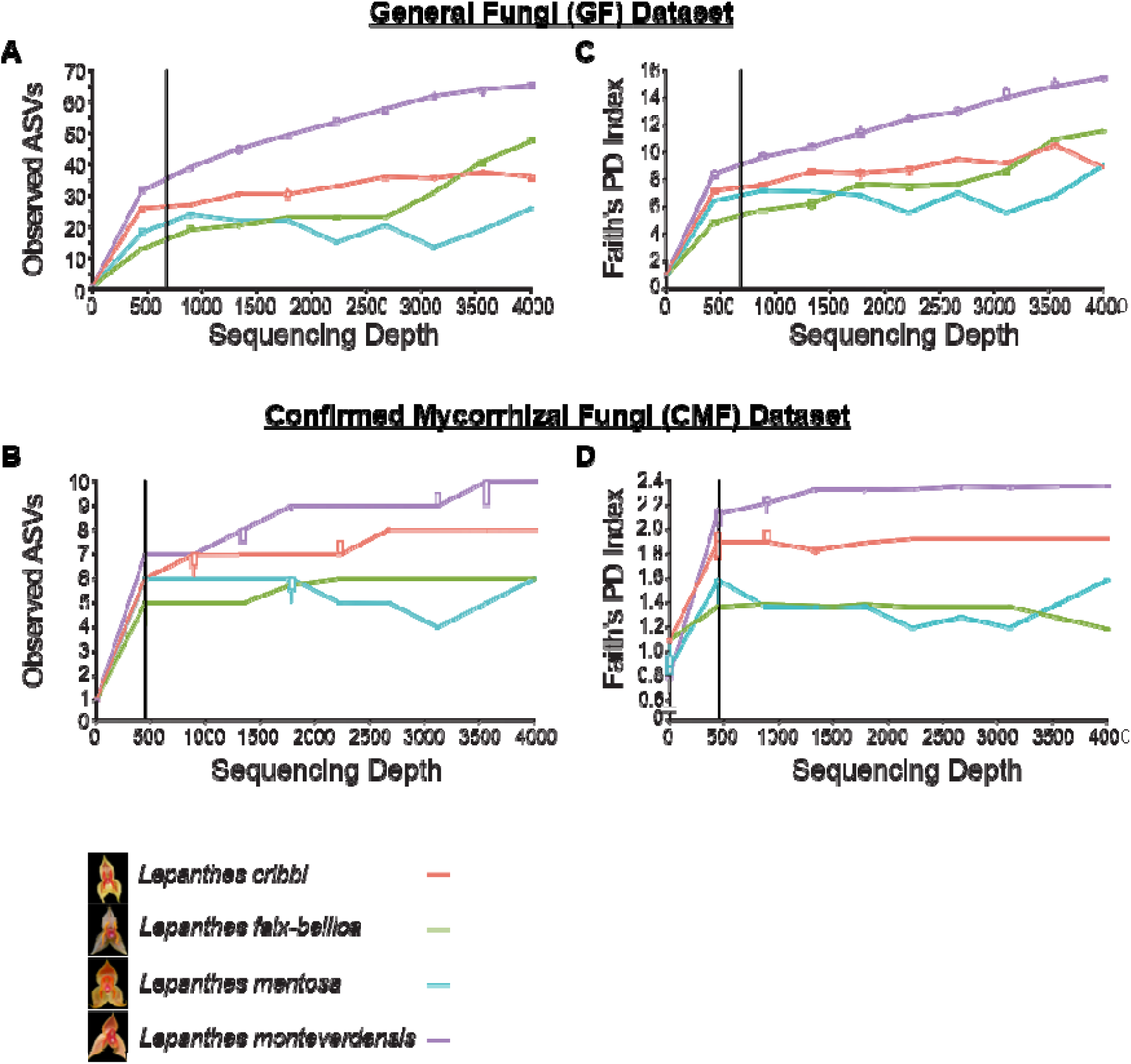
Alpha rarefaction curves. General Fungi (GF) Dataset: curves constructed based on most comprehensive dataset by averaging rarefaction curves of 404 samples over a grouping variable of an orchid species. Confirmed Mycorrhizal Fungi (CMF) Dataset: curves constructed by averaging rarefaction curves of 382 samples over a grouping variable of an orchid species. Alpha rarefaction curves show accumulation of fungal ASVs **(A & B**) and Faith’s PD Index **(C & D)** over sequencing depth. Black vertical lines indicate chosen sampling depth.

### Alpha Diversity Analyses

Orchid species had statistically significant different fungal communities in terms of their richness as shown by a Kruskal-Wallis H test performed on the GF dataset (H-value: 18.16, p-value: 0.0004) (**Table 1**). The root-associated fungal community was the richest in *L. monteverdensis*, measured as phylogenetic diversity in GF communities. Fungal richness differed most significantly between *L. monteverdensis* and *L. falx-bellica* (PD = 10.1 and PD = 6.4 respectively, Kruskal-Wallis test for Faith’s PD p-value = 0.000085). *Lepanthes monteverdensis* associated with richer GF community in comparison also to other orchid species. Specifically, significant difference in richness was found between *L. monteverdensis* and *L. cribbii* (PD = 7.5) as well as between *L. monteverdensis* and *L. mentosa* (PD = 6.7) (p-value = 0.03 and 0.004, respectively). The other three orchid species had similarly rich mycobiont communities (**Table 1**).

**Table 1.**
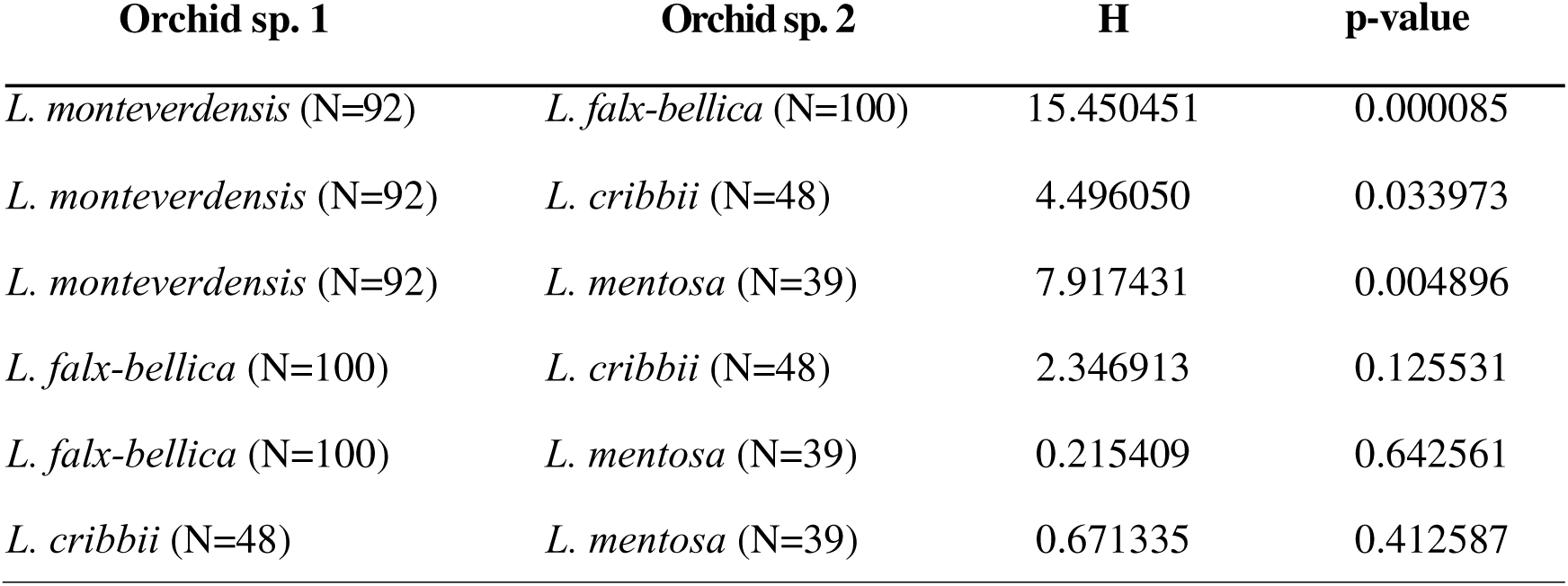
Results for Kruskal-Wallis tests applied pairwise to test for difference in fungal community richness as measured by Faith’s PD. Orchid species were treated as grouping factor. N = number of samples, and H = Kruskal-Wallis test statistic. Faith’s PD has been calculated with the most comprehensive GF dataset.

A Kruskal-Wallis H test performed on the CMF dataset also showed that orchid fungal communities varied in richness (H-value: 18.58, p-value: 0.0003) (**Table 2**). More specifically, the analysis supports results of the analysis performed with GF dataset in that *L. monteverdensis* mycobionts make the richest community (PD = 2.1). As in analysis with GF dataset, communities varied in richness between *L. monteverdensis* and *L. falx-bellica* (PD = 1.7, Kruskal-Wallis test p-value = 0.00003), as well as between *L. monteverdensis* and *L. mentosa* (PD = 1.7) (p-value = 0.01). In contrast to alpha diversity analysis with GF dataset, we found no difference between *L. monteverdensis* and *L. cribbii* (PD = 1.9) (p-value = 0.17). Additionally, we found that fungal community associated with *L. falx-bellica* was on average less rich than *L. cribbii* (PD = 1.9) (p-value = 0.04) (**Table 2**). The results of Kruskal-Wallis H test for Faith’s PD performed on GF and CMF datasets are summarized in **Fig. 4A**.

**Table 2.**
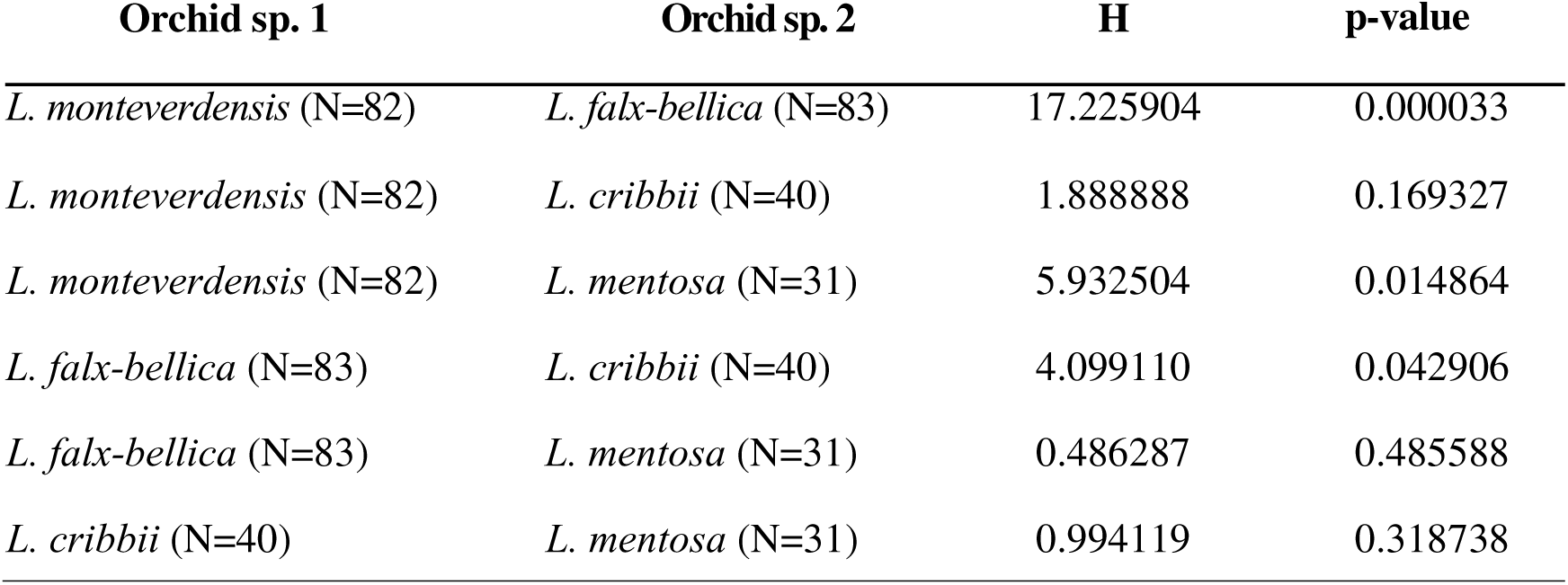
Results for Kruskal-Wallis tests applied pairwise to test for difference in fungal community richness as measured by Faith’s PD. Orchid species were treated as grouping factor. N = number of samples and H = Kruskal-Wallis test statistic. Faith’s PD has been calculated with the CMF dataset.

### Beta Diversity Analyses

We found strong evidence for differences in fungal composition (Jaccard Similarity Index S_j_) among *Lepanthes* species in the GF dataset (pseudo-F value: 1.92, p-value: 0.001, **Table 3**, **Fig. 3A** and **Supp. Figure 2A**), and the CMF dataset for every PERMANOVA test we performed (pseudo-F value: 3.29, p-value: 0.001, **Table 4**, **Fig. 3B** and **Supp. Figure 2B**). PCoA and NMDS clustering generated from Jaccard indices using both GF and CMF ASVs showed the most apparent distinct clustering for mycobionts of *L. monteverdensis* (**Fig. 3A** & **B**, **Supp. Fig. 2A** & **B**, **Supp. Table 5** & **7**).

**Figure 3.**
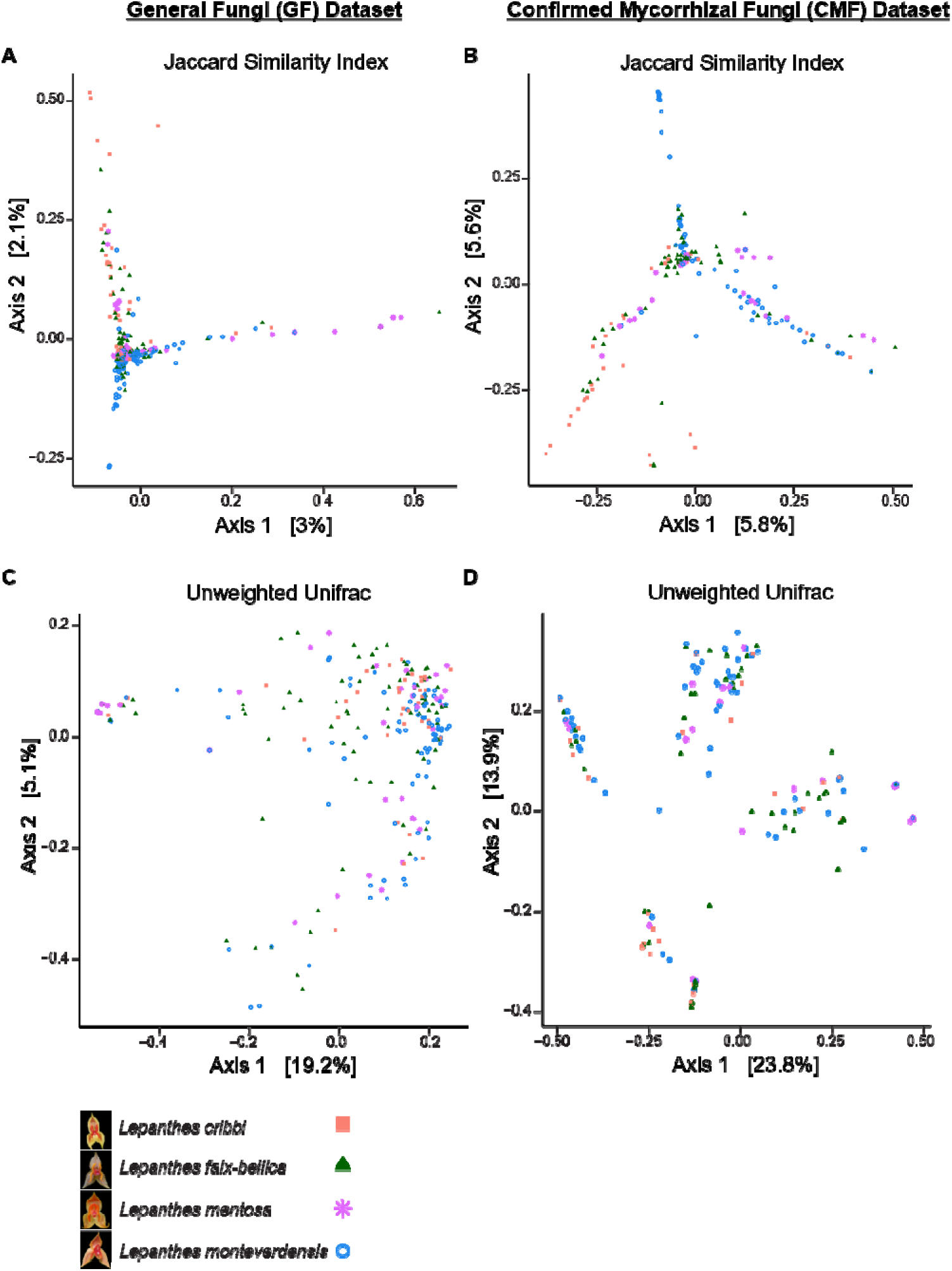
Ordination diagrams of Principal Coordinate Analysis (PCoA). General Fungi (GF) Dataset: diagrams with diversity metric scores calculated on most comprehensive GF dataset (404 samples). Confirmed Mycorrhizal Fungi (CMF) Dataset: diagrams with diversity metric scores calculated on CMF dataset (382 samples). Diagrams show PCoA calculated on Jaccard distances (converted from Similarity Indices S_j_) **(A & B)** and Unweighted UniFrac Distances U_AB_ **(C & D)** between individual orchid plant fungal communities. The classification of samples into one of the four orchid sister species is displayed by different color and symbol of individual community scores.

**Table 3.**
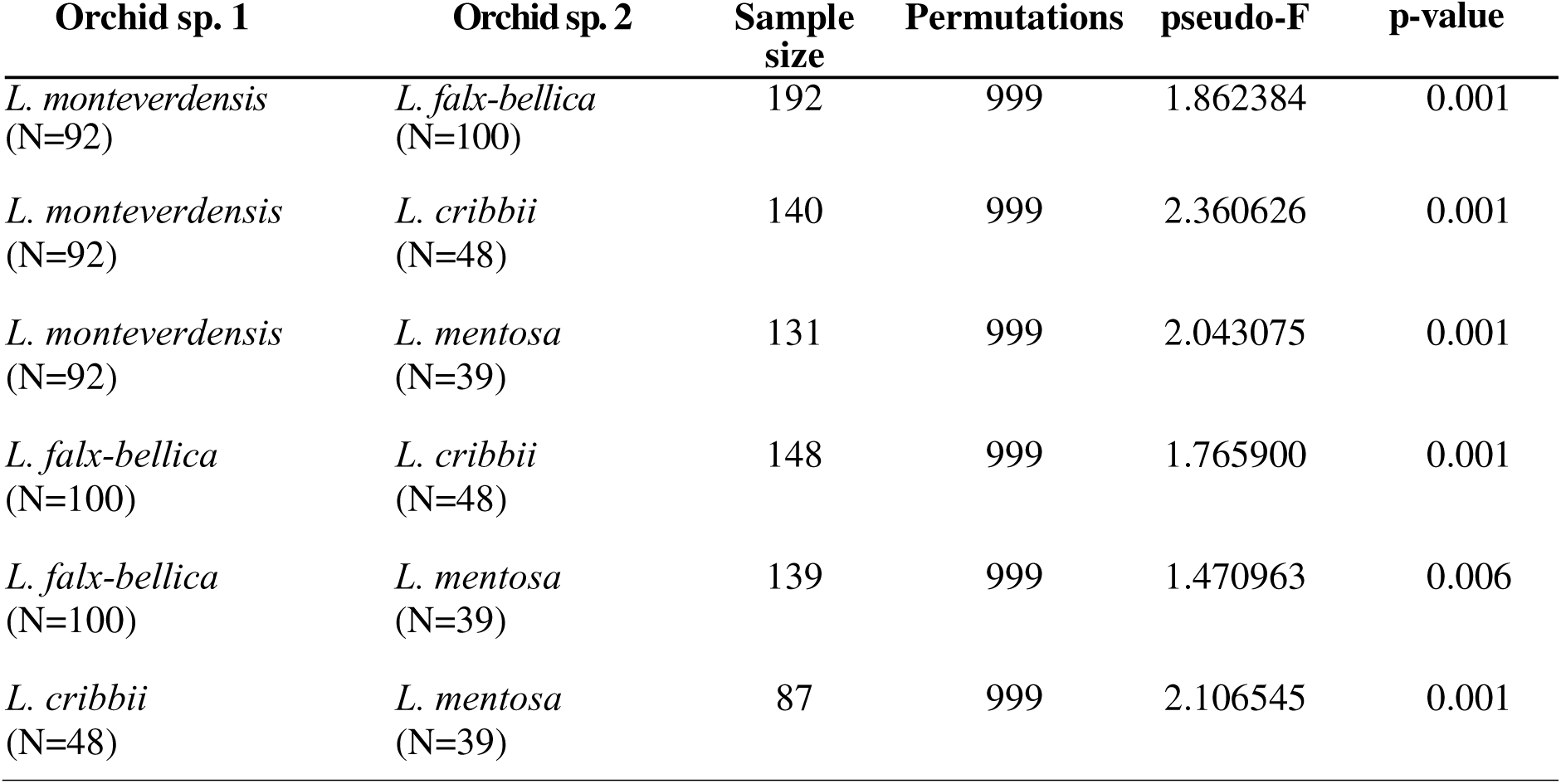
Results for pairwise permutational non-parametric multivariate analysis of variance (PERMANOVA) tests to test for difference in the qualitative composition of orchid fungal communities based on ASV identity. Analysis has been applied to the matrix of Jaccard distances (converted from Similarity Indices S_j_) calculated between orchid individual plant fungal communities. Permutations = number of permutations applied, pseudo-F = PERMANOVA test statistic. Jaccard distances have been calculated with the most comprehensive GF dataset.

**Table 4.**
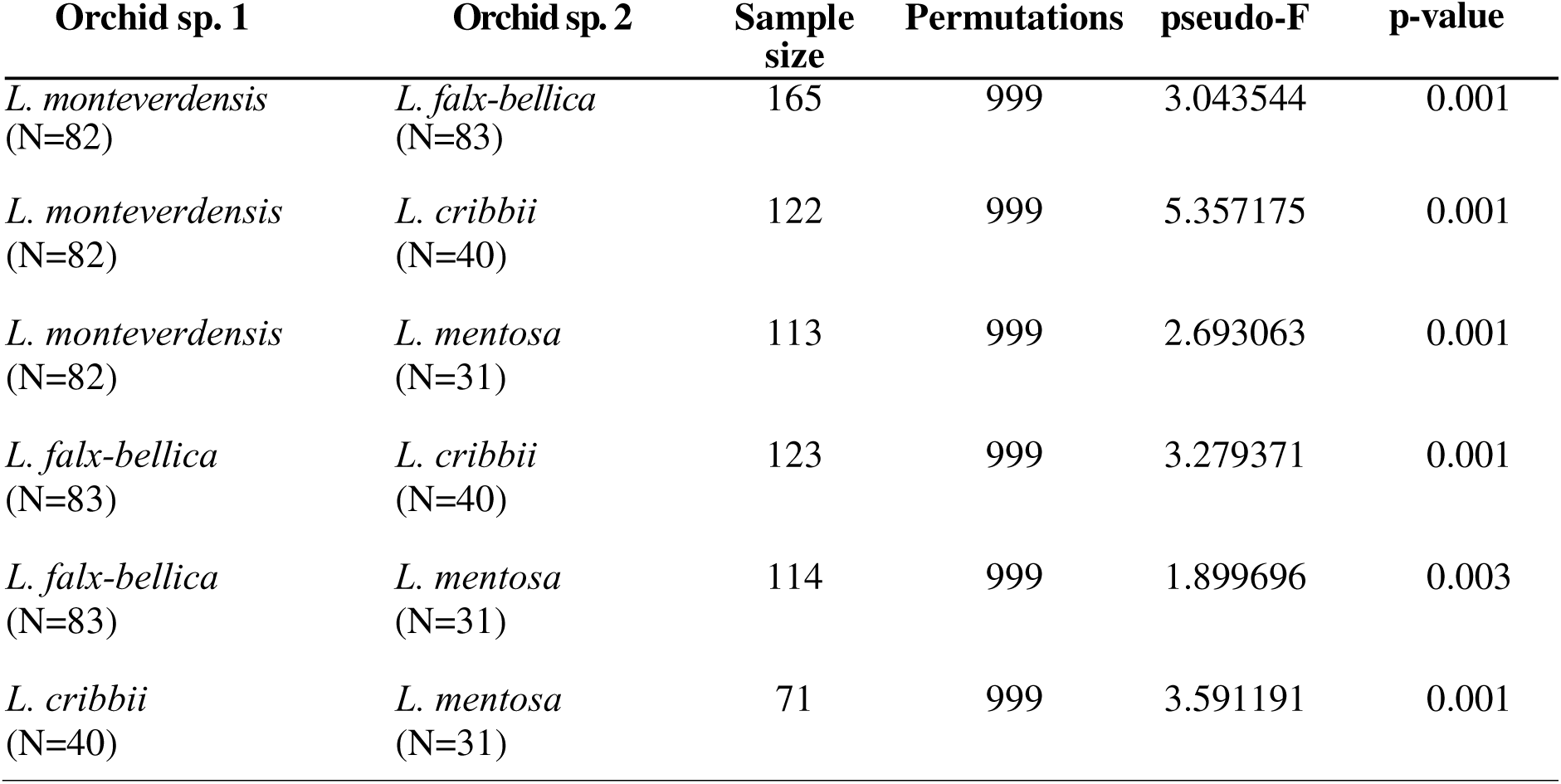
Results for pairwise permutational non-parametric multivariate analysis of variance (PERMANOVA) tests to test for difference in the qualitative composition of orchid fungal communities based on ASV identity. Analysis has been applied to the matrix of Jaccard distances (converted from Similarity Indices S_j_) calculated between orchid individual plant fungal communities. Permutations = number of permutations applied, pseudo-F = PERMANOVA test statistic. Jaccard distances have been calculated with the CMF dataset.

When fungal phylogenetic relationships are considered, we again found strong differences among *Lepanthes* species using the GF dataset (pseudo-F value: 2.13, p-value: 0.001, **Table 5**, **Fig. 3C** & **Supp. Figure 2C**) (Unweighted Unifrac Distance U_AB_). Specifically, mycobionts of *L. monteverdensis* were phylogenetically different from *L. falx-bellica* (p-value = 0.001), *L. cribbii* (p-value = 0.019) and *L. mentosa* (p-value = 0.001). There was a notable but non-significant difference between *L. falx-bellica* and *L. mentosa* (p-value of 0.079). Test results using the smaller CMF dataset (pseudo-F value: 3.84, p-value: 0.001) were consistent with results obtained from GF dataset. Specifically, *L. monteverdensis* mycobionts had different phylogenetic composition than fungi associated with *L. falx-bellica* (p-value = 0.001), *L. cribbii* (p-value = 0.049) and *L. mentosa* (p-value = 0.017) (**Table 6**). Additionally, the CMF community differed significantly between *L. falx-bellica* and *L. cribbii* (p-value = 0.039). PCoA and NMDS clustering using UniFrac distances generally followed the same pattern as that produced by Jaccard indices, except for no apparent clustering in PCoA produced with CMF dataset (**Fig.3C** & **D, Supp. Fig. 2C** & **D, Supp. Table 6** & **8**). The results of PERMANOVA test for β-diversity analyses performed on GF and CMF datasets are summarized in **Fig. 4A**. PERMANOVA test conducted on U_AB_ distances calculated based on a phylogeny derived from Gblocks-selected alignment blocks corroborated with the corresponding PERMANOVA test (where poorly aligned positions were not removed prior to phylogenetic analysis) (pseudo-F value: 2.2, p-value: 0.003, **Supp. Table 9**).

**Table 5.**
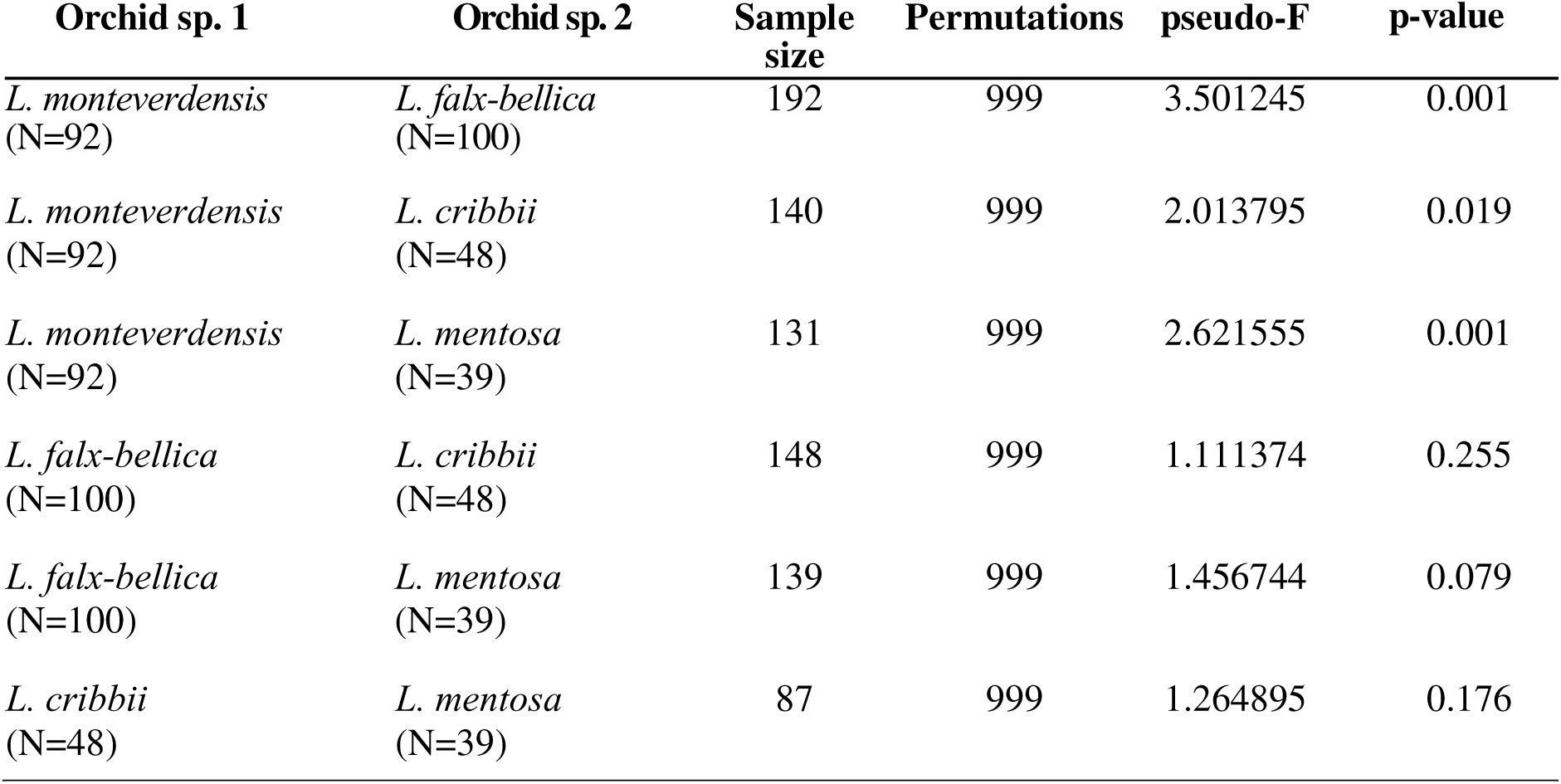
Results for pairwise permutational non-parametric multivariate analysis of variance (PERMANOVA) tests to test for difference in the qualitative composition of orchid fungal communities using phylogenetic relationships. Analysis has been applied to the matrix of Unweighted UniFrac Distances U_AB_ calculated between orchid individual plant fungal communities. Permutations = number of permutations applied, pseudo-F = PERMANOVA test statistic. Unweighted UniFrac Distances have been calculated with the most comprehensive GF dataset.

**Table 6.**
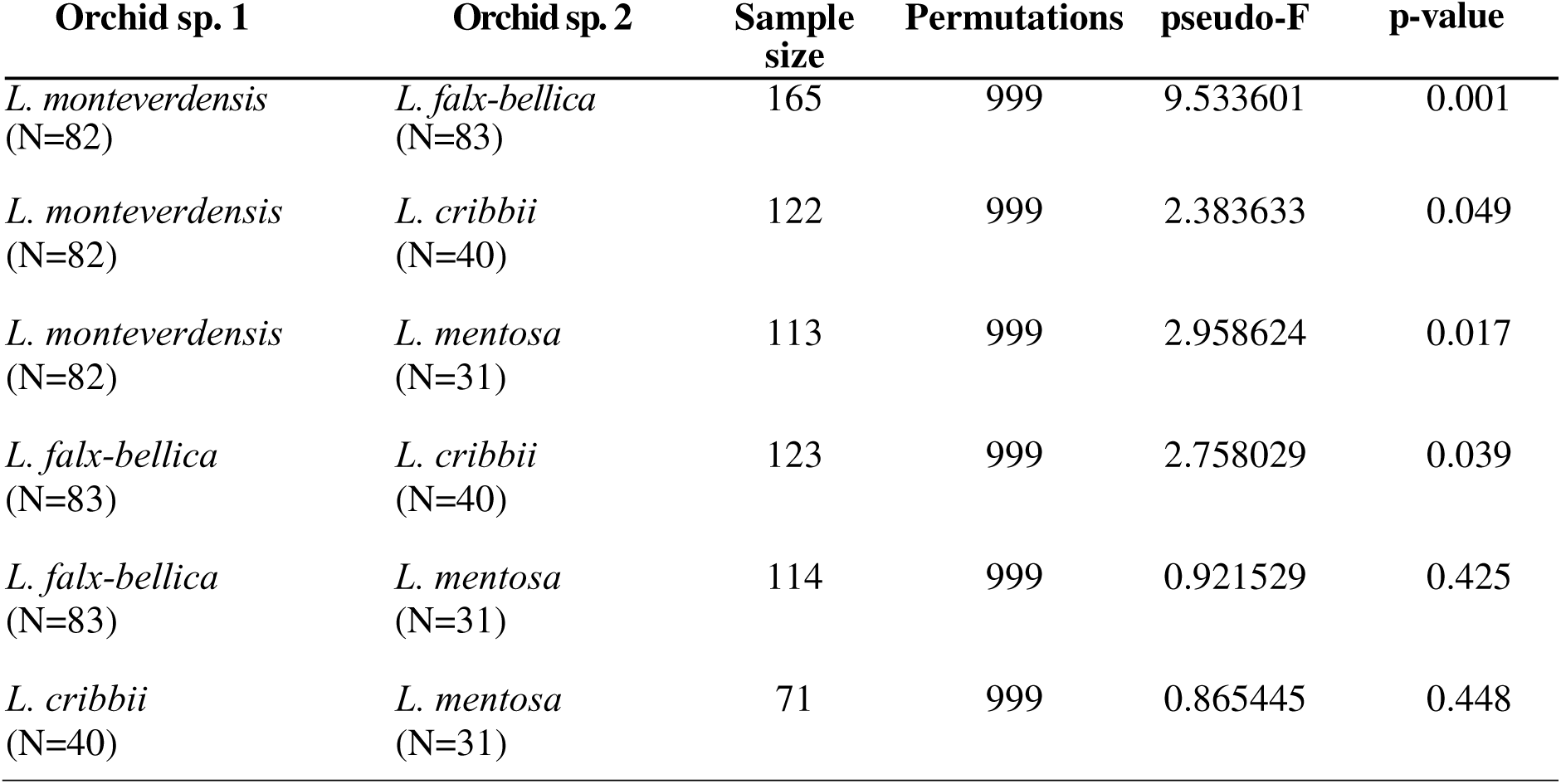
Results for pairwise permutational non-parametric multivariate analysis of variance (PERMANOVA) tests to test for difference in the qualitative composition of orchid fungal communities using phylogenetic relationships. Analysis has been applied to the matrix of Unweighted UniFrac Distances U_AB_ calculated between orchid individual plant fungal communities. Permutations = number of permutations applied, pseudo-F = PERMANOVA test statistic. Unweighted UniFrac Distances have been calculated with the CMF dataset.

**Figure 4.**
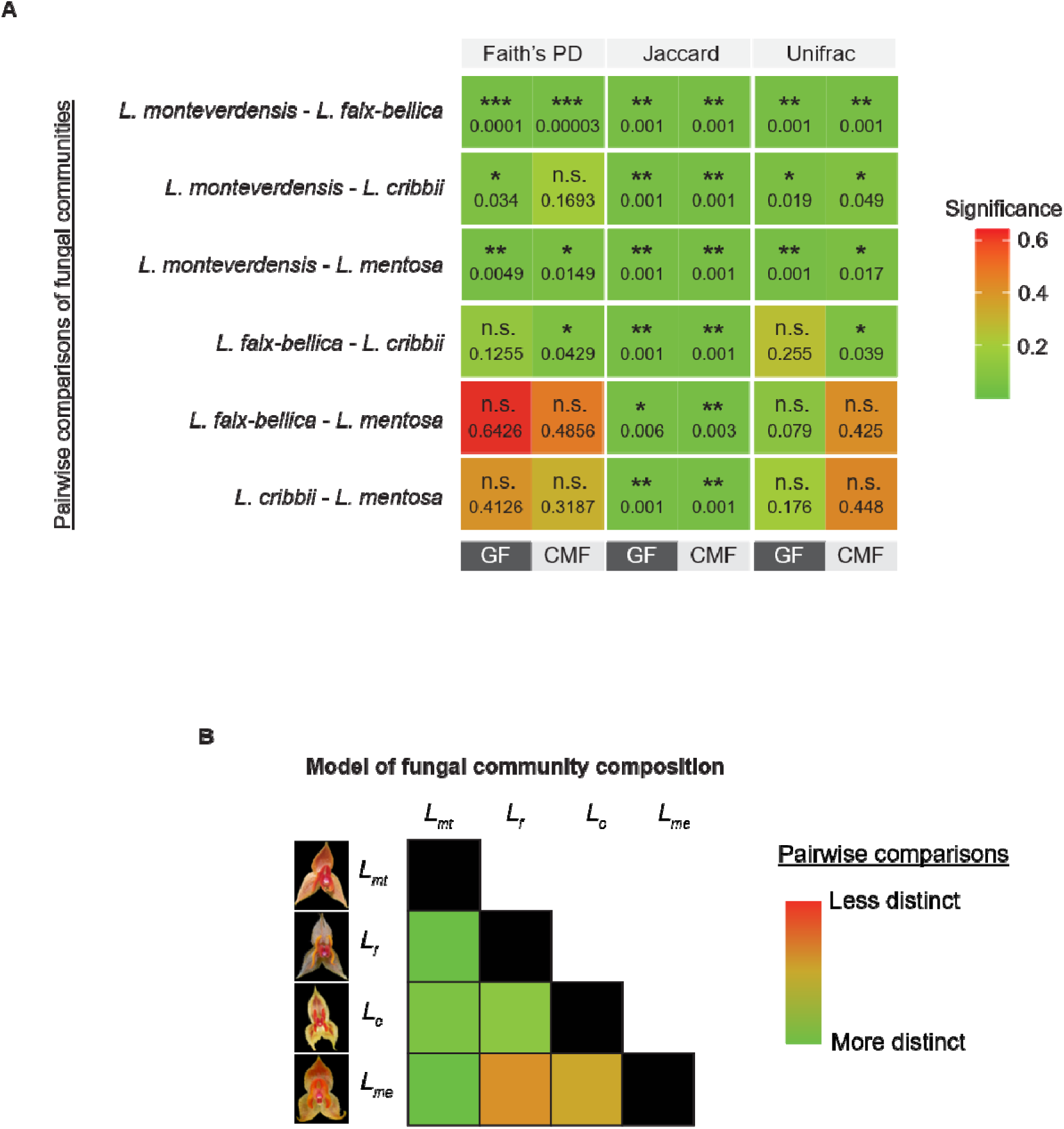
Mycorrhizal community diversity patterns associated with four *Lepanthes* sister species. (**A)** Summary of the results for the α-(Faith’s PD) and β-(Jaccard, Unweighted UniFrac) diversity analyses. GF = results of analyses performed on general fungal dataset, CMF = results of analyses performed on classified mycorrhizal dataset, Significance = p-value of respective test (Kruskal-Wallis or PERMANOVA). Significance levels: *** ≤ 0.0005, ** ≤ 0.005, * ≤ 0.05, n.s. = nonsignificant. **(B)** Conceptual model of orchid fungal communities in the order of most to least distinct.

## DISCUSSION

### Full-length ITS sequencing might more accurately characterize a fungal community

In microbial and plant ecology, researchers of fungal communities have traditionally utilized short DNA sequence fragments, typically ranging from 100 to 500 base pairs from either the ITS1 or ITS2 subregions. Due to the reduced number of base pairs in such fragments **—** compared to the full-length ITS region **—** it is thought that these minibarcodes are less informative (Kõljalg et al. 2013; Schlaeppi et al. 2016). The ITS2 subregion for example has been found to overestimate the number of fungal taxonomic units, as well as richness and/or diversity measures which could be an artifact of more random errors and hypervariability of the region (Tedersoo et al. 2018).

Third-generation high throughput sequencing technologies, such as the Pacific Biosciences (PacBio) sequencing platform, have the potential to sequence molecules up to 60 kbp and as such have opened up an exciting opportunity to utilize the whole ITS region with low error rate. Tedersoo et al. (2018) found that the full ITS dataset matched reference database records less often than the more limited ITS2 dataset, but when it did, it was more likely to be identified at a species level. This suggests that the majority of fungal DNA information comes from sequencing the ITS2 subregion and secondly that the full ITS dataset offers better taxonomic resolution. Consequently, our GF dataset (prior to mycorrhizal fungal guild assignment) likely represents true fungal taxa and as such is more informative about the fungal community than the CMF dataset which is only 13.3% of the size of GF dataset. These longer sequences have the advantage of covering both hypervariable ITS subregions as well as the more conservative flanking 5.8S, LSU and SSU subregions (Bleidorn 2016). Consequently, their use allows better estimates of fungal diversity as well as more accurate phylogenetic placement (see Kolaříková et al. 2021). Use of the full ITS barcode improves taxonomic identification at all taxonomic ranks. Tedersoo et al. (2018) found 9% higher resolution at the class level and 33% higher resolution at the genus level. Moreover, there were 2.2 - 3.2 times fewer unidentified Basidiomycota than reported with datasets of shorter ITS1 and ITS2 sequences. This is particularly relevant to our study since all major groups of orchid mycorrhizal fungi (OMF) belong to Basidiomycota.

Interestingly, the only study we are aware of where both CMF and GF datasets were assessed have reported a similar ratio of fungi recovered by both datasets as estimated by one of the ITS2-targeting primers employed (4.4-7.6 fold difference depending on orchid species, 4.5-9.6 fold in our data) while the other ITS2-targeting primer set potentially performed worse in overall and mycorrhizal fungal recovery (25.6-30.7 fold difference) (Waud et al. 2016). This study recovered about 6x fewer fungal taxonomic units in the whole fungal dataset and about 10x fewer fungal units in a confirmed mycorrhizal dataset, a finding that could be an indication of lower fungal diversity in terrestrial temperate orchids comparing to tropical epiphyte orchids.

### Mycorrhizal community diversity

The four sympatric *Lepanthes* species associate with 6,543 GF ASVs and 870 CMF ASVs. The GF ASVs may include fungi that are orchid-specific endophytes, pathogens, or yet-to-be described mycorrhizal symbionts. We suspect however that a number of GF ASVs are truly mycorrhizal but are not included in the CMF dataset for two reasons. First, taxonomical assignment inevitably depends on contributions of previous researchers to databases like UNITE and FUNGuild. Second, tropical OMF are underrepresented due to a limited number of fungal studies in equatorial regions. Finally, it is noteworthy that some recent reports have found fungi outside of widely accepted OMF groups to associate frequently with orchids, that in other plants act as endophytes or even pathogens (Li et al. 2021). These fungi may be beneficial to orchids through supporting growth, development, stress resistance, and some may possibly act as OMF. Interestingly, though, the overall picture of the fungal diversity patterns that emerges from both GF and CMF datasets remains the same. Based on the analyses, *L. monteverdensis* associates with the most distinct and diverse mycorrhizal assemblage relative to its three sympatric congeners. This is evidenced by the number of fungal ASVs as well as different community composition, phylogenetic richness, and phylogenetic distance from mycobionts associated with the other *Lepanthes* spp. While the difference in CMF richness between *L. monteverdensis* and *L. cribbii* (Faith’s PD) was not significant, there was a significant difference among these species in community qualitative composition and phylogenetic distance (Jaccard and UniFrac scores, respectively). By comparison to the mycobiont community assemblage of *L. monteverdensis*, *L. falx-bellica* shares some of its mycobiont composition with other *Lepanthes*. While all measures support its fungal diversity as separate from that of *L. monteverdensis*, this species shares some diversity with that of *L. cribbii* and *L. mentosa*. In the GF dataset, the Kruskal-Wallis test p-values for Faith’s PD comparisons among fungi in *L. falx-bellica* - *L. cribbii* individuals were low but not significant. Since this metric measures branch length in a phylogenetic tree, it simply implies that the mycorrhizal communities that these orchid species partner with have fairly similar amounts of evolutionary changes. UniFrac measures for these species were also not significantly different, indicating high amount of shared phylogeny in their fungal assemblages. However, the ASVs that make up those communities are actually different, as evidenced by the qualitative Jaccard measure. By comparison, there was limited evidence emerging from CMF dataset to support that *L. falx-bellica* mycobionts were different from those of *L. cribbii.* The PERMANOVA test detected a significant difference in the amount of unique evolutionary history of those ASVs that belong to each orchid species communities in the CMF dataset (UniFrac index), and *L. cribbii* had higher richness (Faith’s PD). In contrast, all measures in both CMF and GF datasets detected no difference between *L. mentosa* and either *L. falx-bellica* or *L. cribbii,* except for Jaccard. Thus, these results suggest that congeneric orchids investigated 1) differed in their mycobiont composition (**Fig. 4A**), and 2) those differences were not uniformly distributed among species (**Fig. 4B**).

To generate a comprehensive understanding of diversity present in the fungal communities among orchid species we selected several metrics of α- and β-diversity. We attempted to include quantitative information by summing up ASV occurrences across orchid populations (Shannon Diversity, Pielou’s Evenness and Bray-Curtis indices; **Supp. Tables 10-15**). Transforming the data for quantitative diversity estimates was necessary since read abundance does not correspond to fungal abundance as demonstrated by Palmer et al. 2017 and was further corroborated by our SynMock analysis. However, doing so greatly diminished the sample size (from 404 to 88 in the GF dataset and from 382 to 89 in the CMF dataset) as well as sampling depth (from 679 to 100 in the GF dataset and from 445 to 23 in the CMF dataset) potentially decreasing the power to detect any differences; for this reason we remain skeptical about the results of this set of analyses and thus omitted them from the summary figure (**Fig. 4**).

All four narrowly endemic *Lepanthes* spp. have been found growing on the same phorophyte in different configurations. Out of the sampled populations, *L. falx-bellica* has been found to share the same phorophyte most frequently with *L. cribbii* (17.1% of sampled trees), followed by with *L. monteverdensis* (8.6% of phorophytes). Note that our sampling data (**Supp. Table 2**) indicates that *L. falx-bellica* was growing by itself on 31.4% of phorophytes, however this value is overestimated since some of these populations did co-occur on the same tree with *L. cribbii* as well, however it was not recorded (P.Tuczapski, pers. obs.). *L. mentosa* co-occurred with *L. cribbii* on 2.9% of phorophytes. Three species, namely *L. cribbii, L. falx-bellica* and *L. monteverdensis* were found on the same phorophyte 2.9% of the time, as were all four species.

Among the instances where *Lepanthes* sp. were found growing alone, *L. monteverdensis* occurred on 17.1% of its phorophytes, *L. mentosa* on 14.3%, and *L. cribbii* on 2.9% of its phorophytes. We speculate that niche partitioning might be a mechanism driving segregation of mycobiont partners among orchid species. However, orchids vary in the degree to which their mycobiont composition differs. The frequency of co-occurrence of given orchid species with its congeners does not seem to explain fully this distribution of mycorrhizal differences. Specifically, we might expect that *L. monteverdensis*, which associates with the largest number of ASVs as well as most distinct fungal community, would have the highest co-occurrence rate in the phorophytes sampled. However, that was not the case. *L. falx-bellica* was the species that co-occurred with its congeners the most often (>31.4% vs. 14.3% in the case of *L. monteverdensis* out of 35 sampled trees), yet it shared some of its mycobiont composition with other sister taxa.

We hypothesize that our results reflect the relative recency of taxonomic divergence of some *Lepanthes* spp. versus others. It is likely that *L. monteverdensis* is the oldest of these taxa. Given reliance of orchid life cycle on their OMF, we speculate that apparent range of the degree of mycorrhizal diversity might be explained if these four sister orchid species have not been separated as taxonomic units for long enough for them to “switch over” to different mycobionts, or for mycorrhizal fungi to accumulate enough changes for tests of diversity metrics to detect. Notably, mycorrhizal partners of *L. mentosa* were different from those of *L. falx-bellica* and *L. cribbii* only as detected by Jaccard indices. This result indicates that while the compositions of the investigated fungal communities were different, phylogenetically they were similar. This could indicate that *L. mentosa* diverged more recently and is at an earlier stage of partitioning the OMF niche space, or its OMF had least time to diverge from *L. falx-bellica*’s and *L. cribbii*’s OMF. Alternatively, perhaps the pressure posed on *L. mentosa* to escape competition for resources was the lowest since it did not occupy the same niche as other orchids. While *L. mentosa*’s range overlaps with other species, it was found to be growing mainly at the lower altitude (1482-1617 m asl) sites. This may explain little difference in the mycorrhizal diversity with *L. cribbii*, which range has little overlap with that of *L. mentosa* and its upper limit (1584-1750 m asl) extends over *L. mentosa*’s range, thus possibly those two species do not commonly co-occur. However, it doesn’t explain the same result when compared to mycobionts of *L. falx-bellica,* which was found often on similar altitudes (1502-1712 m asl) and theoretically should be able to compete with *L. mentosa* more often. Finally, since *L. mentosa* was the least sampled species, we advise caution when interpreting data generated from that species. In conclusion, evidence suggests that congeneric sympatric species of orchids associate with different mycorrhizal assemblages, and those differences are more pronounced in some orchid species.

### Role of mycorrhizae on New World orchids

This study provides the first fine-scale insight utilizing high-throughput third generation sequencing technology into potential importance of mycorrhizal symbionts for niche partitioning and/or speciation in the understudied system, namely rapidly speciating epiphytic orchids of the New World tropics.

Previous studies revealed segregation of mycorrhizal communities between co-occurring orchid species and have pointed at its possible role in niche partitioning and coexistence.

However, this has been investigated mostly in a small number of plants of terrestrial, often distantly related, temperate species that occupy relatively extensive geographical regions (i.e. hundreds to thousands of square meters, sometimes spanning over thousands of kilometers; see e.g. Waterman et al. 2011, Jacquemyn et al. 2012a, Jacquemyn et al. 2012b, Jacquemyn et al. 2014, Waud et al. 2016). Moreover, prior studies have failed to account for the phylogenetic signal in the OMF assemblages and/or fine-scale taxonomic resolution of mycobionts, and their reported number is often dubious. This study provides an extreme case scenario of syntopy in which representatives of sister species often co-occur on the same phorophyte (i.e. host tree) or even branch. Epiphytic orchids are adapted to much harsher abiotic environments (*e.g*., nutrient and water limitations) relative to that experienced by terrestrial counterparts (Martos et al. 2012). Thus, epiphytic orchids have been hypothesized to experience much stronger selective pressures not only to partner with mutualistic fungi with specific functions, but also that allow to partition limited nutrient resources present in a confined space. In this context, finding an appropriate partner may be crucial for survival.

It has been speculated that plants might speciate by adapting to new fungal taxa (Thompson 1987; Cowling et al. 1990; Otero and Flanagan 2006) that would allow for access to novel sources of nutrients and ecological niches. Furthermore, it has been argued (Waterman et al. 2011) that if a given mutualism acts as an important driver of speciation, one should expect to identify different partners particularly in recently diverged species. To our knowledge, the results of our analyses provide the first report of different fungal communities in sister orchid species. It indicates that, at least in the hyper-diverse, rapidly diversifying genus *Lepanthes*, sister species indeed do tap into different mycobionts. Our finding provides an interesting contrast to previous study (Waterman et al. 2011) where authors argue that shifts in pollination traits drive plant speciation and are important for coexistence while mycorrhiza allows coexistence but not speciation. While pollinator specialization of *Lepanthes* is out of scope of this study, we acknowledge that it is likely a strong force contributing to diversification rate, given that scarce reports point at sexual deception as a mechanism employed by species of the genus (Blanco & Barboza 2005). Previous studies of sexual deception in orchids have revealed that pollination in this mating system is highly specific, with each orchid species employing males of single pollinator species (Peakall and Whitehead 2013). In contrast, ancestral character trait models performed on the second largest Neotropical orchid group, Cymbidiae, revealed no correlation between shifts in pollinator syndromes and diversification rates, which suggests that pollinator specificity might not be important for orchid fitness, and consequently, speciation (Pérez-Escobar et al. 2017). In that context, the fact that incipient sister orchid species utilize different fungal taxa might be further accelerating the process of producing new plant species. Thus, we suggest that the potential role of fungi in driving speciation in orchids should not be dismissed and deserves reconsideration. We recognize that this study is by no means a comprehensive estimation of mycobiont potential role in speciation process. For instance, it remains to be tested whether neotropical orchids associate with different fungi at different life stages, what is the extent of specialization (reciprocal selectiveness of both partners), whether there are keystone fungal nutrient providers vs. facultative partners, and whether there is a temporal effect on mycorrhizal community turnover e.g. associated with the seasons. Nevertheless, this study offers insight into the possible role of mycorrhizae in co-existence and diversification of epiphytic orchids.

## Acknowledgements

We thank A. Salazar and Monteverde Orchid Garden for field assistance; Reserva Biológica del Bosque Nuboso de Monteverde and landowners for access to populations.; D. Bogarín and M. F. Campos from Jardín Botánico Lankester, Universidad de Costa Rica provided valuable research consultations; J. Espeleta, P. A. Barriga, and Y. M. Corrales, Centro Científico Tropical for facilitating permit acquisition from the following authorities: MINAE, SINAC, CITES, USDA. D. L. Lindner for providing SynMock samples. M. McCormick, A. Chung for valuable comments on this manuscript. W. A. Haber for tree species identification; S. Rubin for laboratory assistance; K.C. Bui for figure curation. Funding was provided by the American Orchid Society, Botanical Society of America, Tinker Foundation, and the UGA Plant Biology Department to P.T.T.

## Data Accessibility

DNA sequences are deposited in the SRA (BioProject PRJNA1345165); accession numbers: SRX31179501 - SRX31179909. Scripts, along with input metadata, feature tables and ASV FASTA files necessary to conduct analyses are available in UGA Open Scholar repository (DOI: https://doi.org/10.71927/uga.27592).

## Sample metadata

Metadata are stored in the SRA (BioProject PRJNA1345165) using the MIGS: eukaryote, symbiont-associated; version 6.0 Package.

Metadata with unique sample identifier tags that can be matched to both the deposited genetic data and deposited in SRA metadata, date of sample collection, altitude and phorophyte ID are presented in Supplementary Table 16 (Supporting Information). The precision of geographic coordinates has been reduced to prevent misuse and for species protection.

All analysis scripts and R code used for data processing and statistical analyses will be available in Dryad upon publication (DOI: **[insert DOI]**).

## Benefit-Sharing Statement

Benefits Generated: The report of the research presented here has been shared with the provider of research permits and the institution facilitating permit acquisition, Centro Científico Tropical. A science communication article derived from this research (Tuczapski, P., 2021. Do *Lepanthes* orchids use fungal friends to live in harmony? *Orchids* –*The Bulletin of the American Orchid Society* Vol. 90(9)) was also shared with researchers at Centro Científico Tropical and Jardín Botánico Lankester. More broadly, our group is committed to fostering international scientific partnerships and supporting institutional capacity building.

We consulted research activities with Centro Científico Tropical, which provided the facilities, resources, and expertise. We also hired rangers from Reserva Biológica del Bosque Nuboso de Monteverde, who organized a field expedition into a remote area of the Reserve that is normally inaccessible to the public. In addition, we coordinated sample searches with guides recommended by the Reserve authorities. The contributions of all individuals to the research are described in the ACKNOWLEDGEMENTS. Lastly, as described above, all data has been shared with the broader public via appropriate biological databases.

## Author Contributions

Piotr T. Tuczapski: Conceptualization (lead); Data curation (lead); Formal analysis (lead); Sample acquisition (lead), Methodology (lead); Visualization (lead); Funding acquisition (lead); Writing-original draft (equal).

Dorset W. Trapnell: Conceptualization (supporting); Funding acquisition (supporting); Sample acquisition (supporting); Resources (lead); Supervision (lead); Writing-original draft (equal); Writing-review & editing (lead).

