## Supplemental Figures 1-2 for "Distinct mycorrhizal communities in sympatric Lepanthes orchids revealed by long-read sequencing"

SUPPLEMENTARY FIGURES

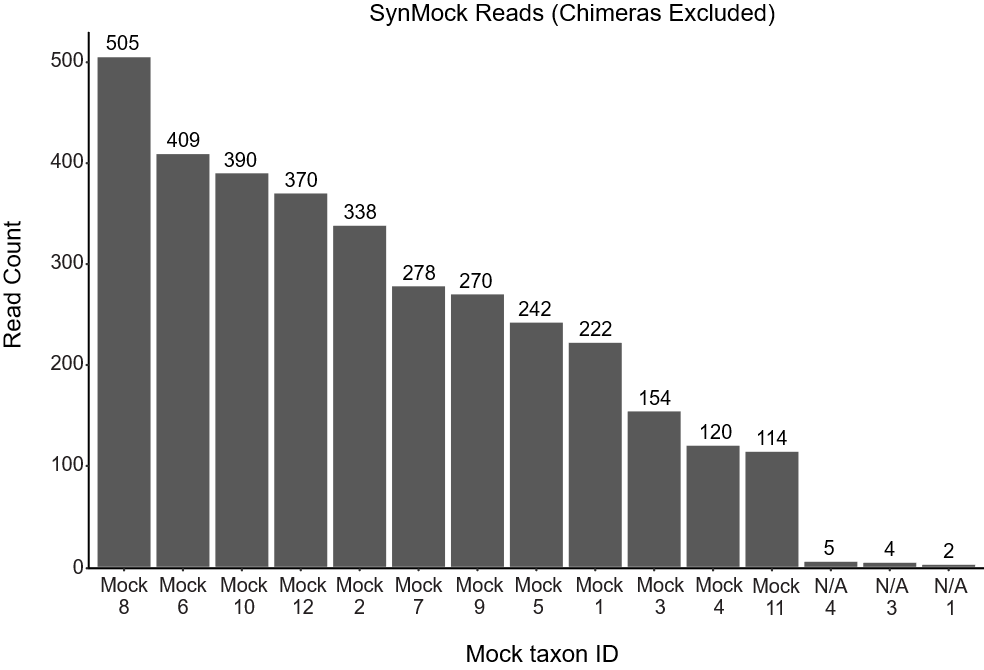

**Supplementary Figure 1.** **Recovered read count of synthetic non-biological fungal community (SynMock) sample members.** N/A = unidentified sequences not used in the community library pool.

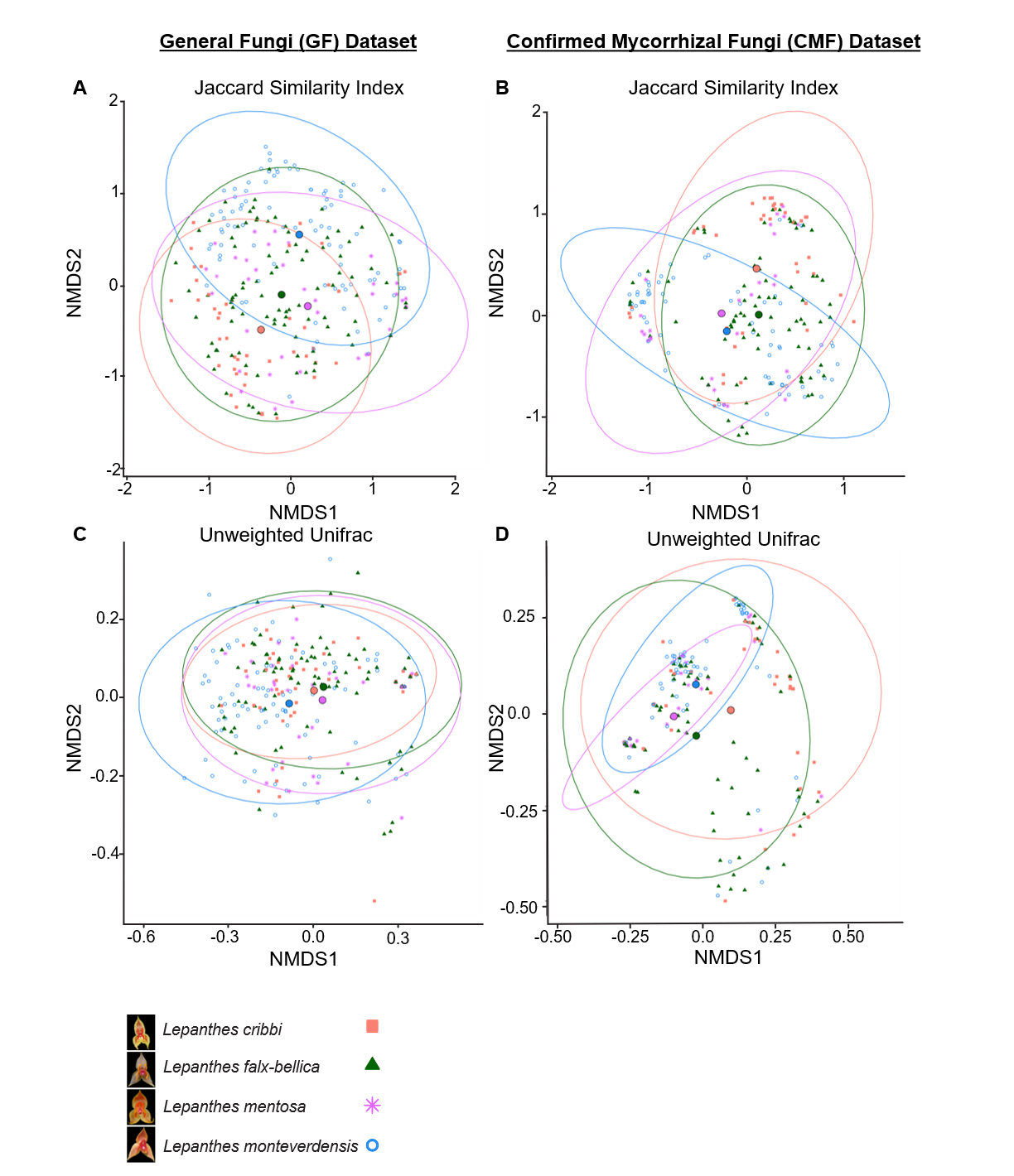

**Supplementary Figure 2.** **Ordination diagrams of Non-metric Multidimensional Scaling (NMDS).** General Fungi (GF) Dataset: diagrams with diversity metric scores calculated on most comprehensive GF dataset (404 samples). Confirmed Mycorrhizal Fungi (CMF) Dataset: diagrams with diversity metric scores calculated on CMF dataset (382 samples). Diagrams show NMDS calculated on Jaccard distances (converted from Similarity Indices S_j_) **(A & B)** and Unweighted UniFrac Distances U_AB_ **(C & D)** between individual orchid plant fungal communities. Number of dimensions of k = 5 (A & B) and k = 3 (C & D) has been selected for constructing plots. Centroids and 95% confidence ellipses have been indicated. Stress value: 0.188 (A & B), 0.142 (C), 0.167 (D).
