## Supplemental Tables 1-15 for "Distinct mycorrhizal communities in sympatric Lepanthes orchids revealed by long-read sequencing"

SUPPLEMENTARY TABLES

**Supplementary Table 1. Total number of individuals and populations sampled of four focal orchid species.**

| **Orchid species** | **No. of samples** | **No. of populations** | **No. of sites** |
| --- | --- | --- | --- |
| *L. monteverdensis* | 102 | 11 | 5 |
| *L. falx-bellica* | 183 | 22 | 7 |
| *L. cribbii* | 74 | 10 | 5 |
| *L. mentosa* | 50 | 7 | 2 |
| **Total** | **409** | **50** | **8** |

**Supplementary Table 2. Sampling data for four *Lepanthes* species.** Site ID = name of sample site, Pop ID = identity of population (i.e. tree) sampled, Tree species = phorophyte species, Lc = *Lepanthes cribbii*, Lf = *Lepanthes falx-bellica*, Lme = *Lepanthes mentosa*, Lmt = *Lepanthes monteverdensis*, *N*_O_ = number of orchid individuals sampled. The last row contains column totals.

| **Site ID** | **Pop ID** | **Tree species** | ***N_O_*** | | | |
| --- | --- | --- | --- | --- | --- | --- |
|  |  |  | **Lc** | **Lf** | **Lme** | **Lmt** |
| B | 1 | *Pouteria exfoliata, cf.* (Sapotaceae) | - | - | - | 10 |
| B | 2 | *Quercus brenesii (= Q. cortesii)* (Fagaceae) | - | 10 | - | 10 |
| C | 3 | *Miconia oerstediana* (Melastomataceae) | - | 5 | - | 10 |
| D | 4 | *Eugenia valerii* (Myrtaceae) | - | - | - | 11 |
| D | 5 | *Calyptranthes monteverdensis* (Myrtaceae) | - | 7 | - | 10 |
| H | 6 | *Calyptranthes monteverdensis* (Myrtaceae) | 10 | 9 | - | - |
| H | 7 | *Calyptranthes monteverdensis* (Myrtaceae) | 9 | 10 | - | - |
| H | 8 | *Calyptranthes monteverdensis* (Myrtaceae) | 6 | 10 | - | - |
| H | 9 | *Rondeletia buddleioides* (Rubiaceae) | 10 | 9 | - | 9 |
| H | 10 | *Calyptranthes pittieri, cf.* (Myrtaceae) | 5 | - | - | - |
| J | 11 | *Sapium rigidifolium* (Euphorbiaceae) | - | - | 5 | - |
| J | 12 | *Trichilia havanensis* (Meliaceae) | 10 | - | 6 | - |
| J | 35 | *Miconia conorufescens* (Melastomataceae) | - | - | 10 | - |
| K | 13 | *Cojoba costaricensis* (Fabaceae) | 10 | 10 | - | - |
| K | 14 | *Miconia durandii* (Melastomataceae) | - | 8 | - | - |
| K | 15 | *Eugenia austin-smithii* (Myrtaceae) | 10 | 10 | - | - |
| K | 16 | *Monteverdia recondita* (Celastraceae) | - | 7 | - | - |
| K | 17 | *Guatteria verrucosa* (Annonaceae) | - | 8 | - | - |
| K | 18 | *Sapium rigidifolium* (Euphorbiaceae) | - | 7 | - | - |
| K | 19 | *Sapium rigidifolium* (Euphorbiaceae) | - | 9 | - | - |
| N | 20 | *Guarea kunthiana* (Meliaceae) | - | 10 | - | - |
| N | 21 | *Gonzalagunia rosea* (Rubiaceae) | - | - | 8 | - |
| N | 22 | *Miconia durandii* (Melastomataceae) | - | - | 9 | - |
| N | 23 | *Miconia oerstediana* (Melastomataceae) | - | - | 10 | - |
| N | 24 | *Miconia oerstediana* (Melastomataceae) | - | - | - | 10 |
| N | 25 | *Myrsine coriacea* (Myrsinaceae) | 1 | 5 | 2 | 4 |
| N | 26 | *Pouteria exfoliata* (Sapotaceae) | - | - | - | 10 |
| N | 27 | *Viburnum venustum* (Viburnaceae) | - | - | - | 8 |
| N | 32 | *Salacia petenensis* (Hippocrateaceae) | - | 8 | - | - |
| N | 33 | *Salacia petenensis* (Hippocrateaceae) | - | 8 | - | - |
| N | 34 | *Clusia sp.* (Clusiaceae) | - | - | - | 10 |
| U | 28 | *Pouteria exfoliata* (Sapotaceae) | - | 10 | - | - |
| U | 29 | *Monteverdia recondita* (Celastraceae) | - | 10 | - | - |
| U | 30 | *Guarea rhopalocarpa* (Meliaceae) | 3 | 8 | - | - |
| U | 31 | *Pleurothyrium palmanum* (Lauraceae) | - | 5 | - | - |
| **8** | **35** | **23** | **74** | **183** | **50** | **102** |

**Supplementary Table 3. Sampling data for bark samples.** Site ID = name of sample site, Pop ID = identity of population (i.e. tree) sampled, Tree species = phorophyte species, Lc = *Lepanthes cribbii*, Lf = *Lepanthes falx-bellica*, Lme = *Lepanthes mentosa*, Lmt = *Lepanthes monteverdensis.* *N_B_* = number of phorophyte bark samples collected in proximity of the orchid sampled. The last row contains column totals. Tree bark samples were collected from the immediate vicinity of 58 individuals of *L. monteverdensis* from 6 populations (mean 9.7 bark samples/population), 95 individuals of *L. falx-bellica* from 11 populations (mean 8.6 bark samples/population), 70 individuals of *L. cribbii* from 8 populations (8.8 bark samples/population), and 11 individuals of *L. mentosa* from 2 populations (5.5 bark samples/population). Bark samples were processed for sequencing as follows. One-tenth of a gram of each bark sample was mixed with 0.8 mL of DNA extraction buffer (0.1 M Na_2_HPO_4_ (pH 5.6)) for 30s at 30 Hz. Supernatant was discarded after centrifugation (1 min, 18,000 rpm), and samples were subjected to bead milling (45s, 30 Hz) with 0.5 mL of phosphate buffer (0.1 M Na_2_HPO_4_ (pH 5.6) and 0.1 mL of 10% SDS (w/v). The supernatant was collected after centrifugation at 18,000 rpm for 3 min and mixed with 0.25 mL KCl (2.5 M) and incubated at 4°C for 15 min. The supernatant was collected after centrifugation at 18,000 rpm for 5 min, and 0.6 mL of chloroform : isoamyl alcohol (24 : 1) was added, mixed, and centrifuged at 18,000 rpm for 5 min twice. The supernatant obtained was precipitated by addition of 0.8 volume of 100% isopropanol. Finally, the pellet was recovered by centrifugation at 18,000 rpm for 30 min, washed in cold 70% ethanol and dissolved in 30 μL TLE buffer (Tris-HCl 10 mM; EDTA 0.1 mM).

| **Site ID** | **Pop ID** | **Tree species** | ***N_B_*** | | | |
| --- | --- | --- | --- | --- | --- | --- |
|  |  |  | **Lc** | **Lf** | **Lme** | **Lmt** |
| B | 1 | *Pouteria exfoliata, cf.* (Sapotaceae) | - | - | - | 10 |
| B | 2 | *Quercus brenesii (= Q. cortesii)* (Fagaceae) | - | 10 | - | 10 |
| C | 3 | *Miconia oerstediana* (Melastomataceae) | - | 5 | - | 10 |
| D | 4 | *Eugenia valerii* (Myrtaceae) | - | - | - | 9 |
| D | 5 | *Calyptranthes monteverdensis* (Myrtaceae) | - | 4 | - | 10 |
| H | 6 | *Calyptranthes monteverdensis* (Myrtaceae) | 10 | 9 | - | - |
| H | 7 | *Calyptranthes monteverdensis* (Myrtaceae) | 9 | 10 | - | - |
| H | 8 | *Calyptranthes monteverdensis* (Myrtaceae) | 6 | 10 | - | - |
| H | 9 | *Rondeletia buddleioides* (Rubiaceae) | 10 | 9 | - | 9 |
| H | 10 | *Calyptranthes pittieri, cf.* (Myrtaceae) | 5 | - | - | - |
| J | 11 | *Sapium rigidifolium* (Euphorbiaceae) | - | - | 5 | - |
| J | 12 | *Trichilia havanensis* (Meliaceae) | 10 | - | 6 | - |
| J | 35 | *Miconia conorufescens* (Melastomataceae) | - | - | - | - |
| K | 13 | *Cojoba costaricensis* (Fabaceae) | 10 | 10 | - | - |
| K | 14 | *Miconia durandii* (Melastomataceae) | - | 8 | - | - |
| K | 15 | *Eugenia austin-smithii* (Myrtaceae) | 10 | 10 | - | - |
| U | 29 | *Monteverdia recondita* (Celastraceae) | - | 10 | - | - |
| U | 30 | *Guarea rhopalocarpa* (Meliaceae) | - | - | - | - |
| U | 31 | *Pleurothyrium palmanum* (Lauraceae) | - | - | - | - |
| **7** | **19** | **16** | **70** | **95** | **11** | **58** |

**Supplementary Table 4. Sequencing data of synthetic non-biological fungal community (SynMock) sample submitted to sequencing with real samples as PCR/sequencing control.** ID = recovered community member, N with chimeras = number of sequences recovered prior to chimera removal step, N after chimera removal = number of sequences recovered post chimera removal step as implemented with DADA2, Similarity with original seq = similarity percentage with original published synthetic sequence (Palmer et al. 2017), C = concentration of a mock community member, Length = sequence length.

| ID | N with chimeras | N after chimera removal | Similarity with original seq (%) | *C* (ng/μL) | Length (bp) |
| --- | --- | --- | --- | --- | --- |
| NA1 | 2 | 2 | - | - |  |
| NA2 | 3 | - | - | - |  |
| NA3 | 4 | 4 | - | - |  |
| NA4 | 5 | 5 | - | - |  |
| NA5 | 10 | - | - | - |  |
| NA6 | 12 | - | - | - |  |
| NA7 | 18 | - | - | - |  |
| Mock 11 | 114 | 114 | 100% | 20 | 668 |
| Mock 4 | 120 | 120 | 100% | 20 | 668 |
| Mock 3 | 154 | 154 | 100% | 20 | 668 |
| Mock 1 | 222 | 222 | 100% | 20 | 668 |
| Mock 5 | 242 | 242 | 100% | 20 | 668 |
| Mock 9 | 270 | 270 | 100% | 20 | 598 |
| Mock 7 | 278 | 278 | 100% | 20 | 598 |
| Mock 2 | 338 | 338 | 100% | 20 | 668 |
| Mock 12 | 370 | 370 | 100% | 20 | 668 |
| Mock 10 | 390 | 390 | 100% | 20 | 598 |
| Mock 6 | 409 | 409 | 100% | 20 | 668 |
| Mock 8 | 505 | 505 | 100% | 20 | 598 |

**Supplementary Table 5. Tukey test results conducted after a test for multivariate homogeneity of group dispersions (p-value: 2.58e-07) implemented by *betadisper* function in *vegan* package in R.** Orchid species were treated as groups and Jaccard distances were calculated between pairs of samples (individual orchids) of the GF dataset. Diff = difference among observed mean distance-to-centroid of the levels of the grouping factor, lwr = lower bound of the 95% confidence interval, upr = upper bound of the 95% confidence interval, p adj = p-value after adjustment for the multiple comparisons.

| **Groups compared** | **diff** | **lwr** | **upr** | **p adj** |
| --- | --- | --- | --- | --- |
| *Lepanthes falx-bellica-Lepanthes cribbii* | 0.013739 | 0.006343 | 0.021135 | 1.55e-05^*^ |
| *Lepanthes mentosa-Lepanthes cribbii* | -0.00032 | -0.00933 | 0.008688 | 9.997e-01 |
| *Lepanthes monteverdensis-Lepanthes cribbii* | 0.009907 | 0.002457 | 0.017356 | 3.776e-03^*^ |
| *Lepanthes mentosa-Lepanthes falx-bellica* | -0.01406 | -0.02196 | -0.00616 | 3.855e-05^*^ |
| *Lepanthes monteverdensis-Lepanthes falx-bellica* | -0.00383 | -0.00989 | 0.002229 | 3.61e-01 |
| *Lepanthes monteverdensis-Lepanthes mentosa* | 0.010229 | 0.002277 | 0.01818 | 5.527e-03^*^ |

**Supplementary Table 6. Tukey test results conducted after a test for multivariate homogeneity of group dispersions (p-value: 1.823e-3) implemented by *betadisper* function in *vegan* package in R.** Orchid species were treated as groups and UniFrac distances were calculated between pairs of samples (individual orchids) of the GF dataset. Diff = difference among observed mean distance-to-centroid of the levels of the grouping factor, lwr = lower bound of the 95% confidence interval, upr = upper bound of the 95% confidence interval, p adj = p-value after adjustment for the multiple comparisons.

| **Groups compared** | **diff** | **lwr** | **upr** | **p adj** |
| --- | --- | --- | --- | --- |
| *Lepanthes falx-bellica-Lepanthes cribbii* | 0.007771 | -0.01352 | 0.029061 | 0.781252 |
| *Lepanthes mentosa-Lepanthes cribbii* | -0.01435 | -0.04028 | 0.01159 | 0.481644 |
| *Lepanthes monteverdensis-Lepanthes cribbii* | 0.018841 | -0.0026 | 0.040285 | 0.107378 |
| *Lepanthes mentosa-Lepanthes falx-bellica* | -0.02212 | -0.04486 | 0.000626 | 0.06001 |
| *Lepanthes monteverdensis-Lepanthes falx-bellica* | 0.011069 | -0.00638 | 0.028518 | 0.357935 |
| *Lepanthes monteverdensis-Lepanthes mentosa* | 0.033188 | 0.010298 | 0.056078 | 0.001244^*^ |

**Supplementary Table 7. Tukey test results conducted after a test for multivariate homogeneity of group dispersions (p-value: 5.959e-05) implemented by *betadisper* function in *vegan* package in R.** Orchid species were treated as groups and Jaccard distances were calculated between pairs of samples (individual orchids) of the CMF dataset. Diff = difference among observed mean distance-to-centroid of the levels of the grouping factor, lwr = lower bound of the 95% confidence interval, upr = upper bound of the 95% confidence interval, p adj = p-value after adjustment for the multiple comparisons.

| **Groups compared** | **diff** | **lwr** | **upr** | **p adj** |
| --- | --- | --- | --- | --- |
| *Lepanthes falx-bellica-Lepanthes cribbii* | 0.043293 | 0.019197 | 0.067389 | 3.399e-05^*^ |
| *Lepanthes mentosa-Lepanthes cribbii* | 0.016872 | -0.01356 | 0.047306 | 0.478603 |
| *Lepanthes monteverdensis-Lepanthes cribbii* | 0.029903 | 0.005475 | 0.054331 | 0.009412^*^ |
| *Lepanthes mentosa-Lepanthes falx-bellica* | -0.02642 | -0.05349 | 0.000647 | 0.058576 |
| *Lepanthes monteverdensis-Lepanthes falx-bellica* | -0.01339 | -0.03347 | 0.006689 | 0.312355 |
| *Lepanthes monteverdensis-Lepanthes mentosa* | 0.01303 | -0.01433 | 0.040395 | 0.606556 |

**Supplementary Table 8. Tukey test results conducted after a test for multivariate homogeneity of group dispersions (p-value: 4.043e-4) implemented by *betadisper* function in *vegan* package in R.** Orchid species were treated as groups and UniFrac distances were calculated between pairs of samples (individual orchids) of the CMF dataset. Diff = difference among observed mean distance-to-centroid of the levels of the grouping factor, lwr = lower bound of the 95% confidence interval, upr = upper bound of the 95% confidence interval, p adj = p-value after adjustment for the multiple comparisons.

| **Groups compared** | **diff** | **lwr** | **upr** | **p adj** |
| --- | --- | --- | --- | --- |
| *Lepanthes falx-bellica-Lepanthes cribbii* | -0.02144 | -0.11072 | 0.067845 | 0.925019 |
| *Lepanthes mentosa-Lepanthes cribbii* | -0.1509 | -0.26366 | -0.03813 | 0.003562^*^ |
| *Lepanthes monteverdensis-Lepanthes cribbii* | -0.09681 | -0.18732 | -0.00629 | 0.030867^*^ |
| *Lepanthes mentosa-Lepanthes falx-bellica* | -0.12946 | -0.22975 | -0.02916 | 0.005386^*^ |
| *Lepanthes monteverdensis-Lepanthes falx-bellica* | -0.07537 | -0.14977 | -0.00096 | 0.045818^*^ |
| *Lepanthes monteverdensis-Lepanthes mentosa* | 0.05409 | -0.04731 | 0.155485 | 0.512424 |

**Supplementary Table 9. Results for pairwise permutational non-parametric multivariate analysis of variance (PERMANOVA) tests to test for difference in the qualitative composition of orchid fungal communities using phylogenetic relationships.** Analysis has been applied to the matrix of Unweighted UniFrac Distances U_AB_ calculated between orchid individual plant fungal communities. Phylogeny used for U_AB_ calculations was generated based off multiple sequence alignment after eliminating poorly aligned positions and divergent regions of an alignment with Gblocks ver. 0.91b. Permutations = number of permutations applied, pseudo-F = PERMANOVA test statistic. Unweighted UniFrac Distances have been calculated with the most comprehensive GF dataset.

| **Orchid sp. 1** | **Orchid sp. 2** | **Sample size** | **Permutations** | **pseudo-F** | **p-value** |
| --- | --- | --- | --- | --- | --- |
| *L. monteverdensis* (N=92) | *L. falx-bellica* (N=100) | 192 | 999 | 3.447106 | 0.001 |
| *L. monteverdensis* (N=92) | *L. cribbii* (N=48) | 140 | 999 | 2.079164 | 0.013 |
| *L. monteverdensis* (N=92) | *L. mentosa* (N=39) | 131 | 999 | 2.797335 | 0.005 |
| *L. falx-bellica* (N=100) | *L. cribbii* (N=48) | 148 | 999 | 1.143380 | 0.228 |
| *L. falx-bellica* (N=100) | *L. mentosa* (N=39) | 139 | 999 | 1.536077 | 0.065 |
| *L. cribbii* (N=48) | *L. mentosa* (N=39) | 87 | 999 | 1.438275 | 0.118 |

**Supplementary Table 10. Results for Kruskal-Wallis tests applied pairwise to test for difference in fungal community diversity as measured by Shannon-Wiener Diversity Index *H*.** ASV occurrence in individual plants was counted. Population was treated as a sample unit and the orchid species as a group. N = number of samples, and H = Kruskal-Wallis test statistic. Shannon’s *H* has been calculated with the most comprehensive GF dataset.

| **Orchid sp. 1** | **Orchid sp. 2** | **H** | **p-value** |
| --- | --- | --- | --- |
| *L. monteverdensis* (N=11) | *L. falx-bellica* (N=16) | 0.58574 | 0.444071 |
| *L. monteverdensis* (N=11) | *L. cribbii* (N=6) | 2.766954 | 0.096229 |
| *L. monteverdensis* (N=11) | *L. mentosa* (N=5) | 0.012853 | 0.909736 |
| *L. falx-bellica* (N=16) | *L. cribbii* (N=6) | 0.348811 | 0.554787 |
| *L. falx-bellica* (N=16) | *L. mentosa* (N=5) | 0.206518 | 0.649510 |
| *L. cribbii* (N=6) | *L. mentosa* (N=5) | 1.640791 | 0.200217 |

**Supplementary Table 11. Results for Kruskal-Wallis tests applied pairwise to test for difference in fungal community diversity as measured by Shannon-Wiener Diversity Index *H*.** ASV occurrence in individual plants was counted. Population was treated as a sample unit and the orchid species as a group. N = number of samples, and H = Kruskal-Wallis test statistic. Shannon’s *H* has been calculated with the CMF dataset.

| **Orchid sp. 1** | **Orchid sp. 2** | **H** | **p-value** |
| --- | --- | --- | --- |
| *L. monteverdensis* (N=11) | *L. falx-bellica* (N=15) | 0.529939 | 0.466632 |
| *L. monteverdensis* (N=11) | *L. cribbii* (N=6) | 1.992004 | 0.158132 |
| *L. monteverdensis* (N=11) | *L. mentosa* (N=5) | 0.158619 | 0.690431 |
| *L. falx-bellica* (N=15) | *L. cribbii* (N=6) | 1.107422 | 0.292643 |
| *L. falx-bellica* (N=15) | *L. mentosa* (N=5) | 0.275321 | 0.599785 |
| *L. cribbii* (N=6) | *L. mentosa* (N=5) | 1.421254 | 0.233197 |

**Supplementary Table 12. Results for Kruskal-Wallis tests applied pairwise to test for difference in fungal community evenness as measured by Pielou’s Evenness Index *E*.** ASV occurrence in individual plants was counted. Population was treated as a sample unit and the orchid species as a group. N = number of samples, and H = Kruskal-Wallis test statistic. Pielou’s *E* has been calculated with the most comprehensive GF dataset.

| **Orchid sp. 1** | **Orchid sp. 2** | **H** | **p-value** |
| --- | --- | --- | --- |
| *L. monteverdensis* (N=11) | *L. falx-bellica* (N=16) | 0.412029 | 0.520941 |
| *L. monteverdensis* (N=11) | *L. cribbii* (N=6) | 2.929964 | 0.086949 |
| *L. monteverdensis* (N=11) | *L. mentosa* (N=5) | 0.003223 | 0.954729 |
| *L. falx-bellica* (N=16) | *L. cribbii* (N=6) | 0.719563 | 0.396287 |
| *L. falx-bellica* (N=16) | *L. mentosa* (N=5) | 0.245934 | 0.619953 |
| *L. cribbii* (N=6) | *L. mentosa* (N=5) | 0.833333 | 0.361310 |

**Supplementary Table 13. Results for Kruskal-Wallis tests applied pairwise to test for difference in fungal community evenness as measured by Pielou’s Evenness Index *E*.** ASV occurrence in individual plants was counted. Population was treated as a sample unit and the orchid species as a group. N = number of samples, and H = Kruskal-Wallis test statistic. Pielou’s *E* has been calculated with the most comprehensive CMF dataset.

| **Orchid sp. 1** | **Orchid sp. 2** | **H** | **p-value** |
| --- | --- | --- | --- |
| *L. monteverdensis* (N=11) | *L. falx-bellica* (N=15) | 0.648690 | 0.420581 |
| *L. monteverdensis* (N=11) | *L. cribbii* (N=6) | 1.992004 | 0.158132 |
| *L. monteverdensis* (N=11) | *L. mentosa* (N=5) | 0.080928 | 0.776044 |
| *L. falx-bellica* (N=15) | *L. cribbii* (N=6) | 1.105982 | 0.292957 |
| *L. falx-bellica* (N=15) | *L. mentosa* (N=5) | 0.068726 | 0.793200 |
| *L. cribbii* (N=6) | *L. mentosa* (N=5) | 1.017584 | 0.313093 |

**Supplementary Table 14. Results for pairwise permutational non-parametric multivariate analysis of variance (PERMANOVA) tests to test for difference in the quantitative composition of orchid fungal communities using phylogenetic relationships.** ASV occurrence in individual plants was recorded. Analysis has been applied to the matrix of Bray-Curtis Dissimilarity Indexes S_BC_ calculated between orchid population fungal communities. Permutations = number of permutations applied, pseudo-F = PERMANOVA test statistic. Bray-Curtis’s S_BC_ have been calculated with the most comprehensive GF dataset.

| **Orchid sp. 1** | **Orchid sp. 2** | **Sample size** | **Permutations** | **pseudo-F** | **p-value** |
| --- | --- | --- | --- | --- | --- |
| *L. monteverdensis* (N=11) | *L. falx-bellica* (N=16) | 27 | 999 | 1.233710 | 0.001 |
| *L. monteverdensis* (N=11) | *L. cribbii* (N=6) | 17 | 999 | 1.245775 | 0.001 |
| *L. monteverdensis* (N=11) | *L. mentosa* (N=5) | 16 | 999 | 1.046356 | 0.176 |
| *L. falx-bellica* (N=16) | *L. cribbii* (N=6) | 22 | 999 | 0.931400 | 0.798 |
| *L. falx-bellica* (N=16) | *L. mentosa* (N=5) | 21 | 999 | 1.021773 | 0.375 |
| *L. cribbii* (N=6) | *L. mentosa* (N=5) | 11 | 999 | 1.041885 | 0.282 |

**Supplementary Table 15. Results for pairwise permutational non-parametric multivariate analysis of variance (PERMANOVA) tests to test for difference in the quantitative composition of orchid fungal communities using phylogenetic relationships.** ASV occurrence in individual plants was recorded. Analysis has been applied to the matrix of Bray-Curtis Dissimilarity Indexes S_BC_ calculated between orchid population fungal communities. Permutations = number of permutations applied, pseudo-F = PERMANOVA test statistic. Bray-Curtis’s S_BC_ have been calculated with the CMF dataset.

| **Orchid sp. 1** | **Orchid sp. 2** | **Sample size** | **Permutations** | **pseudo-F** | **p-value** |
| --- | --- | --- | --- | --- | --- |
| *L. monteverdensis* (N=11) | *L. falx-bellica* (N=15) | 26 | 999 | 1.091479 | 0.26 |
| *L. monteverdensis* (N=11) | *L. cribbii* (N=6) | 17 | 999 | 1.216385 | 0.133 |
| *L. monteverdensis* (N=11) | *L. mentosa* (N=5) | 16 | 999 | 1.046341 | 0.312 |
| *L. falx-bellica* (N=15) | *L. cribbii* (N=6) | 21 | 999 | 0.791584 | 0.834 |
| *L. falx-bellica* (N=15) | *L. mentosa* (N=5) | 20 | 999 | 1.132687 | 0.193 |
| *L. cribbii* (N=6) | *L. mentosa* (N=5) | 11 | 999 | 1.191012 | 0.175 |
