## Supplemental Information for "Distinct mycorrhizal communities in sympatric Lepanthes orchids revealed by long-read sequencing"

**Additional File for Review but Not for Publication**

**ASVs versus OTUs for characterization of fungal communities**

A common approach in fungal research is to apply a 97% sequence similarity threshold to circumscribe Operational Taxonomic Units (OTUs) (Hughes, Petersen, & Lickey 2009), a method also used to delimit mycorrhizal species in basidiomycetes (Jacquemyn et al. 2010). It has been argued that Tulasnellaceae OTUs should be delimited using a less conservative 95% threshold due to high sequence diversity in *Tulasnella* spp. (Novotná et al. 2018). We chose instead to use the output generated by *DADA2* denoising and dereplication methods, to identify ASVs for downstream analyses for several reasons. First, the average quality score of raw sequences was reasonably high (98.9%) and all mock community ASVs were 100% accurate. Second, based on our SynMock results we reasoned that any variable length sequences introduced during PCR or sequencing should be in low frequency and/or occurrence, and even if they survived filtering, they would have little impact on α- and β-diversity patterns. Third, previous research has found that the risk of overestimating the number of fungal community members is more likely to happen with the use of ITS2 as opposed to the full ITS region (Tedersoo et al. 2018). Moreover, applying ITS sequence similarity cutoffs is likely to underestimate fungal diversity (Waterman et al. 2011). Lastly, the read coverage we obtained per sample is sufficient to recover all community members as evidenced by rarefaction curves. The GF curves reach saturation at about 15-35 (5-7 in CMF dataset) ASVs/orchid species which roughly corresponds to the approximate orchid mycorrhizae OTU number reports available in scarce studies of tropical orchids (Novotná et al. 2018; Cevallos et al. 2017). However, contrary to prior studies we recovered many more total fungal taxonomic units, a finding potentially due to more intensive sampling as well as higher sequencing coverage. Differences in methodology could also have contributed to this discrepancy, e.g. Novotná et al. (2018) sequenced only fungi that were previously successfully cultured, potentially omitting non-culturable fungi, while Cevallos et al. (2017) employed a shorter ITS2 minibarcode. Regardless, this would not affect the overall diversity estimates given that the rarefaction curve of Faith’s PD reached asymptote. Thus, in addition to the low risk of rare ASVs confounding the results, the ASVs found in the present study are likely to be biologically meaningful. That said, we recognize that some properties inherent to fungal biology might inflate ASV numbers (Palmer et al. 2017).
