## Supplemental Table 16 for "Distinct mycorrhizal communities in sympatric Lepanthes orchids revealed by long-read sequencing"

| barcode | sample name | orchid sp. |
|---|---|---|
| categorical | categorical | categorical |
| bc1003--bc1018 | LMo-3-8-B | Lepanthes monteverdensis |
| bc1001--bc1040 | LMo-3-9-B | Lepanthes monteverdensis |
| bc1007--bc1018 | LMo-3-10-B | Lepanthes monteverdensis |
| bc1001--bc1049 | LMo-3-165-B | Lepanthes monteverdensis |
| bc1001--bc1050 | LMo-3-166-B | Lepanthes monteverdensis |
| bc1001--bc1011 | LMo-3-167-B | Lepanthes monteverdensis |
| bc1001--bc1012 | LMo-3-168-B | Lepanthes monteverdensis |
| bc1001--bc1058 | LMo-3-169-B | Lepanthes monteverdensis |
| bc1005--bc1015 | LMo-3-170-B | Lepanthes monteverdensis |
| bc1005--bc1040 | LMo-3-171-B | Lepanthes monteverdensis |
| bc1005--bc1007 | LMo-5-12-B | Lepanthes monteverdensis |
| bc1005--bc1008 | LMo-5-13-B | Lepanthes monteverdensis |
| bc1016--bc1017 | LMo-5-172-B | Lepanthes monteverdensis |
| bc1005--bc1011 | LMo-5-173-B | Lepanthes monteverdensis |
| bc1005--bc1012 | LMo-5-174-B | Lepanthes monteverdensis |
| bc1006--bc1017 | LMo-5-175-B | Lepanthes monteverdensis |
| bc1003--bc1006 | LMo-5-176-B | Lepanthes monteverdensis |
| bc1006--bc1006 | LMo-5-177-B | Lepanthes monteverdensis |
| bc1006--bc1007 | LMo-5-178-B | Lepanthes monteverdensis |
| bc1007--bc1017 | LMo-5-179-B | Lepanthes monteverdensis |
| bc1006--bc1050 | LFB-5-14-B | Lepanthes falx-bellica |
| bc1006--bc1052 | LFB-5-15-B | Lepanthes falx-bellica |
| bc1006--bc1057 | LFB-5-16-B | Lepanthes falx-bellica |
| bc1006--bc1058 | LFB-5-180-B | Lepanthes falx-bellica |
| bc1003--bc1008 | LFB-5-181-B | Lepanthes falx-bellica |
| bc1006--bc1008 | LFB-5-182-B | Lepanthes falx-bellica |
| bc1007--bc1008 | LFB-5-183-B | Lepanthes falx-bellica |
| bc1008--bc1008 | LFB-5-184-B | Lepanthes falx-bellica |
| bc1008--bc1050 | LFB-5-185-B | Lepanthes falx-bellica |
| bc1009--bc1017 | LFB-5-186-B | Lepanthes falx-bellica |
| bc1008--bc1057 | LFB-6-17-C | Lepanthes falx-bellica |
| bc1011--bc1017 | LFB-6-18-C | Lepanthes falx-bellica |
| bc1012--bc1017 | LFB-6-166-C | Lepanthes falx-bellica |
| bc1009--bc1040 | LFB-6-167-C | Lepanthes falx-bellica |
| bc1007--bc1009 | LMo-6-19-C | Lepanthes monteverdensis |
| bc1009--bc1049 | LFB-6-20-C | Lepanthes falx-bellica |
| bc1009--bc1009 | LMo-6-21-C | Lepanthes monteverdensis |
| bc1009--bc1052 | LMo-6-24-C | Lepanthes monteverdensis |
| bc1009--bc1057 | LMo-6-25-C | Lepanthes monteverdensis |
| bc1009--bc1058 | LMo-6-26-C | Lepanthes monteverdensis |
| bc1003--bc1010 | LMo-6-27-C | Lepanthes monteverdensis |
| bc1006--bc1010 | LMo-6-28-C | Lepanthes monteverdensis |

| | | |
|---|---|---|
| bc1007--bc1010 | LMo-6-29-C | Lepanthes monteverdensis |
| bc1008--bc1010 | LMo-6-164-C | Lepanthes monteverdensis |
| bc1009--bc1010 | LMo-6-165-C | Lepanthes monteverdensis |
| bc1010--bc1011 | LMo-7-30-D | Lepanthes monteverdensis |
| bc1010--bc1012 | LMo-7-31-D | Lepanthes monteverdensis |
| bc1010--bc1016 | LMo-7-32-D | Lepanthes monteverdensis |
| bc1003--bc1011 | LMo-7-33/4-D | Lepanthes monteverdensis |
| bc1011--bc1040 | LMo-7-35-D | Lepanthes monteverdensis |
| bc1007--bc1011 | LMo-7-36-D | Lepanthes monteverdensis |
| bc1008--bc1011 | LMo-7-43-D | Lepanthes monteverdensis |
| bc1011--bc1050 | LMo-7-44-D | Lepanthes monteverdensis |
| bc1011--bc1011 | LMo-7-45-D | Lepanthes monteverdensis |
| bc1011--bc1057 | LMo-7-46-D | Lepanthes monteverdensis |
| bc1011--bc1058 | LMo-7-47-D | Lepanthes monteverdensis |
| bc1003--bc1012 | LMo-8-48-E | Lepanthes monteverdensis |
| bc1012--bc1040 | LFB-8-49-E | Lepanthes falx-bellica |
| bc1007--bc1012 | LMo-8-50-E | Lepanthes monteverdensis |
| bc1012--bc1049 | LMo-8-51-E | Lepanthes monteverdensis |
| bc1012--bc1050 | LFB-8-52-E | Lepanthes falx-bellica |
| bc1012--bc1052 | LMo-8-53-E | Lepanthes monteverdensis |
| bc1012--bc1012 | LMo-8-54-E | Lepanthes monteverdensis |
| bc1012--bc1058 | LFB-8-55-E | Lepanthes falx-bellica |
| bc1003--bc1015 | LFB-8-56-E | Lepanthes falx-bellica |
| bc1006--bc1015 | LFB-8-57-E | Lepanthes falx-bellica |
| bc1007--bc1015 | LFB-8-58-E | Lepanthes falx-bellica |
| bc1008--bc1015 | LMo-8-59-E | Lepanthes monteverdensis |
| bc1009--bc1015 | LMo-8-60-E | Lepanthes monteverdensis |
| bc1011--bc1015 | LMo-8-61-E | Lepanthes monteverdensis |
| bc1012--bc1015 | LMo-8-62-E | Lepanthes monteverdensis |
| bc1015--bc1016 | LMo-8-68-E | Lepanthes monteverdensis |
| bc1003--bc1016 | LFB-8-72-E | Lepanthes falx-bellica |
| bc1016--bc1040 | LC-9-74-F | Lepanthes cribbii |
| bc1007--bc1016 | LC-9-75-F | Lepanthes cribbii |
| bc1008--bc1016 | LC-9-76-F | Lepanthes cribbii |
| bc1016--bc1050 | LC-9-77-F | Lepanthes cribbii |
| bc1016--bc1052 | LC-9-78-F | Lepanthes cribbii |
| bc1016--bc1057 | LC-9-79-F | Lepanthes cribbii |
| bc1016--bc1016 | LC-9-141-F | Lepanthes cribbii |
| bc1003--bc1017 | LC-9-142-F | Lepanthes cribbii |
| bc1001--bc1018 | LC-9-143-F | Lepanthes cribbii |
| bc1001--bc1019 | LC-9-144-F | Lepanthes cribbii |
| bc1001--bc1020 | LFB-9-80-F | Lepanthes falx-bellica |
| bc1001--bc1021 | LFB-9-81-F | Lepanthes falx-bellica |
| bc1001--bc1024 | LFB-9-82-F | Lepanthes falx-bellica |

| | | |
|---|---|---|
| bc1001--bc1026 | LFB-9-83-F | Lepanthes falx-bellica |
| bc1001--bc1028 | LFB-9-84-F | Lepanthes falx-bellica |
| bc1001--bc1029 | LFB-9-145-F | Lepanthes falx-bellica |
| bc1005--bc1018 | LFB-9-146-F | Lepanthes falx-bellica |
| bc1005--bc1019 | LFB-9-147-F | Lepanthes falx-bellica |
| bc1005--bc1020 | LFB-9-148-F | Lepanthes falx-bellica |
| bc1005--bc1021 | LC-10-85-F | Lepanthes cribbii |
| bc1005--bc1024 | LC-10-87-F | Lepanthes cribbii |
| bc1005--bc1026 | LC-10-88-F | Lepanthes cribbii |
| bc1005--bc1028 | LC-10-89-F | Lepanthes cribbii |
| bc1005--bc1029 | LC-10-90-F | Lepanthes cribbii |
| bc1006--bc1060 | LC-10-149-F | Lepanthes cribbii |
| bc1006--bc1019 | LC-10-150-F | Lepanthes cribbii |
| bc1006--bc1020 | LC-10-151-F | Lepanthes cribbii |
| bc1006--bc1065 | LC-10-152-F | Lepanthes cribbii |
| bc1006--bc1066 | LFB-10-91-F | Lepanthes falx-bellica |
| bc1006--bc1067 | LFB-10-92-F | Lepanthes falx-bellica |
| bc1006--bc1028 | LFB-10-93-F | Lepanthes falx-bellica |
| bc1006--bc1073 | LFB-10-94-F | Lepanthes falx-bellica |
| bc1008--bc1060 | LFB-10-95-F | Lepanthes falx-bellica |
| bc1008--bc1019 | LFB-10-153-F | Lepanthes falx-bellica |
| bc1008--bc1020 | LFB-10-154-F | Lepanthes falx-bellica |
| bc1008--bc1065 | LFB-10-155-F | Lepanthes falx-bellica |
| bc1008--bc1066 | LFB-10-156-F | Lepanthes falx-bellica |
| bc1008--bc1067 | LFB-10-157-F | Lepanthes falx-bellica |
| bc1008--bc1028 | LC-11-96-F | Lepanthes cribbii |
| bc1008--bc1073 | LC-11-97-F | Lepanthes cribbii |
| bc1009--bc1060 | LC-11-98-F | Lepanthes cribbii |
| bc1009--bc1019 | LC-11-99-F | Lepanthes cribbii |
| bc1009--bc1020 | LC-11-100-F | Lepanthes cribbii |
| bc1009--bc1065 | LC-11-158-F | Lepanthes cribbii |
| bc1009--bc1066 | LFB-11-101-F | Lepanthes falx-bellica |
| bc1009--bc1067 | LFB-11-102-F | Lepanthes falx-bellica |
| bc1009--bc1028 | LFB-11-103-F | Lepanthes falx-bellica |
| bc1009--bc1073 | LFB-11-104-F | Lepanthes falx-bellica |
| bc1010--bc1018 | LFB-11-105-F | Lepanthes falx-bellica |
| bc1010--bc1019 | LFB-11-159-F | Lepanthes falx-bellica |
| bc1010--bc1020 | LFB-11-160-F | Lepanthes falx-bellica |
| bc1010--bc1021 | LFB-11-161-F | Lepanthes falx-bellica |
| bc1010--bc1024 | LFB-11-162-F | Lepanthes falx-bellica |
| bc1010--bc1026 | LFB-11-163-F | Lepanthes falx-bellica |
| bc1010--bc1028 | LFB-12-106-F | Lepanthes falx-bellica |
| bc1010--bc1029 | LFB-12-107-F | Lepanthes falx-bellica |
| bc1011--bc1060 | LFB-12-108-F | Lepanthes falx-bellica |

| | | |
|---|---|---|
| bc1011--bc1019 | LFB-12-109-F | Lepanthes falx-bellica |
| bc1011--bc1020 | LFB-12-110-F | Lepanthes falx-bellica |
| bc1011--bc1065 | LFB-12-111-F | Lepanthes falx-bellica |
| bc1011--bc1066 | LFB-12-112-F | Lepanthes falx-bellica |
| bc1011--bc1067 | LFB-12-114-F | Lepanthes falx-bellica |
| bc1011--bc1028 | LFB-12-115-F | Lepanthes falx-bellica |
| bc1011--bc1073 | LC-12-116-F | Lepanthes cribbii |
| bc1012--bc1060 | LC-12-117-F | Lepanthes cribbii |
| bc1012--bc1019 | LC-12-118-F | Lepanthes cribbii |
| bc1012--bc1020 | LC-12-119-F | Lepanthes cribbii |
| bc1012--bc1065 | LC-12-120-F | Lepanthes cribbii |
| bc1012--bc1066 | LC-12-121-F | Lepanthes cribbii |
| bc1012--bc1067 | LC-12-122-F | Lepanthes cribbii |
| bc1012--bc1028 | LC-12-123-F | Lepanthes cribbii |
| bc1012--bc1073 | LC-12-124-F | Lepanthes cribbii |
| bc1015--bc1018 | LC-12-125-F | Lepanthes cribbii |
| bc1015--bc1019 | LMo-12-126-F | Lepanthes monteverdensis |
| bc1015--bc1020 | LMo-12-127-F | Lepanthes monteverdensis |
| bc1015--bc1021 | LMo-12-128-F | Lepanthes monteverdensis |
| bc1015--bc1024 | LMo-12-129-F | Lepanthes monteverdensis |
| bc1015--bc1026 | LMo-12-130-F | Lepanthes monteverdensis |
| bc1015--bc1028 | LMo-12-131-F | Lepanthes monteverdensis |
| bc1015--bc1029 | LMo-12-132-F | Lepanthes monteverdensis |
| bc1016--bc1060 | LMo-12-133-F | Lepanthes monteverdensis |
| bc1016--bc1019 | LMo-12-134-F | Lepanthes monteverdensis |
| bc1016--bc1020 | LC-13-126-G | Lepanthes cribbii |
| bc1016--bc1065 | LC-13-127-G | Lepanthes cribbii |
| bc1016--bc1066 | LC-13-128-G | Lepanthes cribbii |
| bc1016--bc1067 | LC-13-129-G | Lepanthes cribbii |
| bc1016--bc1028 | LC-13-130-G | Lepanthes cribbii |
| bc1001--bc1031 | LMe-16-187-J | Lepanthes mentosa |
| bc1001--bc1032 | LMe-16-188-J | Lepanthes mentosa |
| bc1001--bc1033 | LMe-16-189-J | Lepanthes mentosa |
| bc1001--bc1034 | LMe-16-190-J | Lepanthes mentosa |
| bc1001--bc1038 | LMe-16-191-J | Lepanthes mentosa |
| bc1001--bc1044 | LMe-17-193-J | Lepanthes mentosa |
| bc1001--bc1045 | LMe-17-194-J | Lepanthes mentosa |
| bc1005--bc1031 | LMe-17-195-J | Lepanthes mentosa |
| bc1005--bc1032 | LMe-17-196-J | Lepanthes mentosa |
| bc1005--bc1033 | LMe-17-197-J | Lepanthes mentosa |
| bc1005--bc1034 | LMe-17-198-J | Lepanthes mentosa |
| bc1005--bc1038 | LC-17-199-J | Lepanthes cribbii |
| bc1005--bc1043 | LC-17-200-J | Lepanthes cribbii |
| bc1005--bc1044 | LC-17-201-J | Lepanthes cribbii |

| | | |
|---|---|---|
| bc1005--bc1045 | LC-17-202-J | Lepanthes cribbii |
| bc1006--bc1075 | LC-17-203-J | Lepanthes cribbii |
| bc1006--bc1032 | LC-17-204-J | Lepanthes cribbii |
| bc1006--bc1081 | LC-17-205-J | Lepanthes cribbii |
| bc1006--bc1084 | LC-17-206-J | Lepanthes cribbii |
| bc1006--bc1038 | LC-17-207-J | Lepanthes cribbii |
| bc1006--bc1043 | LC-17-208-J | Lepanthes cribbii |
| bc1006--bc1044 | LFB-19-214-K | Lepanthes falx-bellica |
| bc1006--bc1093 | LFB-19-215-K | Lepanthes falx-bellica |
| bc1008--bc1075 | LFB-19-216-K | Lepanthes falx-bellica |
| bc1008--bc1032 | LFB-19-217-K | Lepanthes falx-bellica |
| bc1008--bc1081 | LFB-19-218-K | Lepanthes falx-bellica |
| bc1008--bc1084 | LFB-19-219-K | Lepanthes falx-bellica |
| bc1008--bc1038 | LFB-19-220-K | Lepanthes falx-bellica |
| bc1008--bc1043 | LFB-19-221-K | Lepanthes falx-bellica |
| bc1008--bc1044 | LC-19-224-K | Lepanthes cribbii |
| bc1008--bc1093 | LC-19-225-K | Lepanthes cribbii |
| bc1009--bc1075 | LC-19-226-K | Lepanthes cribbii |
| bc1009--bc1032 | LC-19-227-K | Lepanthes cribbii |
| bc1009--bc1081 | LC-19-228-K | Lepanthes cribbii |
| bc1009--bc1084 | LC-19-229-K | Lepanthes cribbii |
| bc1009--bc1038 | LC-19-230-K | Lepanthes cribbii |
| bc1009--bc1043 | LC-19-231-K | Lepanthes cribbii |
| bc1009--bc1044 | LC-19-232-K | Lepanthes cribbii |
| bc1009--bc1093 | LC-19-233-K | Lepanthes cribbii |
| bc1010--bc1031 | LFB-19-234-K | Lepanthes falx-bellica |
| bc1010--bc1032 | LFB-19-235-K | Lepanthes falx-bellica |
| bc1010--bc1033 | LFB-20-237-K | Lepanthes falx-bellica |
| bc1010--bc1034 | LFB-20-238-K | Lepanthes falx-bellica |
| bc1010--bc1038 | LFB-20-239-K | Lepanthes falx-bellica |
| bc1010--bc1043 | LFB-20-240-K | Lepanthes falx-bellica |
| bc1010--bc1044 | LFB-20-241-K | Lepanthes falx-bellica |
| bc1010--bc1045 | LFB-20-242-K | Lepanthes falx-bellica |
| bc1011--bc1075 | LFB-20-243-K | Lepanthes falx-bellica |
| bc1011--bc1032 | LFB-20-244-K | Lepanthes falx-bellica |
| bc1011--bc1081 | LFB-21-245-K | Lepanthes falx-bellica |
| bc1011--bc1084 | LFB-21-246-K | Lepanthes falx-bellica |
| bc1011--bc1038 | LFB-21-247-K | Lepanthes falx-bellica |
| bc1011--bc1043 | LFB-21-248-K | Lepanthes falx-bellica |
| bc1011--bc1044 | LFB-21-249-K | Lepanthes falx-bellica |
| bc1011--bc1093 | LFB-21-250-K | Lepanthes falx-bellica |
| bc1012--bc1075 | LFB-21-251-K | Lepanthes falx-bellica |
| bc1012--bc1032 | LFB-21-252-K | Lepanthes falx-bellica |
| bc1012--bc1081 | LFB-21-253-K | Lepanthes falx-bellica |

| | | |
|---|---|---|
| bc1012--bc1084 | LFB-21-254-K | Lepanthes falx-bellica |
| bc1012--bc1038 | LC-21-255-K | Lepanthes cribbii |
| bc1012--bc1043 | LC-21-256-K | Lepanthes cribbii |
| bc1012--bc1044 | LC-21-257-K | Lepanthes cribbii |
| bc1012--bc1093 | LC-21-258-K | Lepanthes cribbii |
| bc1015--bc1031 | LC-21-259-K | Lepanthes cribbii |
| bc1015--bc1032 | LC-21-260-K | Lepanthes cribbii |
| bc1015--bc1033 | LC-21-261-K | Lepanthes cribbii |
| bc1015--bc1034 | LC-21-262-K | Lepanthes cribbii |
| bc1015--bc1038 | LC-21-263-K | Lepanthes cribbii |
| bc1015--bc1043 | LC-21-264-K | Lepanthes cribbii |
| bc1015--bc1044 | LFB-22-265-K | Lepanthes falx-bellica |
| bc1015--bc1045 | LFB-22-266-K | Lepanthes falx-bellica |
| bc1016--bc1075 | LFB-22-267-K | Lepanthes falx-bellica |
| bc1016--bc1032 | LFB-22-268-K | Lepanthes falx-bellica |
| bc1016--bc1081 | LFB-22-269-K | Lepanthes falx-bellica |
| bc1016--bc1084 | LFB-22-270-K | Lepanthes falx-bellica |
| bc1016--bc1038 | LFB-22-271-K | Lepanthes falx-bellica |
| bc1016--bc1043 | LFB-23-272-K | Lepanthes falx-bellica |
| bc1016--bc1044 | LFB-23-273-K | Lepanthes falx-bellica |
| bc1016--bc1093 | LFB-23-274-K | Lepanthes falx-bellica |
| bc1017--bc1031 | LFB-23-275-K | Lepanthes falx-bellica |
| bc1017--bc1032 | LFB-23-276-K | Lepanthes falx-bellica |
| bc1017--bc1033 | LFB-23-277-K | Lepanthes falx-bellica |
| bc1017--bc1034 | LFB-23-278-K | Lepanthes falx-bellica |
| bc1017--bc1038 | LFB-23-279-K | Lepanthes falx-bellica |
| bc1017--bc1043 | LFB-24-280-K | Lepanthes falx-bellica |
| bc1017--bc1044 | LFB-24-281-K | Lepanthes falx-bellica |
| bc1017--bc1045 | LFB-24-282-K | Lepanthes falx-bellica |
| bc1018--bc1075 | LFB-24-283-K | Lepanthes falx-bellica |
| bc1018--bc1032 | LFB-24-284-K | Lepanthes falx-bellica |
| bc1018--bc1081 | LFB-24-285-K | Lepanthes falx-bellica |
| bc1018--bc1084 | LFB-24-286-K | Lepanthes falx-bellica |
| bc1018--bc1038 | LFB-25-287-K | Lepanthes falx-bellica |
| bc1018--bc1043 | LFB-25-288-K | Lepanthes falx-bellica |
| bc1018--bc1044 | LFB-25-289-K | Lepanthes falx-bellica |
| bc1018--bc1093 | LFB-25-290-K | Lepanthes falx-bellica |
| bc1003--bc1021 | LFB-25-291-K | Lepanthes falx-bellica |
| bc1021--bc1040 | LFB-25-292-K | Lepanthes falx-bellica |
| bc1007--bc1021 | LFB-25-293-K | Lepanthes falx-bellica |
| bc1021--bc1049 | LFB-25-294-K | Lepanthes falx-bellica |
| bc1021--bc1050 | LFB-25-295-K | Lepanthes falx-bellica |
| bc1021--bc1052 | LFB-26-296-L | Lepanthes falx-bellica |
| bc1021--bc1057 | LFB-26-297-L | Lepanthes falx-bellica |

| | | |
|---|---|---|
| bc1021--bc1058 | LFB-26-298-L | Lepanthes falx-bellica |
| bc1003--bc1022 | LFB-26-299-L | Lepanthes falx-bellica |
| bc1006--bc1022 | LFB-26-300-L | Lepanthes falx-bellica |
| bc1007--bc1022 | LFB-26-301-L | Lepanthes falx-bellica |
| bc1008--bc1022 | LFB-26-302-L | Lepanthes falx-bellica |
| bc1009--bc1022 | LFB-26-303-L | Lepanthes falx-bellica |
| bc1011--bc1022 | LFB-26-304-L | Lepanthes falx-bellica |
| bc1012--bc1022 | LFB-26-305-L | Lepanthes falx-bellica |
| bc1016--bc1022 | LMe-27-306-L | Lepanthes mentosa |
| bc1003--bc1024 | LMe-27-307-L | Lepanthes mentosa |
| bc1024--bc1040 | LMe-27-308-L | Lepanthes mentosa |
| bc1007--bc1024 | LMe-27-309-L | Lepanthes mentosa |
| bc1024--bc1049 | LMe-27-310-L | Lepanthes mentosa |
| bc1024--bc1050 | LMe-27-311-L | Lepanthes mentosa |
| bc1024--bc1052 | LMe-27-312-L | Lepanthes mentosa |
| bc1024--bc1057 | LMe-27-313-L | Lepanthes mentosa |
| bc1024--bc1058 | LFB-28-314-M | Lepanthes falx-bellica |
| bc1003--bc1026 | LFB-28-315-M | Lepanthes falx-bellica |
| bc1026--bc1040 | LFB-28-316-M | Lepanthes falx-bellica |
| bc1007--bc1026 | LFB-28-317-M | Lepanthes falx-bellica |
| bc1026--bc1049 | LFB-28-318-M | Lepanthes falx-bellica |
| bc1026--bc1050 | LFB-28-319-M | Lepanthes falx-bellica |
| bc1026--bc1052 | LFB-28-320-M | Lepanthes falx-bellica |
| bc1026--bc1057 | LFB-28-321-M | Lepanthes falx-bellica |
| bc1026--bc1058 | LFB-28-322-M | Lepanthes falx-bellica |
| bc1003--bc1029 | LFB-28-323-M | Lepanthes falx-bellica |
| bc1029--bc1040 | LFB-29-324-M | Lepanthes falx-bellica |
| bc1007--bc1029 | LFB-29-325-M | Lepanthes falx-bellica |
| bc1029--bc1049 | LFB-29-326-M | Lepanthes falx-bellica |
| bc1029--bc1050 | LFB-29-327-M | Lepanthes falx-bellica |
| bc1029--bc1052 | LFB-29-328-M | Lepanthes falx-bellica |
| bc1029--bc1057 | LFB-29-329-M | Lepanthes falx-bellica |
| bc1016--bc1029 | LFB-29-330-M | Lepanthes falx-bellica |
| bc1003--bc1030 | LFB-29-331-M | Lepanthes falx-bellica |
| bc1006--bc1030 | LFB-29-332-M | Lepanthes falx-bellica |
| bc1007--bc1030 | LFB-29-333-M | Lepanthes falx-bellica |
| bc1008--bc1030 | LC-30-334-M | Lepanthes cribbii |
| bc1009--bc1030 | LC-30-335-M | Lepanthes cribbii |
| bc1011--bc1030 | LC-30-336-M | Lepanthes cribbii |
| bc1012--bc1030 | LFB-30-337-M | Lepanthes falx-bellica |
| bc1016--bc1030 | LFB-30-338-M | Lepanthes falx-bellica |
| bc1003--bc1031 | LFB-30-339-M | Lepanthes falx-bellica |
| bc1031--bc1040 | LFB-30-340-M | Lepanthes falx-bellica |
| bc1007--bc1031 | LFB-30-341-M | Lepanthes falx-bellica |

| | | |
|---|---|---|
| bc1031--bc1049 | LFB-30-342-M | Lepanthes falx-bellica |
| bc1031--bc1050 | LFB-30-343-M | Lepanthes falx-bellica |
| bc1031--bc1052 | LFB-30-344-M | Lepanthes falx-bellica |
| bc1031--bc1057 | LFB-31-348-M | Lepanthes falx-bellica |
| bc1031--bc1058 | LFB-31-349-M | Lepanthes falx-bellica |
| bc1003--bc1033 | LFB-31-350-M | Lepanthes falx-bellica |
| bc1033--bc1040 | LFB-31-351-M | Lepanthes falx-bellica |
| bc1007--bc1033 | LFB-31-352-M | Lepanthes falx-bellica |
| bc1033--bc1049 | LMe-32-353-L | Lepanthes mentosa |
| bc1033--bc1050 | LMe-32-354-L | Lepanthes mentosa |
| bc1033--bc1052 | LMe-32-355-L | Lepanthes mentosa |
| bc1033--bc1057 | LMe-32-356-L | Lepanthes mentosa |
| bc1033--bc1058 | LMe-32-357-L | Lepanthes mentosa |
| bc1003--bc1034 | LMe-32-358-L | Lepanthes mentosa |
| bc1034--bc1040 | LMe-32-359-L | Lepanthes mentosa |
| bc1007--bc1034 | LMe-32-360-L | Lepanthes mentosa |
| bc1034--bc1049 | LMe-32-361-L | Lepanthes mentosa |
| bc1034--bc1050 | LMo-33-367-L | Lepanthes monteverdensis |
| bc1034--bc1052 | LMo-33-368-L | Lepanthes monteverdensis |
| bc1034--bc1057 | LMo-33-369-L | Lepanthes monteverdensis |
| bc1034--bc1058 | LMo-33-370-L | Lepanthes monteverdensis |
| bc1003--bc1036 | LMo-33-371-L | Lepanthes monteverdensis |
| bc1006--bc1036 | LMo-33-372-L | Lepanthes monteverdensis |
| bc1007--bc1036 | LMo-33-373-L | Lepanthes monteverdensis |
| bc1008--bc1036 | LMo-33-374-L | Lepanthes monteverdensis |
| bc1009--bc1036 | LMo-33-375-L | Lepanthes monteverdensis |
| bc1011--bc1036 | LMo-33-376-L | Lepanthes monteverdensis |
| bc1012--bc1036 | LFB-34-377-N | Lepanthes falx-bellica |
| bc1016--bc1036 | LFB-34-378-N | Lepanthes falx-bellica |
| bc1003--bc1037 | LFB-34-379-N | Lepanthes falx-bellica |
| bc1006--bc1037 | LFB-34-380-N | Lepanthes falx-bellica |
| bc1007--bc1037 | LFB-34-381-N | Lepanthes falx-bellica |
| bc1008--bc1037 | LFB-34-382-N | Lepanthes falx-bellica |
| bc1009--bc1037 | LFB-34-383-N | Lepanthes falx-bellica |
| bc1011--bc1037 | LFB-34-384-N | Lepanthes falx-bellica |
| bc1012--bc1037 | LFB-35-385-N | Lepanthes falx-bellica |
| bc1016--bc1037 | LFB-35-386-N | Lepanthes falx-bellica |
| bc1003--bc1039 | LFB-35-387-N | Lepanthes falx-bellica |
| bc1006--bc1039 | LFB-35-388-N | Lepanthes falx-bellica |
| bc1007--bc1039 | LFB-35-389-N | Lepanthes falx-bellica |
| bc1008--bc1039 | LFB-35-390-N | Lepanthes falx-bellica |
| bc1009--bc1039 | LFB-35-391-N | Lepanthes falx-bellica |
| bc1011--bc1039 | LFB-35-392-N | Lepanthes falx-bellica |
| bc1012--bc1039 | LMo-36-393-N | Lepanthes monteverdensis |

| | | |
|---|---|---|
| bc1016--bc1039 | LMo-36-394-N | Lepanthes monteverdensis |
| bc1018--bc1021 | LMo-36-395-N | Lepanthes monteverdensis |
| bc1019--bc1021 | LMo-36-396-N | Lepanthes monteverdensis |
| bc1020--bc1021 | LMo-36-397-N | Lepanthes monteverdensis |
| bc1021--bc1021 | LMo-36-398-N | Lepanthes monteverdensis |
| bc1021--bc1066 | LMo-36-399-N | Lepanthes monteverdensis |
| bc1021--bc1067 | LMo-36-400-N | Lepanthes monteverdensis |
| bc1021--bc1028 | LMo-36-401-N | Lepanthes monteverdensis |
| bc1021--bc1073 | LMo-36-402-N | Lepanthes monteverdensis |
| bc1018--bc1022 | LMe-37-403-L | Lepanthes mentosa |
| bc1019--bc1022 | LMe-37-404-L | Lepanthes mentosa |
| bc1020--bc1022 | LMe-37-405-L | Lepanthes mentosa |
| bc1021--bc1022 | LMe-37-406-L | Lepanthes mentosa |
| bc1022--bc1024 | LMe-37-407-L | Lepanthes mentosa |
| bc1022--bc1026 | LMe-37-408-L | Lepanthes mentosa |
| bc1022--bc1028 | LMe-37-409-L | Lepanthes mentosa |
| bc1022--bc1029 | LMe-37-410-L | Lepanthes mentosa |
| bc1018--bc1024 | LMe-37-411-L | Lepanthes mentosa |
| bc1019--bc1024 | LMe-37-412-L | Lepanthes mentosa |
| bc1020--bc1024 | LMo-38-418-L | Lepanthes monteverdensis |
| bc1024--bc1065 | LMo-38-419-L | Lepanthes monteverdensis |
| bc1024--bc1024 | LMo-38-420-L | Lepanthes monteverdensis |
| bc1024--bc1067 | LMo-38-421-L | Lepanthes monteverdensis |
| bc1024--bc1028 | LC-38-422-L | Lepanthes cribbii |
| bc1024--bc1073 | LFB-38-423-L | Lepanthes falx-bellica |
| bc1018--bc1026 | LFB-38-424-L | Lepanthes falx-bellica |
| bc1019--bc1026 | LFB-38-425-L | Lepanthes falx-bellica |
| bc1020--bc1026 | LFB-38-426-L | Lepanthes falx-bellica |
| bc1026--bc1065 | LFB-38-427-L | Lepanthes falx-bellica |
| bc1026--bc1066 | LMo-39-428-L | Lepanthes monteverdensis |
| bc1026--bc1026 | LMo-39-429-L | Lepanthes monteverdensis |
| bc1026--bc1028 | LMo-39-430-L | Lepanthes monteverdensis |
| bc1026--bc1073 | LMo-39-431-L | Lepanthes monteverdensis |
| bc1018--bc1029 | LMo-39-432-L | Lepanthes monteverdensis |
| bc1019--bc1029 | LMo-39-433-L | Lepanthes monteverdensis |
| bc1020--bc1029 | LMo-39-434-L | Lepanthes monteverdensis |
| bc1029--bc1065 | LMo-39-435-L | Lepanthes monteverdensis |
| bc1029--bc1066 | LMo-39-436-L | Lepanthes monteverdensis |
| bc1029--bc1067 | LMo-39-437-L | Lepanthes monteverdensis |
| bc1028--bc1029 | LMo-40-438-L | Lepanthes monteverdensis |
| bc1029--bc1029 | LMo-40-439-L | Lepanthes monteverdensis |
| bc1018--bc1030 | LMo-40-440-L | Lepanthes monteverdensis |
| bc1019--bc1030 | LMo-40-441-L | Lepanthes monteverdensis |
| bc1020--bc1030 | LMo-40-442-L | Lepanthes monteverdensis |

| | | |
|---|---|---|
| bc1021--bc1030 | LMo-40-443-L | Lepanthes monteverdensis |
| bc1024--bc1030 | LMo-40-444-L | Lepanthes monteverdensis |
| bc1026--bc1030 | LMo-40-445-L | Lepanthes monteverdensis |
| bc1029--bc1030 | LMe-41-447-O | Lepanthes mentosa |
| bc1031--bc1060 | LMe-41-448-O | Lepanthes mentosa |
| bc1019--bc1031 | LMe-41-449-O | Lepanthes mentosa |
| bc1020--bc1031 | LMe-41-450-O | Lepanthes mentosa |
| bc1021--bc1031 | LMe-41-451-O | Lepanthes mentosa |
| bc1024--bc1031 | LMe-41-452-O | Lepanthes mentosa |
| bc1026--bc1031 | LMe-41-453-O | Lepanthes mentosa |
| bc1028--bc1031 | LMe-41-454-O | Lepanthes mentosa |
| bc1029--bc1031 | LMe-41-455-O | Lepanthes mentosa |
| bc1033--bc1060 | LMe-41-456-O | Lepanthes mentosa |
| bc1019--bc1033 | LMe-38-457-L | Lepanthes mentosa |
| bc1020--bc1033 | LMe-38-458-L | Lepanthes mentosa |
| bc1115--bc1191 | SynMock | SynMock |

| | |
|---|---|
| orchid population | tree sp. |
| categorical | categorical |
| Lepanthes monteverdensis-3-B | Pouteria exfoliata, cf. (Sapotaceae) |
| Lepanthes monteverdensis-3-B | Pouteria exfoliata, cf. (Sapotaceae) |
| Lepanthes monteverdensis-3-B | Pouteria exfoliata, cf. (Sapotaceae) |
| Lepanthes monteverdensis-3-B | Pouteria exfoliata, cf. (Sapotaceae) |
| Lepanthes monteverdensis-3-B | Pouteria exfoliata, cf. (Sapotaceae) |
| Lepanthes monteverdensis-3-B | Pouteria exfoliata, cf. (Sapotaceae) |
| Lepanthes monteverdensis-3-B | Pouteria exfoliata, cf. (Sapotaceae) |
| Lepanthes monteverdensis-3-B | Pouteria exfoliata, cf. (Sapotaceae) |
| Lepanthes monteverdensis-3-B | Pouteria exfoliata, cf. (Sapotaceae) |
| Lepanthes monteverdensis-3-B | Pouteria exfoliata, cf. (Sapotaceae) |
| Lepanthes monteverdensis-5-B | Quercus brenesii (=Q.cortesii) (Fagaceae) |
| Lepanthes monteverdensis-5-B | Quercus brenesii (=Q.cortesii) (Fagaceae) |
| Lepanthes monteverdensis-5-B | Quercus brenesii (=Q.cortesii) (Fagaceae) |
| Lepanthes monteverdensis-5-B | Quercus brenesii (=Q.cortesii) (Fagaceae) |
| Lepanthes monteverdensis-5-B | Quercus brenesii (=Q.cortesii) (Fagaceae) |
| Lepanthes monteverdensis-5-B | Quercus brenesii (=Q.cortesii) (Fagaceae) |
| Lepanthes monteverdensis-5-B | Quercus brenesii (=Q.cortesii) (Fagaceae) |
| Lepanthes monteverdensis-5-B | Quercus brenesii (=Q.cortesii) (Fagaceae) |
| Lepanthes monteverdensis-5-B | Quercus brenesii (=Q.cortesii) (Fagaceae) |
| Lepanthes monteverdensis-5-B | Quercus brenesii (=Q.cortesii) (Fagaceae) |
| Lepanthes falx-bellica-5-B | Quercus brenesii (=Q.cortesii) (Fagaceae) |
| Lepanthes falx-bellica-5-B | Quercus brenesii (=Q.cortesii) (Fagaceae) |
| Lepanthes falx-bellica-5-B | Quercus brenesii (=Q.cortesii) (Fagaceae) |
| Lepanthes falx-bellica-5-B | Quercus brenesii (=Q.cortesii) (Fagaceae) |
| Lepanthes falx-bellica-5-B | Quercus brenesii (=Q.cortesii) (Fagaceae) |
| Lepanthes falx-bellica-5-B | Quercus brenesii (=Q.cortesii) (Fagaceae) |
| Lepanthes falx-bellica-5-B | Quercus brenesii (=Q.cortesii) (Fagaceae) |
| Lepanthes falx-bellica-5-B | Quercus brenesii (=Q.cortesii) (Fagaceae) |
| Lepanthes falx-bellica-5-B | Quercus brenesii (=Q.cortesii) (Fagaceae) |
| Lepanthes falx-bellica-5-B | Quercus brenesii (=Q.cortesii) (Fagaceae) |
| Lepanthes falx-bellica-6-C | Miconia oerstediana (Melastomataceae) |
| Lepanthes falx-bellica-6-C | Miconia oerstediana (Melastomataceae) |
| Lepanthes falx-bellica-6-C | Miconia oerstediana (Melastomataceae) |
| Lepanthes falx-bellica-6-C | Miconia oerstediana (Melastomataceae) |
| Lepanthes monteverdensis-6-C | Miconia oerstediana (Melastomataceae) |
| Lepanthes falx-bellica-6-C | Miconia oerstediana (Melastomataceae) |
| Lepanthes monteverdensis-6-C | Miconia oerstediana (Melastomataceae) |
| Lepanthes monteverdensis-6-C | Miconia oerstediana (Melastomataceae) |
| Lepanthes monteverdensis-6-C | Miconia oerstediana (Melastomataceae) |
| Lepanthes monteverdensis-6-C | Miconia oerstediana (Melastomataceae) |
| Lepanthes monteverdensis-6-C | Miconia oerstediana (Melastomataceae) |
| Lepanthes monteverdensis-6-C | Miconia oerstediana (Melastomataceae) |

| | |
|---|---|
| Lepanthes monteverdensis-6-C | Miconia oerstediana (Melastomataceae) |
| Lepanthes monteverdensis-6-C | Miconia oerstediana (Melastomataceae) |
| Lepanthes monteverdensis-6-C | Miconia oerstediana (Melastomataceae) |
| Lepanthes monteverdensis-7-D | Eugenia valerii (Myrtaceae) |
| Lepanthes monteverdensis-7-D | Eugenia valerii (Myrtaceae) |
| Lepanthes monteverdensis-7-D | Eugenia valerii (Myrtaceae) |
| Lepanthes monteverdensis-7-D | Eugenia valerii (Myrtaceae) |
| Lepanthes monteverdensis-7-D | Eugenia valerii (Myrtaceae) |
| Lepanthes monteverdensis-7-D | Eugenia valerii (Myrtaceae) |
| Lepanthes monteverdensis-7-D | Eugenia valerii (Myrtaceae) |
| Lepanthes monteverdensis-7-D | Eugenia valerii (Myrtaceae) |
| Lepanthes monteverdensis-7-D | Eugenia valerii (Myrtaceae) |
| Lepanthes monteverdensis-7-D | Eugenia valerii (Myrtaceae) |
| Lepanthes monteverdensis-7-D | Eugenia valerii (Myrtaceae) |
| Lepanthes monteverdensis-8-E | Calyptranthes monteverdensis (Myrtaceae) |
| Lepanthes falx-bellica-8-E | Calyptranthes monteverdensis (Myrtaceae) |
| Lepanthes monteverdensis-8-E | Calyptranthes monteverdensis (Myrtaceae) |
| Lepanthes monteverdensis-8-E | Calyptranthes monteverdensis (Myrtaceae) |
| Lepanthes falx-bellica-8-E | Calyptranthes monteverdensis (Myrtaceae) |
| Lepanthes monteverdensis-8-E | Calyptranthes monteverdensis (Myrtaceae) |
| Lepanthes monteverdensis-8-E | Calyptranthes monteverdensis (Myrtaceae) |
| Lepanthes falx-bellica-8-E | Calyptranthes monteverdensis (Myrtaceae) |
| Lepanthes falx-bellica-8-E | Calyptranthes monteverdensis (Myrtaceae) |
| Lepanthes falx-bellica-8-E | Calyptranthes monteverdensis (Myrtaceae) |
| Lepanthes falx-bellica-8-E | Calyptranthes monteverdensis (Myrtaceae) |
| Lepanthes monteverdensis-8-E | Calyptranthes monteverdensis (Myrtaceae) |
| Lepanthes monteverdensis-8-E | Calyptranthes monteverdensis (Myrtaceae) |
| Lepanthes monteverdensis-8-E | Calyptranthes monteverdensis (Myrtaceae) |
| Lepanthes monteverdensis-8-E | Calyptranthes monteverdensis (Myrtaceae) |
| Lepanthes monteverdensis-8-E | Calyptranthes monteverdensis (Myrtaceae) |
| Lepanthes falx-bellica-8-E | Calyptranthes monteverdensis (Myrtaceae) |
| Lepanthes cribbii-9-F | Calyptranthes monteverdensis (Myrtaceae) |
| Lepanthes cribbii-9-F | Calyptranthes monteverdensis (Myrtaceae) |
| Lepanthes cribbii-9-F | Calyptranthes monteverdensis (Myrtaceae) |
| Lepanthes cribbii-9-F | Calyptranthes monteverdensis (Myrtaceae) |
| Lepanthes cribbii-9-F | Calyptranthes monteverdensis (Myrtaceae) |
| Lepanthes cribbii-9-F | Calyptranthes monteverdensis (Myrtaceae) |
| Lepanthes cribbii-9-F | Calyptranthes monteverdensis (Myrtaceae) |
| Lepanthes cribbii-9-F | Calyptranthes monteverdensis (Myrtaceae) |
| Lepanthes cribbii-9-F | Calyptranthes monteverdensis (Myrtaceae) |
| Lepanthes cribbii-9-F | Calyptranthes monteverdensis (Myrtaceae) |
| Lepanthes falx-bellica-9-F | Calyptranthes monteverdensis (Myrtaceae) |
| Lepanthes falx-bellica-9-F | Calyptranthes monteverdensis (Myrtaceae) |
| Lepanthes falx-bellica-9-F | Calyptranthes monteverdensis (Myrtaceae) |

| | |
|---|---|
| Lepanthes falx-bellica-9-F | Calyptranthes monteverdensis (Myrtaceae) |
| Lepanthes falx-bellica-9-F | Calyptranthes monteverdensis (Myrtaceae) |
| Lepanthes falx-bellica-9-F | Calyptranthes monteverdensis (Myrtaceae) |
| Lepanthes falx-bellica-9-F | Calyptranthes monteverdensis (Myrtaceae) |
| Lepanthes falx-bellica-9-F | Calyptranthes monteverdensis (Myrtaceae) |
| Lepanthes falx-bellica-9-F | Calyptranthes monteverdensis (Myrtaceae) |
| Lepanthes cribbii-10-F | Calyptranthes monteverdensis (Myrtaceae) |
| Lepanthes cribbii-10-F | Calyptranthes monteverdensis (Myrtaceae) |
| Lepanthes cribbii-10-F | Calyptranthes monteverdensis (Myrtaceae) |
| Lepanthes cribbii-10-F | Calyptranthes monteverdensis (Myrtaceae) |
| Lepanthes cribbii-10-F | Calyptranthes monteverdensis (Myrtaceae) |
| Lepanthes cribbii-10-F | Calyptranthes monteverdensis (Myrtaceae) |
| Lepanthes cribbii-10-F | Calyptranthes monteverdensis (Myrtaceae) |
| Lepanthes cribbii-10-F | Calyptranthes monteverdensis (Myrtaceae) |
| Lepanthes cribbii-10-F | Calyptranthes monteverdensis (Myrtaceae) |
| Lepanthes falx-bellica-10-F | Calyptranthes monteverdensis (Myrtaceae) |
| Lepanthes falx-bellica-10-F | Calyptranthes monteverdensis (Myrtaceae) |
| Lepanthes falx-bellica-10-F | Calyptranthes monteverdensis (Myrtaceae) |
| Lepanthes falx-bellica-10-F | Calyptranthes monteverdensis (Myrtaceae) |
| Lepanthes falx-bellica-10-F | Calyptranthes monteverdensis (Myrtaceae) |
| Lepanthes falx-bellica-10-F | Calyptranthes monteverdensis (Myrtaceae) |
| Lepanthes falx-bellica-10-F | Calyptranthes monteverdensis (Myrtaceae) |
| Lepanthes falx-bellica-10-F | Calyptranthes monteverdensis (Myrtaceae) |
| Lepanthes falx-bellica-10-F | Calyptranthes monteverdensis (Myrtaceae) |
| Lepanthes falx-bellica-10-F | Calyptranthes monteverdensis (Myrtaceae) |
| Lepanthes cribbii-11-F | Calyptranthes monteverdensis (Myrtaceae) |
| Lepanthes cribbii-11-F | Calyptranthes monteverdensis (Myrtaceae) |
| Lepanthes cribbii-11-F | Calyptranthes monteverdensis (Myrtaceae) |
| Lepanthes cribbii-11-F | Calyptranthes monteverdensis (Myrtaceae) |
| Lepanthes cribbii-11-F | Calyptranthes monteverdensis (Myrtaceae) |
| Lepanthes cribbii-11-F | Calyptranthes monteverdensis (Myrtaceae) |
| Lepanthes falx-bellica-11-F | Calyptranthes monteverdensis (Myrtaceae) |
| Lepanthes falx-bellica-11-F | Calyptranthes monteverdensis (Myrtaceae) |
| Lepanthes falx-bellica-11-F | Calyptranthes monteverdensis (Myrtaceae) |
| Lepanthes falx-bellica-11-F | Calyptranthes monteverdensis (Myrtaceae) |
| Lepanthes falx-bellica-11-F | Calyptranthes monteverdensis (Myrtaceae) |
| Lepanthes falx-bellica-11-F | Calyptranthes monteverdensis (Myrtaceae) |
| Lepanthes falx-bellica-11-F | Calyptranthes monteverdensis (Myrtaceae) |
| Lepanthes falx-bellica-11-F | Calyptranthes monteverdensis (Myrtaceae) |
| Lepanthes falx-bellica-11-F | Calyptranthes monteverdensis (Myrtaceae) |
| Lepanthes falx-bellica-11-F | Calyptranthes monteverdensis (Myrtaceae) |
| Lepanthes falx-bellica-12-F | Rondeletia buddleioides (Rubiaceae) |
| Lepanthes falx-bellica-12-F | Rondeletia buddleioides (Rubiaceae) |
| Lepanthes falx-bellica-12-F | Rondeletia buddleioides (Rubiaceae) |

| | |
|---|---|
| Lepanthes falx-bellica-12-F | Rondeletia buddleioides (Rubiaceae) |
| Lepanthes falx-bellica-12-F | Rondeletia buddleioides (Rubiaceae) |
| Lepanthes falx-bellica-12-F | Rondeletia buddleioides (Rubiaceae) |
| Lepanthes falx-bellica-12-F | Rondeletia buddleioides (Rubiaceae) |
| Lepanthes falx-bellica-12-F | Rondeletia buddleioides (Rubiaceae) |
| Lepanthes falx-bellica-12-F | Rondeletia buddleioides (Rubiaceae) |
| Lepanthes cribbii-12-F | Rondeletia buddleioides (Rubiaceae) |
| Lepanthes cribbii-12-F | Rondeletia buddleioides (Rubiaceae) |
| Lepanthes cribbii-12-F | Rondeletia buddleioides (Rubiaceae) |
| Lepanthes cribbii-12-F | Rondeletia buddleioides (Rubiaceae) |
| Lepanthes cribbii-12-F | Rondeletia buddleioides (Rubiaceae) |
| Lepanthes cribbii-12-F | Rondeletia buddleioides (Rubiaceae) |
| Lepanthes cribbii-12-F | Rondeletia buddleioides (Rubiaceae) |
| Lepanthes cribbii-12-F | Rondeletia buddleioides (Rubiaceae) |
| Lepanthes cribbii-12-F | Rondeletia buddleioides (Rubiaceae) |
| Lepanthes cribbii-12-F | Rondeletia buddleioides (Rubiaceae) |
| Lepanthes monteverdensis-12-F | Rondeletia buddleioides (Rubiaceae) |
| Lepanthes monteverdensis-12-F | Rondeletia buddleioides (Rubiaceae) |
| Lepanthes monteverdensis-12-F | Rondeletia buddleioides (Rubiaceae) |
| Lepanthes monteverdensis-12-F | Rondeletia buddleioides (Rubiaceae) |
| Lepanthes monteverdensis-12-F | Rondeletia buddleioides (Rubiaceae) |
| Lepanthes monteverdensis-12-F | Rondeletia buddleioides (Rubiaceae) |
| Lepanthes monteverdensis-12-F | Rondeletia buddleioides (Rubiaceae) |
| Lepanthes monteverdensis-12-F | Rondeletia buddleioides (Rubiaceae) |
| Lepanthes monteverdensis-12-F | Rondeletia buddleioides (Rubiaceae) |
| Lepanthes cribbii-13-G | Calyptranthes pittieri, cf. (Myrtaceae) |
| Lepanthes cribbii-13-G | Calyptranthes pittieri, cf. (Myrtaceae) |
| Lepanthes cribbii-13-G | Calyptranthes pittieri, cf. (Myrtaceae) |
| Lepanthes cribbii-13-G | Calyptranthes pittieri, cf. (Myrtaceae) |
| Lepanthes cribbii-13-G | Calyptranthes pittieri, cf. (Myrtaceae) |
| Lepanthes mentosa-16-J | Sapium rigidifolium (Euphorbiaceae) |
| Lepanthes mentosa-16-J | Sapium rigidifolium (Euphorbiaceae) |
| Lepanthes mentosa-16-J | Sapium rigidifolium (Euphorbiaceae) |
| Lepanthes mentosa-16-J | Sapium rigidifolium (Euphorbiaceae) |
| Lepanthes mentosa-16-J | Sapium rigidifolium (Euphorbiaceae) |
| Lepanthes mentosa-17-J | Trichilia havanensis (Meliaceae) |
| Lepanthes mentosa-17-J | Trichilia havanensis (Meliaceae) |
| Lepanthes mentosa-17-J | Trichilia havanensis (Meliaceae) |
| Lepanthes mentosa-17-J | Trichilia havanensis (Meliaceae) |
| Lepanthes mentosa-17-J | Trichilia havanensis (Meliaceae) |
| Lepanthes mentosa-17-J | Trichilia havanensis (Meliaceae) |
| Lepanthes cribbii-17-J | Trichilia havanensis (Meliaceae) |
| Lepanthes cribbii-17-J | Trichilia havanensis (Meliaceae) |
| Lepanthes cribbii-17-J | Trichilia havanensis (Meliaceae) |

| | |
|---|---|
| Lepanthes cribbii-17-J | Trichilia havanensis (Meliaceae) |
| Lepanthes cribbii-17-J | Trichilia havanensis (Meliaceae) |
| Lepanthes cribbii-17-J | Trichilia havanensis (Meliaceae) |
| Lepanthes cribbii-17-J | Trichilia havanensis (Meliaceae) |
| Lepanthes cribbii-17-J | Trichilia havanensis (Meliaceae) |
| Lepanthes cribbii-17-J | Trichilia havanensis (Meliaceae) |
| Lepanthes cribbii-17-J | Trichilia havanensis (Meliaceae) |
| Lepanthes falx-bellica-19-K | Cojoba costaricensis (Fabaceae) |
| Lepanthes falx-bellica-19-K | Cojoba costaricensis (Fabaceae) |
| Lepanthes falx-bellica-19-K | Cojoba costaricensis (Fabaceae) |
| Lepanthes falx-bellica-19-K | Cojoba costaricensis (Fabaceae) |
| Lepanthes falx-bellica-19-K | Cojoba costaricensis (Fabaceae) |
| Lepanthes falx-bellica-19-K | Cojoba costaricensis (Fabaceae) |
| Lepanthes falx-bellica-19-K | Cojoba costaricensis (Fabaceae) |
| Lepanthes falx-bellica-19-K | Cojoba costaricensis (Fabaceae) |
| Lepanthes cribbii-19-K | Cojoba costaricensis (Fabaceae) |
| Lepanthes cribbii-19-K | Cojoba costaricensis (Fabaceae) |
| Lepanthes cribbii-19-K | Cojoba costaricensis (Fabaceae) |
| Lepanthes cribbii-19-K | Cojoba costaricensis (Fabaceae) |
| Lepanthes cribbii-19-K | Cojoba costaricensis (Fabaceae) |
| Lepanthes cribbii-19-K | Cojoba costaricensis (Fabaceae) |
| Lepanthes cribbii-19-K | Cojoba costaricensis (Fabaceae) |
| Lepanthes cribbii-19-K | Cojoba costaricensis (Fabaceae) |
| Lepanthes cribbii-19-K | Cojoba costaricensis (Fabaceae) |
| Lepanthes cribbii-19-K | Cojoba costaricensis (Fabaceae) |
| Lepanthes falx-bellica-19-K | Cojoba costaricensis (Fabaceae) |
| Lepanthes falx-bellica-19-K | Cojoba costaricensis (Fabaceae) |
| Lepanthes falx-bellica-20-K | Miconia durandii (Melastomataceae) |
| Lepanthes falx-bellica-20-K | Miconia durandii (Melastomataceae) |
| Lepanthes falx-bellica-20-K | Miconia durandii (Melastomataceae) |
| Lepanthes falx-bellica-20-K | Miconia durandii (Melastomataceae) |
| Lepanthes falx-bellica-20-K | Miconia durandii (Melastomataceae) |
| Lepanthes falx-bellica-20-K | Miconia durandii (Melastomataceae) |
| Lepanthes falx-bellica-20-K | Miconia durandii (Melastomataceae) |
| Lepanthes falx-bellica-20-K | Miconia durandii (Melastomataceae) |
| Lepanthes falx-bellica-21-K | Eugenia austin-smithii (Myrtaceae) |
| Lepanthes falx-bellica-21-K | Eugenia austin-smithii (Myrtaceae) |
| Lepanthes falx-bellica-21-K | Eugenia austin-smithii (Myrtaceae) |
| Lepanthes falx-bellica-21-K | Eugenia austin-smithii (Myrtaceae) |
| Lepanthes falx-bellica-21-K | Eugenia austin-smithii (Myrtaceae) |
| Lepanthes falx-bellica-21-K | Eugenia austin-smithii (Myrtaceae) |
| Lepanthes falx-bellica-21-K | Eugenia austin-smithii (Myrtaceae) |
| Lepanthes falx-bellica-21-K | Eugenia austin-smithii (Myrtaceae) |
| Lepanthes falx-bellica-21-K | Eugenia austin-smithii (Myrtaceae) |

| | |
|---|---|
| Lepanthes falx-bellica-21-K | Eugenia austin-smithii (Myrtaceae) |
| Lepanthes cribbii-21-K | Eugenia austin-smithii (Myrtaceae) |
| Lepanthes cribbii-21-K | Eugenia austin-smithii (Myrtaceae) |
| Lepanthes cribbii-21-K | Eugenia austin-smithii (Myrtaceae) |
| Lepanthes cribbii-21-K | Eugenia austin-smithii (Myrtaceae) |
| Lepanthes cribbii-21-K | Eugenia austin-smithii (Myrtaceae) |
| Lepanthes cribbii-21-K | Eugenia austin-smithii (Myrtaceae) |
| Lepanthes cribbii-21-K | Eugenia austin-smithii (Myrtaceae) |
| Lepanthes cribbii-21-K | Eugenia austin-smithii (Myrtaceae) |
| Lepanthes cribbii-21-K | Eugenia austin-smithii (Myrtaceae) |
| Lepanthes cribbii-21-K | Eugenia austin-smithii (Myrtaceae) |
| Lepanthes falx-bellica-22-K | Monteverdia recondita (Celastraceae) |
| Lepanthes falx-bellica-22-K | Monteverdia recondita (Celastraceae) |
| Lepanthes falx-bellica-22-K | Monteverdia recondita (Celastraceae) |
| Lepanthes falx-bellica-22-K | Monteverdia recondita (Celastraceae) |
| Lepanthes falx-bellica-22-K | Monteverdia recondita (Celastraceae) |
| Lepanthes falx-bellica-22-K | Monteverdia recondita (Celastraceae) |
| Lepanthes falx-bellica-22-K | Monteverdia recondita (Celastraceae) |
| Lepanthes falx-bellica-23-K | Guatteria verrucosa (Annonaceae) |
| Lepanthes falx-bellica-23-K | Guatteria verrucosa (Annonaceae) |
| Lepanthes falx-bellica-23-K | Guatteria verrucosa (Annonaceae) |
| Lepanthes falx-bellica-23-K | Guatteria verrucosa (Annonaceae) |
| Lepanthes falx-bellica-23-K | Guatteria verrucosa (Annonaceae) |
| Lepanthes falx-bellica-23-K | Guatteria verrucosa (Annonaceae) |
| Lepanthes falx-bellica-23-K | Guatteria verrucosa (Annonaceae) |
| Lepanthes falx-bellica-23-K | Guatteria verrucosa (Annonaceae) |
| Lepanthes falx-bellica-24-K | Sapium rigidifolium (Euphorbiaceae) |
| Lepanthes falx-bellica-24-K | Sapium rigidifolium (Euphorbiaceae) |
| Lepanthes falx-bellica-24-K | Sapium rigidifolium (Euphorbiaceae) |
| Lepanthes falx-bellica-24-K | Sapium rigidifolium (Euphorbiaceae) |
| Lepanthes falx-bellica-24-K | Sapium rigidifolium (Euphorbiaceae) |
| Lepanthes falx-bellica-24-K | Sapium rigidifolium (Euphorbiaceae) |
| Lepanthes falx-bellica-24-K | Sapium rigidifolium (Euphorbiaceae) |
| Lepanthes falx-bellica-25-K | Sapium rigidifolium (Euphorbiaceae) |
| Lepanthes falx-bellica-25-K | Sapium rigidifolium (Euphorbiaceae) |
| Lepanthes falx-bellica-25-K | Sapium rigidifolium (Euphorbiaceae) |
| Lepanthes falx-bellica-25-K | Sapium rigidifolium (Euphorbiaceae) |
| Lepanthes falx-bellica-25-K | Sapium rigidifolium (Euphorbiaceae) |
| Lepanthes falx-bellica-25-K | Sapium rigidifolium (Euphorbiaceae) |
| Lepanthes falx-bellica-25-K | Sapium rigidifolium (Euphorbiaceae) |
| Lepanthes falx-bellica-25-K | Sapium rigidifolium (Euphorbiaceae) |
| Lepanthes falx-bellica-25-K | Sapium rigidifolium (Euphorbiaceae) |
| Lepanthes falx-bellica-26-L | Guarea kunthiana (Meliaceae) |
| Lepanthes falx-bellica-26-L | Guarea kunthiana (Meliaceae) |

| | |
|---|---|
| Lepanthes falx-bellica-26-L | Guarea kunthiana (Meliaceae) |
| Lepanthes falx-bellica-26-L | Guarea kunthiana (Meliaceae) |
| Lepanthes falx-bellica-26-L | Guarea kunthiana (Meliaceae) |
| Lepanthes falx-bellica-26-L | Guarea kunthiana (Meliaceae) |
| Lepanthes falx-bellica-26-L | Guarea kunthiana (Meliaceae) |
| Lepanthes falx-bellica-26-L | Guarea kunthiana (Meliaceae) |
| Lepanthes falx-bellica-26-L | Guarea kunthiana (Meliaceae) |
| Lepanthes falx-bellica-26-L | Guarea kunthiana (Meliaceae) |
| Lepanthes mentosa-27-L | Gonzalagunia rosea (Rubiaceae) |
| Lepanthes mentosa-27-L | Gonzalagunia rosea (Rubiaceae) |
| Lepanthes mentosa-27-L | Gonzalagunia rosea (Rubiaceae) |
| Lepanthes mentosa-27-L | Gonzalagunia rosea (Rubiaceae) |
| Lepanthes mentosa-27-L | Gonzalagunia rosea (Rubiaceae) |
| Lepanthes mentosa-27-L | Gonzalagunia rosea (Rubiaceae) |
| Lepanthes mentosa-27-L | Gonzalagunia rosea (Rubiaceae) |
| Lepanthes mentosa-27-L | Gonzalagunia rosea (Rubiaceae) |
| Lepanthes falx-bellica-28-M | Pouteria exfoliata (Sapotaceae) |
| Lepanthes falx-bellica-28-M | Pouteria exfoliata (Sapotaceae) |
| Lepanthes falx-bellica-28-M | Pouteria exfoliata (Sapotaceae) |
| Lepanthes falx-bellica-28-M | Pouteria exfoliata (Sapotaceae) |
| Lepanthes falx-bellica-28-M | Pouteria exfoliata (Sapotaceae) |
| Lepanthes falx-bellica-28-M | Pouteria exfoliata (Sapotaceae) |
| Lepanthes falx-bellica-28-M | Pouteria exfoliata (Sapotaceae) |
| Lepanthes falx-bellica-28-M | Pouteria exfoliata (Sapotaceae) |
| Lepanthes falx-bellica-28-M | Pouteria exfoliata (Sapotaceae) |
| Lepanthes falx-bellica-28-M | Pouteria exfoliata (Sapotaceae) |
| Lepanthes falx-bellica-29-M | Monteverdia recondita (Celastraceae) |
| Lepanthes falx-bellica-29-M | Monteverdia recondita (Celastraceae) |
| Lepanthes falx-bellica-29-M | Monteverdia recondita (Celastraceae) |
| Lepanthes falx-bellica-29-M | Monteverdia recondita (Celastraceae) |
| Lepanthes falx-bellica-29-M | Monteverdia recondita (Celastraceae) |
| Lepanthes falx-bellica-29-M | Monteverdia recondita (Celastraceae) |
| Lepanthes falx-bellica-29-M | Monteverdia recondita (Celastraceae) |
| Lepanthes falx-bellica-29-M | Monteverdia recondita (Celastraceae) |
| Lepanthes falx-bellica-29-M | Monteverdia recondita (Celastraceae) |
| Lepanthes falx-bellica-29-M | Monteverdia recondita (Celastraceae) |
| Lepanthes cribbii-30-M | Guarea rhopalocarpa (Meliaceae) |
| Lepanthes cribbii-30-M | Guarea rhopalocarpa (Meliaceae) |
| Lepanthes cribbii-30-M | Guarea rhopalocarpa (Meliaceae) |
| Lepanthes falx-bellica-30-M | Guarea rhopalocarpa (Meliaceae) |
| Lepanthes falx-bellica-30-M | Guarea rhopalocarpa (Meliaceae) |
| Lepanthes falx-bellica-30-M | Guarea rhopalocarpa (Meliaceae) |
| Lepanthes falx-bellica-30-M | Guarea rhopalocarpa (Meliaceae) |
| Lepanthes falx-bellica-30-M | Guarea rhopalocarpa (Meliaceae) |

| | |
|---|---|
| Lepanthes falx-bellica-30-M | Guarea rhopalocarpa (Meliaceae) |
| Lepanthes falx-bellica-30-M | Guarea rhopalocarpa (Meliaceae) |
| Lepanthes falx-bellica-30-M | Guarea rhopalocarpa (Meliaceae) |
| Lepanthes falx-bellica-31-M | PleurothyriumÂ palmanum (Lauraceae) |
| Lepanthes falx-bellica-31-M | PleurothyriumÂ palmanum (Lauraceae) |
| Lepanthes falx-bellica-31-M | PleurothyriumÂ palmanum (Lauraceae) |
| Lepanthes falx-bellica-31-M | PleurothyriumÂ palmanum (Lauraceae) |
| Lepanthes falx-bellica-31-M | PleurothyriumÂ palmanum (Lauraceae) |
| Lepanthes mentosa-32-L | Miconia durandii (Melastomataceae) |
| Lepanthes mentosa-32-L | Miconia durandii (Melastomataceae) |
| Lepanthes mentosa-32-L | Miconia durandii (Melastomataceae) |
| Lepanthes mentosa-32-L | Miconia durandii (Melastomataceae) |
| Lepanthes mentosa-32-L | Miconia durandii (Melastomataceae) |
| Lepanthes mentosa-32-L | Miconia durandii (Melastomataceae) |
| Lepanthes mentosa-32-L | Miconia durandii (Melastomataceae) |
| Lepanthes mentosa-32-L | Miconia durandii (Melastomataceae) |
| Lepanthes mentosa-32-L | Miconia durandii (Melastomataceae) |
| Lepanthes monteverdensis-33-L | Miconia oerstediana (Melastomataceae) |
| Lepanthes monteverdensis-33-L | Miconia oerstediana (Melastomataceae) |
| Lepanthes monteverdensis-33-L | Miconia oerstediana (Melastomataceae) |
| Lepanthes monteverdensis-33-L | Miconia oerstediana (Melastomataceae) |
| Lepanthes monteverdensis-33-L | Miconia oerstediana (Melastomataceae) |
| Lepanthes monteverdensis-33-L | Miconia oerstediana (Melastomataceae) |
| Lepanthes monteverdensis-33-L | Miconia oerstediana (Melastomataceae) |
| Lepanthes monteverdensis-33-L | Miconia oerstediana (Melastomataceae) |
| Lepanthes monteverdensis-33-L | Miconia oerstediana (Melastomataceae) |
| Lepanthes monteverdensis-33-L | Miconia oerstediana (Melastomataceae) |
| Lepanthes falx-bellica-34-N | Salacia petenensis (Hippocrateaceae) |
| Lepanthes falx-bellica-34-N | Salacia petenensis (Hippocrateaceae) |
| Lepanthes falx-bellica-34-N | Salacia petenensis (Hippocrateaceae) |
| Lepanthes falx-bellica-34-N | Salacia petenensis (Hippocrateaceae) |
| Lepanthes falx-bellica-34-N | Salacia petenensis (Hippocrateaceae) |
| Lepanthes falx-bellica-34-N | Salacia petenensis (Hippocrateaceae) |
| Lepanthes falx-bellica-34-N | Salacia petenensis (Hippocrateaceae) |
| Lepanthes falx-bellica-34-N | Salacia petenensis (Hippocrateaceae) |
| Lepanthes falx-bellica-35-N | Salacia petenensis (Hippocrateaceae) |
| Lepanthes falx-bellica-35-N | Salacia petenensis (Hippocrateaceae) |
| Lepanthes falx-bellica-35-N | Salacia petenensis (Hippocrateaceae) |
| Lepanthes falx-bellica-35-N | Salacia petenensis (Hippocrateaceae) |
| Lepanthes falx-bellica-35-N | Salacia petenensis (Hippocrateaceae) |
| Lepanthes falx-bellica-35-N | Salacia petenensis (Hippocrateaceae) |
| Lepanthes falx-bellica-35-N | Salacia petenensis (Hippocrateaceae) |
| Lepanthes falx-bellica-35-N | Salacia petenensis (Hippocrateaceae) |
| Lepanthes monteverdensis-36-N | Clusia sp. (Clusiaceae) |

| | |
|---|---|
| Lepanthes monteverdensis-36-N | Clusia sp. (Clusiaceae) |
| Lepanthes monteverdensis-36-N | Clusia sp. (Clusiaceae) |
| Lepanthes monteverdensis-36-N | Clusia sp. (Clusiaceae) |
| Lepanthes monteverdensis-36-N | Clusia sp. (Clusiaceae) |
| Lepanthes monteverdensis-36-N | Clusia sp. (Clusiaceae) |
| Lepanthes monteverdensis-36-N | Clusia sp. (Clusiaceae) |
| Lepanthes monteverdensis-36-N | Clusia sp. (Clusiaceae) |
| Lepanthes monteverdensis-36-N | Clusia sp. (Clusiaceae) |
| Lepanthes monteverdensis-36-N | Clusia sp. (Clusiaceae) |
| Lepanthes mentosa-37-L | Miconia oerstediana (Melastomataceae) |
| Lepanthes mentosa-37-L | Miconia oerstediana (Melastomataceae) |
| Lepanthes mentosa-37-L | Miconia oerstediana (Melastomataceae) |
| Lepanthes mentosa-37-L | Miconia oerstediana (Melastomataceae) |
| Lepanthes mentosa-37-L | Miconia oerstediana (Melastomataceae) |
| Lepanthes mentosa-37-L | Miconia oerstediana (Melastomataceae) |
| Lepanthes mentosa-37-L | Miconia oerstediana (Melastomataceae) |
| Lepanthes mentosa-37-L | Miconia oerstediana (Melastomataceae) |
| Lepanthes mentosa-37-L | Miconia oerstediana (Melastomataceae) |
| Lepanthes mentosa-37-L | Miconia oerstediana (Melastomataceae) |
| Lepanthes monteverdensis-38-L | Myrsine coriacea (Myrsinaceae) |
| Lepanthes monteverdensis-38-L | Myrsine coriacea (Myrsinaceae) |
| Lepanthes monteverdensis-38-L | Myrsine coriacea (Myrsinaceae) |
| Lepanthes monteverdensis-38-L | Myrsine coriacea (Myrsinaceae) |
| Lepanthes cribbii-38-L | Myrsine coriacea (Myrsinaceae) |
| Lepanthes falx-bellica-38-L | Myrsine coriacea (Myrsinaceae) |
| Lepanthes falx-bellica-38-L | Myrsine coriacea (Myrsinaceae) |
| Lepanthes falx-bellica-38-L | Myrsine coriacea (Myrsinaceae) |
| Lepanthes falx-bellica-38-L | Myrsine coriacea (Myrsinaceae) |
| Lepanthes falx-bellica-38-L | Myrsine coriacea (Myrsinaceae) |
| Lepanthes monteverdensis-39-L | Pouteria exfoliata (Sapotaceae) |
| Lepanthes monteverdensis-39-L | Pouteria exfoliata (Sapotaceae) |
| Lepanthes monteverdensis-39-L | Pouteria exfoliata (Sapotaceae) |
| Lepanthes monteverdensis-39-L | Pouteria exfoliata (Sapotaceae) |
| Lepanthes monteverdensis-39-L | Pouteria exfoliata (Sapotaceae) |
| Lepanthes monteverdensis-39-L | Pouteria exfoliata (Sapotaceae) |
| Lepanthes monteverdensis-39-L | Pouteria exfoliata (Sapotaceae) |
| Lepanthes monteverdensis-39-L | Pouteria exfoliata (Sapotaceae) |
| Lepanthes monteverdensis-39-L | Pouteria exfoliata (Sapotaceae) |
| Lepanthes monteverdensis-39-L | Pouteria exfoliata (Sapotaceae) |
| Lepanthes monteverdensis-40-L | Viburnum venustum (Viburnaceae) |
| Lepanthes monteverdensis-40-L | Viburnum venustum (Viburnaceae) |
| Lepanthes monteverdensis-40-L | Viburnum venustum (Viburnaceae) |
| Lepanthes monteverdensis-40-L | Viburnum venustum (Viburnaceae) |
| Lepanthes monteverdensis-40-L | Viburnum venustum (Viburnaceae) |

| | |
|---|---|
| Lepanthes monteverdensis-40-L | Viburnum venustum (Viburnaceae) |
| Lepanthes monteverdensis-40-L | Viburnum venustum (Viburnaceae) |
| Lepanthes monteverdensis-40-L | Viburnum venustum (Viburnaceae) |
| Lepanthes mentosa-41-O | Miconia conorufescens (Melastomataceae) |
| Lepanthes mentosa-41-O | Miconia conorufescens (Melastomataceae) |
| Lepanthes mentosa-41-O | Miconia conorufescens (Melastomataceae) |
| Lepanthes mentosa-41-O | Miconia conorufescens (Melastomataceae) |
| Lepanthes mentosa-41-O | Miconia conorufescens (Melastomataceae) |
| Lepanthes mentosa-41-O | Miconia conorufescens (Melastomataceae) |
| Lepanthes mentosa-41-O | Miconia conorufescens (Melastomataceae) |
| Lepanthes mentosa-41-O | Miconia conorufescens (Melastomataceae) |
| Lepanthes mentosa-41-O | Miconia conorufescens (Melastomataceae) |
| Lepanthes mentosa-41-O | Miconia conorufescens (Melastomataceae) |
| Lepanthes mentosa-38-L | Myrsine coriacea (Myrsinaceae) |
| Lepanthes mentosa-38-L | Myrsine coriacea (Myrsinaceae) |
| SynMock | SynMock |

| site | site letter | elevation m a.s.l. | longitude | latitude | origin |
|---|---|---|---|---|---|
| categorical | categorical | numeric | numeric | numeric | categorica |
| Alvaro's Finca (B) | B | 1470 | -84.81 | 10.33 | roots |
| Alvaro's Finca (B) | B | 1470 | -84.81 | 10.33 | roots |
| Alvaro's Finca (B) | B | 1470 | -84.81 | 10.33 | roots |
| Alvaro's Finca (B) | B | 1470 | -84.81 | 10.33 | roots |
| Alvaro's Finca (B) | B | 1470 | -84.81 | 10.33 | roots |
| Alvaro's Finca (B) | B | 1470 | -84.81 | 10.33 | roots |
| Alvaro's Finca (B) | B | 1470 | -84.81 | 10.33 | roots |
| Alvaro's Finca (B) | B | 1470 | -84.81 | 10.33 | roots |
| Alvaro's Finca (B) | B | 1470 | -84.81 | 10.33 | roots |
| Alvaro's Finca (B) | B | 1470 | -84.81 | 10.33 | roots |
| Alvaro's Finca (B) | B | 1502 | -84.81 | 10.33 | roots |
| Alvaro's Finca (B) | B | 1502 | -84.81 | 10.33 | roots |
| Alvaro's Finca (B) | B | 1502 | -84.81 | 10.33 | roots |
| Alvaro's Finca (B) | B | 1502 | -84.81 | 10.33 | roots |
| Alvaro's Finca (B) | B | 1502 | -84.81 | 10.33 | roots |
| Alvaro's Finca (B) | B | 1502 | -84.81 | 10.33 | roots |
| Alvaro's Finca (B) | B | 1502 | -84.81 | 10.33 | roots |
| Alvaro's Finca (B) | B | 1502 | -84.81 | 10.33 | roots |
| Alvaro's Finca (B) | B | 1502 | -84.81 | 10.33 | roots |
| Alvaro's Finca (B) | B | 1502 | -84.81 | 10.33 | roots |
| Alvaro's Finca (B) | B | 1502 | -84.81 | 10.33 | roots |
| Alvaro's Finca (B) | B | 1502 | -84.81 | 10.33 | roots |
| Alvaro's Finca (B) | B | 1502 | -84.81 | 10.33 | roots |
| Alvaro's Finca (B) | B | 1502 | -84.81 | 10.33 | roots |
| Alvaro's Finca (B) | B | 1502 | -84.81 | 10.33 | roots |
| Alvaro's Finca (B) | B | 1502 | -84.81 | 10.33 | roots |
| Alvaro's Finca (B) | B | 1502 | -84.81 | 10.33 | roots |
| Alvaro's Finca (B) | B | 1502 | -84.81 | 10.33 | roots |
| Alvaro's Finca (B) | B | 1502 | -84.81 | 10.33 | roots |
| Alvaro's Finca (B) | B | 1502 | -84.81 | 10.33 | roots |
| Trap's Lodge (C) | C | 1524 | -84.8 | 10.3 | roots |
| Trap's Lodge (C) | C | 1524 | -84.8 | 10.3 | roots |
| Trap's Lodge (C) | C | 1524 | -84.8 | 10.3 | roots |
| Trap's Lodge (C) | C | 1524 | -84.8 | 10.3 | roots |
| Trap's Lodge (C) | C | 1524 | -84.8 | 10.3 | roots |
| Trap's Lodge (C) | C | 1524 | -84.8 | 10.3 | roots |
| Trap's Lodge (C) | C | 1524 | -84.8 | 10.3 | roots |
| Trap's Lodge (C) | C | 1524 | -84.8 | 10.3 | roots |
| Trap's Lodge (C) | C | 1524 | -84.8 | 10.3 | roots |
| Trap's Lodge (C) | C | 1524 | -84.8 | 10.3 | roots |
| Trap's Lodge (C) | C | 1524 | -84.8 | 10.3 | roots |
| Trap's Lodge (C) | C | 1524 | -84.8 | 10.3 | roots |

| | | | | | |
|---|---|---|---|---|---|
| Trap's Lodge (C) | C | 1524 | -84.8 | 10.3 | roots |
| Trap's Lodge (C) | C | 1524 | -84.8 | 10.3 | roots |
| Trap's Lodge (C) | C | 1524 | -84.8 | 10.3 | roots |
| Cerro Amigos (D) | D | 1632 | -84.81 | 10.32 | roots |
| Cerro Amigos (D) | D | 1632 | -84.81 | 10.32 | roots |
| Cerro Amigos (D) | D | 1632 | -84.81 | 10.32 | roots |
| Cerro Amigos (D) | D | 1632 | -84.81 | 10.32 | roots |
| Cerro Amigos (D) | D | 1632 | -84.81 | 10.32 | roots |
| Cerro Amigos (D) | D | 1632 | -84.81 | 10.32 | roots |
| Cerro Amigos (D) | D | 1632 | -84.81 | 10.32 | roots |
| Cerro Amigos (D) | D | 1632 | -84.81 | 10.32 | roots |
| Cerro Amigos (D) | D | 1632 | -84.81 | 10.32 | roots |
| Cerro Amigos (D) | D | 1632 | -84.81 | 10.32 | roots |
| Cerro Amigos (D) | D | 1632 | -84.81 | 10.32 | roots |
| Cerro Amigos (E) | D | 1610 | -84.8 | 10.32 | roots |
| Cerro Amigos (E) | D | 1610 | -84.8 | 10.32 | roots |
| Cerro Amigos (E) | D | 1610 | -84.8 | 10.32 | roots |
| Cerro Amigos (E) | D | 1610 | -84.8 | 10.32 | roots |
| Cerro Amigos (E) | D | 1610 | -84.8 | 10.32 | roots |
| Cerro Amigos (E) | D | 1610 | -84.8 | 10.32 | roots |
| Cerro Amigos (E) | D | 1610 | -84.8 | 10.32 | roots |
| Cerro Amigos (E) | D | 1610 | -84.8 | 10.32 | roots |
| Cerro Amigos (E) | D | 1610 | -84.8 | 10.32 | roots |
| Cerro Amigos (E) | D | 1610 | -84.8 | 10.32 | roots |
| Cerro Amigos (E) | D | 1610 | -84.8 | 10.32 | roots |
| Cerro Amigos (E) | D | 1610 | -84.8 | 10.32 | roots |
| Cerro Amigos (E) | D | 1610 | -84.8 | 10.32 | roots |
| Cerro Amigos (E) | D | 1610 | -84.8 | 10.32 | roots |
| Cerro Amigos (E) | D | 1610 | -84.8 | 10.32 | roots |
| Cerro Amigos (E) | D | 1610 | -84.8 | 10.32 | roots |
| Cerro Amigos (E) | D | 1610 | -84.8 | 10.32 | roots |
| Cerro Amigos (F) | H | 1712 | -84.8 | 10.32 | roots |
| Cerro Amigos (F) | H | 1712 | -84.8 | 10.32 | roots |
| Cerro Amigos (F) | H | 1712 | -84.8 | 10.32 | roots |
| Cerro Amigos (F) | H | 1712 | -84.8 | 10.32 | roots |
| Cerro Amigos (F) | H | 1712 | -84.8 | 10.32 | roots |
| Cerro Amigos (F) | H | 1712 | -84.8 | 10.32 | roots |
| Cerro Amigos (F) | H | 1712 | -84.8 | 10.32 | roots |
| Cerro Amigos (F) | H | 1712 | -84.8 | 10.32 | roots |
| Cerro Amigos (F) | H | 1712 | -84.8 | 10.32 | roots |
| Cerro Amigos (F) | H | 1712 | -84.8 | 10.32 | roots |
| Cerro Amigos (F) | H | 1712 | -84.8 | 10.32 | roots |
| Cerro Amigos (F) | H | 1712 | -84.8 | 10.32 | roots |
| Cerro Amigos (F) | H | 1712 | -84.8 | 10.32 | roots |

| | | | | | |
|---|---|---|---|---|---|
| Cerro Amigos (F) | H | 1712 | -84.8 | 10.32 | roots |
| Cerro Amigos (F) | H | 1712 | -84.8 | 10.32 | roots |
| Cerro Amigos (F) | H | 1712 | -84.8 | 10.32 | roots |
| Cerro Amigos (F) | H | 1712 | -84.8 | 10.32 | roots |
| Cerro Amigos (F) | H | 1712 | -84.8 | 10.32 | roots |
| Cerro Amigos (F) | H | 1712 | -84.8 | 10.32 | roots |
| Cerro Amigos (F) | H | 1712 | -84.8 | 10.32 | roots |
| Cerro Amigos (F) | H | 1712 | -84.8 | 10.32 | roots |
| Cerro Amigos (F) | H | 1712 | -84.8 | 10.32 | roots |
| Cerro Amigos (F) | H | 1712 | -84.8 | 10.32 | roots |
| Cerro Amigos (F) | H | 1712 | -84.8 | 10.32 | roots |
| Cerro Amigos (F) | H | 1712 | -84.8 | 10.32 | roots |
| Cerro Amigos (F) | H | 1712 | -84.8 | 10.32 | roots |
| Cerro Amigos (F) | H | 1712 | -84.8 | 10.32 | roots |
| Cerro Amigos (F) | H | 1712 | -84.8 | 10.32 | roots |
| Cerro Amigos (F) | H | 1712 | -84.8 | 10.32 | roots |
| Cerro Amigos (F) | H | 1712 | -84.8 | 10.32 | roots |
| Cerro Amigos (F) | H | 1712 | -84.8 | 10.32 | roots |
| Cerro Amigos (F) | H | 1712 | -84.8 | 10.32 | roots |
| Cerro Amigos (F) | H | 1712 | -84.8 | 10.32 | roots |
| Cerro Amigos (F) | H | 1712 | -84.8 | 10.32 | roots |
| Cerro Amigos (F) | H | 1712 | -84.8 | 10.32 | roots |
| Cerro Amigos (F) | H | 1712 | -84.8 | 10.32 | roots |
| Cerro Amigos (F) | H | 1712 | -84.8 | 10.32 | roots |
| Cerro Amigos (F) | H | 1712 | -84.8 | 10.32 | roots |
| Cerro Amigos (F) | H | 1712 | -84.8 | 10.32 | roots |
| Cerro Amigos (F) | H | 1712 | -84.8 | 10.32 | roots |
| Cerro Amigos (F) | H | 1712 | -84.8 | 10.32 | roots |
| Cerro Amigos (F) | H | 1712 | -84.8 | 10.32 | roots |
| Cerro Amigos (F) | H | 1712 | -84.8 | 10.32 | roots |
| Cerro Amigos (F) | H | 1712 | -84.8 | 10.32 | roots |
| Cerro Amigos (F) | H | 1712 | -84.8 | 10.32 | roots |
| Cerro Amigos (F) | H | 1712 | -84.8 | 10.32 | roots |
| Cerro Amigos (F) | H | 1712 | -84.8 | 10.32 | roots |
| Cerro Amigos (F) | H | 1712 | -84.8 | 10.32 | roots |
| Cerro Amigos (F) | H | 1712 | -84.8 | 10.32 | roots |
| Cerro Amigos (F) | H | 1712 | -84.8 | 10.32 | roots |
| Cerro Amigos (F) | H | 1712 | -84.8 | 10.32 | roots |
| Cerro Amigos (F) | H | 1712 | -84.8 | 10.32 | roots |
| Cerro Amigos (F) | H | 1712 | -84.8 | 10.32 | roots |
| Cerro Amigos (F) | H | 1712 | -84.8 | 10.32 | roots |
| Cerro Amigos (F) | H | 1712 | -84.8 | 10.32 | roots |
| Cerro Amigos (F) | H | 1712 | -84.8 | 10.32 | roots |
| Cerro Amigos (F) | H | 1712 | -84.8 | 10.32 | roots |

| | | | | | |
|---|---|---|---|---|---|
| Cerro Amigos (F) | H | 1712 | -84.8 | 10.32 | roots |
| Cerro Amigos (F) | H | 1712 | -84.8 | 10.32 | roots |
| Cerro Amigos (F) | H | 1712 | -84.8 | 10.32 | roots |
| Cerro Amigos (F) | H | 1712 | -84.8 | 10.32 | roots |
| Cerro Amigos (F) | H | 1712 | -84.8 | 10.32 | roots |
| Cerro Amigos (F) | H | 1712 | -84.8 | 10.32 | roots |
| Cerro Amigos (F) | H | 1712 | -84.8 | 10.32 | roots |
| Cerro Amigos (F) | H | 1712 | -84.8 | 10.32 | roots |
| Cerro Amigos (F) | H | 1712 | -84.8 | 10.32 | roots |
| Cerro Amigos (F) | H | 1712 | -84.8 | 10.32 | roots |
| Cerro Amigos (F) | H | 1712 | -84.8 | 10.32 | roots |
| Cerro Amigos (F) | H | 1712 | -84.8 | 10.32 | roots |
| Cerro Amigos (F) | H | 1712 | -84.8 | 10.32 | roots |
| Cerro Amigos (F) | H | 1712 | -84.8 | 10.32 | roots |
| Cerro Amigos (F) | H | 1712 | -84.8 | 10.32 | roots |
| Cerro Amigos (F) | H | 1712 | -84.8 | 10.32 | roots |
| Cerro Amigos (F) | H | 1712 | -84.8 | 10.32 | roots |
| Cerro Amigos (F) | H | 1712 | -84.8 | 10.32 | roots |
| Cerro Amigos (F) | H | 1712 | -84.8 | 10.32 | roots |
| Cerro Amigos (F) | H | 1712 | -84.8 | 10.32 | roots |
| Cerro Amigos (F) | H | 1712 | -84.8 | 10.32 | roots |
| Cerro Amigos (F) | H | 1712 | -84.8 | 10.32 | roots |
| Cerro Amigos (F) | H | 1712 | -84.8 | 10.32 | roots |
| Cerro Amigos (F) | H | 1712 | -84.8 | 10.32 | roots |
| Cerro Amigos (F) | H | 1712 | -84.8 | 10.32 | roots |
| Cerro Amigos (G) | H | 1750 | -84.8 | 10.32 | roots |
| Cerro Amigos (G) | H | 1750 | -84.8 | 10.32 | roots |
| Cerro Amigos (G) | H | 1750 | -84.8 | 10.32 | roots |
| Cerro Amigos (G) | H | 1750 | -84.8 | 10.32 | roots |
| Cerro Amigos (G) | H | 1750 | -84.8 | 10.32 | roots |
| San Gerardo (J) | J | 1601 | -84.81 | 10.3 | roots |
| San Gerardo (J) | J | 1601 | -84.81 | 10.3 | roots |
| San Gerardo (J) | J | 1601 | -84.81 | 10.3 | roots |
| San Gerardo (J) | J | 1601 | -84.81 | 10.3 | roots |
| San Gerardo (J) | J | 1601 | -84.81 | 10.3 | roots |
| San Gerardo (J) | J | 1584 | -84.8 | 10.35 | roots |
| San Gerardo (J) | J | 1584 | -84.8 | 10.35 | roots |
| San Gerardo (J) | J | 1584 | -84.8 | 10.35 | roots |
| San Gerardo (J) | J | 1584 | -84.8 | 10.35 | roots |
| San Gerardo (J) | J | 1584 | -84.8 | 10.35 | roots |
| San Gerardo (J) | J | 1584 | -84.8 | 10.35 | roots |
| San Gerardo (J) | J | 1584 | -84.8 | 10.35 | roots |
| San Gerardo (J) | J | 1584 | -84.8 | 10.35 | roots |
| San Gerardo (J) | J | 1584 | -84.8 | 10.35 | roots |

| | | | | | |
|---|---|---|---|---|---|
| San Gerardo (J) | J | 1584 | -84.8 | 10.35 | roots |
| San Gerardo (J) | J | 1584 | -84.8 | 10.35 | roots |
| San Gerardo (J) | J | 1584 | -84.8 | 10.35 | roots |
| San Gerardo (J) | J | 1584 | -84.8 | 10.35 | roots |
| San Gerardo (J) | J | 1584 | -84.8 | 10.35 | roots |
| San Gerardo (J) | J | 1584 | -84.8 | 10.35 | roots |
| San Gerardo (J) | J | 1584 | -84.8 | 10.35 | roots |
| Selvatura (K) | K | 1620 | -84.8 | 10.34 | roots |
| Selvatura (K) | K | 1620 | -84.8 | 10.34 | roots |
| Selvatura (K) | K | 1620 | -84.8 | 10.34 | roots |
| Selvatura (K) | K | 1620 | -84.8 | 10.34 | roots |
| Selvatura (K) | K | 1620 | -84.8 | 10.34 | roots |
| Selvatura (K) | K | 1620 | -84.8 | 10.34 | roots |
| Selvatura (K) | K | 1620 | -84.8 | 10.34 | roots |
| Selvatura (K) | K | 1620 | -84.8 | 10.34 | roots |
| Selvatura (K) | K | 1620 | -84.8 | 10.34 | roots |
| Selvatura (K) | K | 1620 | -84.8 | 10.34 | roots |
| Selvatura (K) | K | 1620 | -84.8 | 10.34 | roots |
| Selvatura (K) | K | 1620 | -84.8 | 10.34 | roots |
| Selvatura (K) | K | 1620 | -84.8 | 10.34 | roots |
| Selvatura (K) | K | 1620 | -84.8 | 10.34 | roots |
| Selvatura (K) | K | 1620 | -84.8 | 10.34 | roots |
| Selvatura (K) | K | 1620 | -84.8 | 10.34 | roots |
| Selvatura (K) | K | 1620 | -84.8 | 10.34 | roots |
| Selvatura (K) | K | 1620 | -84.8 | 10.34 | roots |
| Selvatura (K) | K | 1620 | -84.8 | 10.34 | roots |
| Selvatura (K) | K | 1620 | -84.8 | 10.34 | roots |
| Selvatura (K) | K | 1620 | -84.8 | 10.34 | roots |
| Selvatura (K) | K | 1620 | -84.8 | 10.34 | roots |
| Selvatura (K) | K | 1620 | -84.8 | 10.34 | roots |
| Selvatura (K) | K | 1620 | -84.8 | 10.34 | roots |
| Selvatura (K) | K | 1620 | -84.8 | 10.34 | roots |
| Selvatura (K) | K | 1620 | -84.8 | 10.34 | roots |
| Selvatura (K) | K | 1620 | -84.8 | 10.34 | roots |
| Selvatura (K) | K | 1620 | -84.8 | 10.34 | roots |
| Selvatura (K) | K | 1620 | -84.8 | 10.34 | roots |
| Selvatura (K) | K | 1620 | -84.8 | 10.34 | roots |
| Selvatura (K) | K | 1620 | -84.8 | 10.34 | roots |
| Selvatura (K) | K | 1620 | -84.8 | 10.34 | roots |
| Selvatura (K) | K | 1620 | -84.8 | 10.34 | roots |
| Selvatura (K) | K | 1620 | -84.8 | 10.34 | roots |
| Selvatura (K) | K | 1620 | -84.8 | 10.34 | roots |
| Selvatura (K) | K | 1620 | -84.8 | 10.34 | roots |
| Selvatura (K) | K | 1620 | -84.8 | 10.34 | roots |

| | | | | | |
|---|---|---|---|---|---|
| Selvatura (K) | K | 1620 | -84.8 | 10.34 | roots |
| Selvatura (K) | K | 1620 | -84.8 | 10.34 | roots |
| Selvatura (K) | K | 1620 | -84.8 | 10.34 | roots |
| Selvatura (K) | K | 1620 | -84.8 | 10.34 | roots |
| Selvatura (K) | K | 1620 | -84.8 | 10.34 | roots |
| Selvatura (K) | K | 1620 | -84.8 | 10.34 | roots |
| Selvatura (K) | K | 1620 | -84.8 | 10.34 | roots |
| Selvatura (K) | K | 1620 | -84.8 | 10.34 | roots |
| Selvatura (K) | K | 1620 | -84.8 | 10.34 | roots |
| Selvatura (K) | K | 1620 | -84.8 | 10.34 | roots |
| Selvatura (K) | K | 1620 | -84.8 | 10.34 | roots |
| Selvatura (K) | K | 1620 | -84.8 | 10.34 | roots |
| Selvatura (K) | K | 1620 | -84.8 | 10.34 | roots |
| Selvatura (K) | K | 1620 | -84.8 | 10.34 | roots |
| Selvatura (K) | K | 1620 | -84.8 | 10.34 | roots |
| Selvatura (K) | K | 1620 | -84.8 | 10.34 | roots |
| Selvatura (K) | K | 1620 | -84.8 | 10.34 | roots |
| Selvatura (K) | K | 1620 | -84.8 | 10.34 | roots |
| Selvatura (K) | K | 1620 | -84.8 | 10.34 | roots |
| Selvatura (K) | K | 1620 | -84.8 | 10.34 | roots |
| Selvatura (K) | K | 1620 | -84.8 | 10.34 | roots |
| Selvatura (K) | K | 1620 | -84.8 | 10.34 | roots |
| Selvatura (K) | K | 1620 | -84.8 | 10.34 | roots |
| Selvatura (K) | K | 1620 | -84.8 | 10.34 | roots |
| Selvatura (K) | K | 1620 | -84.8 | 10.34 | roots |
| Selvatura (K) | K | 1620 | -84.8 | 10.34 | roots |
| Selvatura (K) | K | 1620 | -84.8 | 10.34 | roots |
| Selvatura (K) | K | 1620 | -84.8 | 10.34 | roots |
| Selvatura (K) | K | 1620 | -84.8 | 10.34 | roots |
| Selvatura (K) | K | 1620 | -84.8 | 10.34 | roots |
| Selvatura (K) | K | 1620 | -84.8 | 10.34 | roots |
| Selvatura (K) | K | 1620 | -84.8 | 10.34 | roots |
| Selvatura (K) | K | 1620 | -84.8 | 10.34 | roots |
| Selvatura (K) | K | 1620 | -84.8 | 10.34 | roots |
| Selvatura (K) | K | 1620 | -84.8 | 10.34 | roots |
| Selvatura (K) | K | 1620 | -84.8 | 10.34 | roots |
| Selvatura (K) | K | 1620 | -84.8 | 10.34 | roots |
| Selvatura (K) | K | 1620 | -84.8 | 10.34 | roots |
| Selvatura (K) | K | 1620 | -84.8 | 10.34 | roots |
| Selvatura (K) | K | 1620 | -84.8 | 10.34 | roots |
| Selvatura (K) | K | 1620 | -84.8 | 10.34 | roots |
| Selvatura (K) | K | 1620 | -84.8 | 10.34 | roots |
| Sky Adventures (L) | N | 1600 | -84.81 | 10.33 | roots |
| Sky Adventures (L) | N | 1600 | -84.81 | 10.33 | roots |

| | | | | | |
|---|---|---|---|---|---|
| Sky Adventures (L) | N | 1600 | -84.81 | 10.33 | roots |
| Sky Adventures (L) | N | 1600 | -84.81 | 10.33 | roots |
| Sky Adventures (L) | N | 1600 | -84.81 | 10.33 | roots |
| Sky Adventures (L) | N | 1600 | -84.81 | 10.33 | roots |
| Sky Adventures (L) | N | 1600 | -84.81 | 10.33 | roots |
| Sky Adventures (L) | N | 1600 | -84.81 | 10.33 | roots |
| Sky Adventures (L) | N | 1600 | -84.81 | 10.33 | roots |
| Sky Adventures (L) | N | 1600 | -84.81 | 10.33 | roots |
| Sky Adventures (L) | N | 1617 | -84.81 | 10.33 | roots |
| Sky Adventures (L) | N | 1617 | -84.81 | 10.33 | roots |
| Sky Adventures (L) | N | 1617 | -84.81 | 10.33 | roots |
| Sky Adventures (L) | N | 1617 | -84.81 | 10.33 | roots |
| Sky Adventures (L) | N | 1617 | -84.81 | 10.33 | roots |
| Sky Adventures (L) | N | 1617 | -84.81 | 10.33 | roots |
| Sky Adventures (L) | N | 1617 | -84.81 | 10.33 | roots |
| Sky Adventures (L) | N | 1617 | -84.81 | 10.33 | roots |
| Wilford Guindon trail (M) | U | 1564 | -84.79 | 10.3 | roots |
| Wilford Guindon trail (M) | U | 1564 | -84.79 | 10.3 | roots |
| Wilford Guindon trail (M) | U | 1564 | -84.79 | 10.3 | roots |
| Wilford Guindon trail (M) | U | 1564 | -84.79 | 10.3 | roots |
| Wilford Guindon trail (M) | U | 1564 | -84.79 | 10.3 | roots |
| Wilford Guindon trail (M) | U | 1564 | -84.79 | 10.3 | roots |
| Wilford Guindon trail (M) | U | 1564 | -84.79 | 10.3 | roots |
| Wilford Guindon trail (M) | U | 1564 | -84.79 | 10.3 | roots |
| Wilford Guindon trail (M) | U | 1564 | -84.79 | 10.3 | roots |
| Wilford Guindon trail (M) | U | 1564 | -84.79 | 10.3 | roots |
| Wilford Guindon trail (M) | U | 1579 | -84.79 | 10.3 | roots |
| Wilford Guindon trail (M) | U | 1579 | -84.79 | 10.3 | roots |
| Wilford Guindon trail (M) | U | 1579 | -84.79 | 10.3 | roots |
| Wilford Guindon trail (M) | U | 1579 | -84.79 | 10.3 | roots |
| Wilford Guindon trail (M) | U | 1579 | -84.79 | 10.3 | roots |
| Wilford Guindon trail (M) | U | 1579 | -84.79 | 10.3 | roots |
| Wilford Guindon trail (M) | U | 1579 | -84.79 | 10.3 | roots |
| Wilford Guindon trail (M) | U | 1579 | -84.79 | 10.3 | roots |
| Wilford Guindon trail (M) | U | 1579 | -84.79 | 10.3 | roots |
| Wilford Guindon trail (M) | U | 1579 | -84.79 | 10.3 | roots |
| Wilford Guindon trail (M) | U | 1583 | -84.79 | 10.3 | roots |
| Wilford Guindon trail (M) | U | 1583 | -84.79 | 10.3 | roots |
| Wilford Guindon trail (M) | U | 1583 | -84.79 | 10.3 | roots |
| Wilford Guindon trail (M) | U | 1583 | -84.79 | 10.3 | roots |
| Wilford Guindon trail (M) | U | 1583 | -84.79 | 10.3 | roots |
| Wilford Guindon trail (M) | U | 1583 | -84.79 | 10.3 | roots |
| Wilford Guindon trail (M) | U | 1583 | -84.79 | 10.3 | roots |
| Wilford Guindon trail (M) | U | 1583 | -84.79 | 10.3 | roots |

| | | | | | |
|---|---|---|---|---|---|
| Wilford Guindon trail (M) | U | 1583 | -84.79 | 10.3 | roots |
| Wilford Guindon trail (M) | U | 1583 | -84.79 | 10.3 | roots |
| Wilford Guindon trail (M) | U | 1583 | -84.79 | 10.3 | roots |
| Wilford Guindon trail (M) | U | 1619 | -84.79 | 10.3 | roots |
| Wilford Guindon trail (M) | U | 1619 | -84.79 | 10.3 | roots |
| Wilford Guindon trail (M) | U | 1619 | -84.79 | 10.3 | roots |
| Wilford Guindon trail (M) | U | 1619 | -84.79 | 10.3 | roots |
| Wilford Guindon trail (M) | U | 1619 | -84.79 | 10.3 | roots |
| Sky Adventures (L) | N | 1570 | -84.81 | 10.33 | roots |
| Sky Adventures (L) | N | 1570 | -84.81 | 10.33 | roots |
| Sky Adventures (L) | N | 1570 | -84.81 | 10.33 | roots |
| Sky Adventures (L) | N | 1570 | -84.81 | 10.33 | roots |
| Sky Adventures (L) | N | 1570 | -84.81 | 10.33 | roots |
| Sky Adventures (L) | N | 1570 | -84.81 | 10.33 | roots |
| Sky Adventures (L) | N | 1570 | -84.81 | 10.33 | roots |
| Sky Adventures (L) | N | 1570 | -84.81 | 10.33 | roots |
| Sky Adventures (L) | N | 1570 | -84.81 | 10.33 | roots |
| Sky Adventures (L) | N | 1576 | -84.81 | 10.33 | roots |
| Sky Adventures (L) | N | 1576 | -84.81 | 10.33 | roots |
| Sky Adventures (L) | N | 1576 | -84.81 | 10.33 | roots |
| Sky Adventures (L) | N | 1576 | -84.81 | 10.33 | roots |
| Sky Adventures (L) | N | 1576 | -84.81 | 10.33 | roots |
| Sky Adventures (L) | N | 1576 | -84.81 | 10.33 | roots |
| Sky Adventures (L) | N | 1576 | -84.81 | 10.33 | roots |
| Sky Adventures (L) | N | 1576 | -84.81 | 10.33 | roots |
| Sky Adventures (L) | N | 1576 | -84.81 | 10.33 | roots |
| Sky Adventures (L) | N | 1576 | -84.81 | 10.33 | roots |
| Sky Adventures (N) | N | 1574 | -84.81 | 10.33 | roots |
| Sky Adventures (N) | N | 1574 | -84.81 | 10.33 | roots |
| Sky Adventures (N) | N | 1574 | -84.81 | 10.33 | roots |
| Sky Adventures (N) | N | 1574 | -84.81 | 10.33 | roots |
| Sky Adventures (N) | N | 1574 | -84.81 | 10.33 | roots |
| Sky Adventures (N) | N | 1574 | -84.81 | 10.33 | roots |
| Sky Adventures (N) | N | 1574 | -84.81 | 10.33 | roots |
| Sky Adventures (N) | N | 1574 | -84.81 | 10.33 | roots |
| Sky Adventures (N) | N | 1574 | -84.81 | 10.33 | roots |
| Sky Adventures (N) | N | 1574 | -84.81 | 10.33 | roots |
| Sky Adventures (N) | N | 1574 | -84.81 | 10.33 | roots |
| Sky Adventures (N) | N | 1574 | -84.81 | 10.33 | roots |
| Sky Adventures (N) | N | 1574 | -84.81 | 10.33 | roots |
| Sky Adventures (N) | N | 1574 | -84.81 | 10.33 | roots |
| Sky Adventures (N) | N | 1574 | -84.81 | 10.33 | roots |
| Sky Adventures (N) | N | 1574 | -84.81 | 10.33 | roots |
| Sky Adventures (N) | N | 1591 | -84.81 | 10.33 | roots |

| | | | | | |
|---|---|---|---|---|---|
| Sky Adventures (N) | N | 1591 | -84.81 | 10.33 | roots |
| Sky Adventures (N) | N | 1591 | -84.81 | 10.33 | roots |
| Sky Adventures (N) | N | 1591 | -84.81 | 10.33 | roots |
| Sky Adventures (N) | N | 1591 | -84.81 | 10.33 | roots |
| Sky Adventures (N) | N | 1591 | -84.81 | 10.33 | roots |
| Sky Adventures (N) | N | 1591 | -84.81 | 10.33 | roots |
| Sky Adventures (N) | N | 1591 | -84.81 | 10.33 | roots |
| Sky Adventures (N) | N | 1591 | -84.81 | 10.33 | roots |
| Sky Adventures (N) | N | 1591 | -84.81 | 10.33 | roots |
| Sky Adventures (L) | N | 1556 | -84.81 | 10.33 | roots |
| Sky Adventures (L) | N | 1556 | -84.81 | 10.33 | roots |
| Sky Adventures (L) | N | 1556 | -84.81 | 10.33 | roots |
| Sky Adventures (L) | N | 1556 | -84.81 | 10.33 | roots |
| Sky Adventures (L) | N | 1556 | -84.81 | 10.33 | roots |
| Sky Adventures (L) | N | 1556 | -84.81 | 10.33 | roots |
| Sky Adventures (L) | N | 1556 | -84.81 | 10.33 | roots |
| Sky Adventures (L) | N | 1556 | -84.81 | 10.33 | roots |
| Sky Adventures (L) | N | 1556 | -84.81 | 10.33 | roots |
| Sky Adventures (L) | N | 1556 | -84.81 | 10.33 | roots |
| Sky Adventures (L) | N | 1551 | -84.81 | 10.33 | roots |
| Sky Adventures (L) | N | 1551 | -84.81 | 10.33 | roots |
| Sky Adventures (L) | N | 1551 | -84.81 | 10.33 | roots |
| Sky Adventures (L) | N | 1551 | -84.81 | 10.33 | roots |
| Sky Adventures (L) | N | 1551 | -84.81 | 10.33 | roots |
| Sky Adventures (L) | N | 1551 | -84.81 | 10.33 | roots |
| Sky Adventures (L) | N | 1551 | -84.81 | 10.33 | roots |
| Sky Adventures (L) | N | 1551 | -84.81 | 10.33 | roots |
| Sky Adventures (L) | N | 1551 | -84.81 | 10.33 | roots |
| Sky Adventures (L) | N | 1551 | -84.81 | 10.33 | roots |
| Sky Adventures (L) | N | 1562 | -84.81 | 10.33 | roots |
| Sky Adventures (L) | N | 1562 | -84.81 | 10.33 | roots |
| Sky Adventures (L) | N | 1562 | -84.81 | 10.33 | roots |
| Sky Adventures (L) | N | 1562 | -84.81 | 10.33 | roots |
| Sky Adventures (L) | N | 1562 | -84.81 | 10.33 | roots |
| Sky Adventures (L) | N | 1562 | -84.81 | 10.33 | roots |
| Sky Adventures (L) | N | 1562 | -84.81 | 10.33 | roots |
| Sky Adventures (L) | N | 1562 | -84.81 | 10.33 | roots |
| Sky Adventures (L) | N | 1562 | -84.81 | 10.33 | roots |
| Sky Adventures (L) | N | 1562 | -84.81 | 10.33 | roots |
| Sky Adventures (L) | N | 1563 | -84.81 | 10.33 | roots |
| Sky Adventures (L) | N | 1563 | -84.81 | 10.33 | roots |
| Sky Adventures (L) | N | 1563 | -84.81 | 10.33 | roots |
| Sky Adventures (L) | N | 1563 | -84.81 | 10.33 | roots |
| Sky Adventures (L) | N | 1563 | -84.81 | 10.33 | roots |

| | | | | | |
|---|---|---|---|---|---|
| Sky Adventures (L) | N | 1563 | -84.81 | 10.33 | roots |
| Sky Adventures (L) | N | 1563 | -84.81 | 10.33 | roots |
| Sky Adventures (L) | N | 1563 | -84.81 | 10.33 | roots |
| San Gerardo pasture (O) | J | 1482 | -84.81 | 10.35 | roots |
| San Gerardo pasture (O) | J | 1482 | -84.81 | 10.35 | roots |
| San Gerardo pasture (O) | J | 1482 | -84.81 | 10.35 | roots |
| San Gerardo pasture (O) | J | 1482 | -84.81 | 10.35 | roots |
| San Gerardo pasture (O) | J | 1482 | -84.81 | 10.35 | roots |
| San Gerardo pasture (O) | J | 1482 | -84.81 | 10.35 | roots |
| San Gerardo pasture (O) | J | 1482 | -84.81 | 10.35 | roots |
| San Gerardo pasture (O) | J | 1482 | -84.81 | 10.35 | roots |
| San Gerardo pasture (O) | J | 1482 | -84.81 | 10.35 | roots |
| San Gerardo pasture (O) | J | 1482 | -84.81 | 10.35 | roots |
| Sky Adventures (L) | N | 1551 | -84.81 | 10.33 | roots |
| Sky Adventures (L) | N | 1551 | -84.81 | 10.33 | roots |
| SynMock | SynMock | 0 | 0 | 0 | SynMock |

collection date
categorical
Jul-2019
Jul-2019
Jul-2019
Jul-2019
Jul-2019
Jul-2019
Jul-2019
Jul-2019
Jul-2019
Jul-2019
Jul-2019
Jul-2019
Jul-2019
Jul-2019
Jul-2019
Jul-2019
Jul-2019
Jul-2019
Jul-2019
Jul-2019
Jul-2019
Jul-2019
Jul-2019
Jul-2019
Jul-2019
Jul-2019
Jul-2019
Jul-2019
Jul-2019
Jul-2019
Jul-2019
Jul-2019
Jul-2019
Jul-2019
Jul-2019
Jul-2019
Jul-2019
Jul-2019
Jul-2019
Jul-2019
Jul-2019
Jul-2019

Jul-2019
Jul-2019
Jul-2019
Jul-2019
Jul-2019
Jul-2019
Jul-2019
Jul-2019
Jul-2019
Jul-2019
Jul-2019
Jul-2019
Jul-2019
Jul-2019
Jul-2019
Jul-2019
Jul-2019
Jul-2019
Jul-2019
Jul-2019
Jul-2019
Jul-2019
Jul-2019
Jul-2019
Jul-2019
Jul-2019
Jul-2019
Jul-2019
Jul-2019
Jul-2019
Jul-2019
Jul-2019
Jul-2019
Jul-2019
Jul-2019
Jul-2019
Jul-2019
Jul-2019
Jul-2019
Jul-2019
Jul-2019
Jul-2019
Jul-2019
Jul-2019

| |
|---|
| Jul-2019 |
| Jul-2019 |
| Jul-2019 |
| Jul-2019 |
| Jul-2019 |
| Jul-2019 |
| Jul-2019 |
| Jul-2019 |
| Jul-2019 |
| Jul-2019 |
| Jul-2019 |
| Jul-2019 |
| Jul-2019 |
| Jul-2019 |
| Jul-2019 |
| Jul-2019 |
| Jul-2019 |
| Jul-2019 |
| Jul-2019 |
| Jul-2019 |
| Jul-2019 |
| Jul-2019 |
| Jul-2019 |
| Jul-2019 |
| Jul-2019 |
| Jul-2019 |
| Jul-2019 |
| Jul-2019 |
| Jul-2019 |
| Jul-2019 |
| Jul-2019 |
| Jul-2019 |
| Jul-2019 |
| Jul-2019 |
| Jul-2019 |
| Jul-2019 |
| Jul-2019 |
| Jul-2019 |
| Jul-2019 |
| Jul-2019 |
| Jul-2019 |
| Jul-2019 |
| Jul-2019 |
| Jul-2019 |

| |
|---|
| Jul-2019 |
| Jul-2019 |
| Jul-2019 |
| Jul-2019 |
| Jul-2019 |
| Jul-2019 |
| Jul-2019 |
| Jul-2019 |
| Jul-2019 |
| Jul-2019 |
| Jul-2019 |
| Jul-2019 |
| Jul-2019 |
| Jul-2019 |
| Jul-2019 |
| Jul-2019 |
| Jul-2019 |
| Jul-2019 |
| Jul-2019 |
| Jul-2019 |
| Jul-2019 |
| Jul-2019 |
| Jul-2019 |
| Jul-2019 |
| Jul-2019 |
| Jul-2019 |
| Jul-2019 |
| Jul-2019 |
| Jul-2019 |
| Jul-2019 |
| Jul-2021 |
| Jul-2021 |
| Jul-2021 |
| Jul-2021 |
| Jul-2021 |
| Jul-2021 |
| Jul-2021 |
| Jul-2021 |
| Jul-2021 |
| Jul-2021 |
| Jul-2021 |
| Jul-2021 |
| Jul-2021 |
| Jul-2021 |

Jul-2021
Jul-2021
Jul-2021
Jul-2021
Jul-2021
Jul-2021
Jul-2021
Jul-2021
Jul-2021
Jul-2021
Jul-2021
Jul-2021
Jul-2021
Jul-2021
Jul-2021
Jul-2021
Jul-2021
Jul-2021
Jul-2021
Jul-2021
Jul-2021
Jul-2021
Jul-2021
Jul-2021
Jul-2021
Jul-2021
Jul-2021
Jul-2021
Jul-2021
Jul-2021
Jul-2021
Jul-2021
Jul-2021
Jul-2021
Jul-2021
Jul-2021
Jul-2021
Jul-2021
Jul-2021
Jul-2021
Jul-2021
Jul-2021
Jul-2021
Jul-2021

Jul-2021
Jul-2021
Jul-2021
Jul-2021
Jul-2021
Jul-2021
Jul-2021
Jul-2021
Jul-2021
Jul-2021
Jul-2021
Jul-2021
Jul-2021
Jul-2021
Jul-2021
Jul-2021
Jul-2021
Jul-2021
Jul-2021
Jul-2021
Jul-2021
Jul-2021
Jul-2021
Jul-2021
Jul-2021
Jul-2021
Jul-2021
Jul-2021
Jul-2021
Jul-2021
Jul-2021
Jul-2021
Jul-2021
Jul-2021
Jul-2021
Jul-2021
Jul-2021
Jul-2021
Jul-2021
Jul-2021
Jul-2021
Jul-2021
Jul-2021
Jul-2021

Jul-2021
Jul-2021
Jul-2021
Jul-2021
Jul-2021
Jul-2021
Jul-2021
Jul-2021
Jul-2021
Jul-2021
Jul-2021
Jul-2021
Jul-2021
Jul-2021
Jul-2021
Jul-2021
Jul-2021
Jul-2021
Jul-2021
Jul-2021
Jul-2021
Jul-2021
Jul-2021
Jul-2021
Jul-2021
Jul-2021
Jul-2021
Jul-2021
Jul-2021
Jul-2021
Jul-2021
Jul-2021
Jul-2021
Jul-2021
Jul-2021
Jul-2021
Jul-2021
Jul-2021
Jul-2021
Jul-2021
Jul-2021
Jul-2021
Jul-2021
Jul-2021

| |
|---|
| Jul-2021 |
| Jul-2021 |
| Jul-2021 |
| Jul-2021 |
| Jul-2021 |
| Jul-2021 |
| Jul-2021 |
| Jul-2021 |
| Jul-2021 |
| Jul-2021 |
| Jul-2021 |
| Jul-2021 |
| Jul-2021 |
| Jul-2021 |
| Jul-2021 |
| Jul-2021 |
| Jul-2021 |
| Jul-2021 |
| Jul-2021 |
| Jul-2021 |
| Jul-2021 |
| Jul-2021 |
| Jul-2021 |
| Jul-2021 |
| Jul-2021 |
| Jul-2021 |
| Jul-2021 |
| Jul-2021 |
| Jul-2021 |
| Jul-2021 |
| Jul-2021 |
| Jul-2021 |
| Jul-2021 |
| Jul-2021 |
| Jul-2021 |
| Jul-2021 |
| Jul-2021 |
| Jul-2021 |
| Jul-2021 |
| Jul-2021 |
| Jul-2021 |
| Jul-2021 |
| Jul-2021 |
| Jul-2021 |

| |
|---|
| Jul-2021 |
| Jul-2021 |
| Jul-2021 |
| Jul-2021 |
| Jul-2021 |
| Jul-2021 |
| Jul-2021 |
| Jul-2021 |
| Jul-2021 |
| Jul-2021 |
| Jul-2021 |
| Jul-2021 |
| Jul-2021 |
| Jul-2021 |
| Jul-2021 |
| Jul-2021 |
| Jul-2021 |
| Jul-2021 |
| Jul-2021 |
| Jul-2021 |
| Jul-2021 |
| Jul-2021 |
| Jul-2021 |
| Jul-2021 |
| Jul-2021 |
| Jul-2021 |
| Jul-2021 |
| Jul-2021 |
| Jul-2021 |
| Jul-2021 |
| Jul-2021 |
| Jul-2021 |
| Jul-2021 |
| Jul-2021 |
| Jul-2021 |
| Jul-2021 |
| Jul-2021 |
| Jul-2021 |
| Jul-2021 |
| Jul-2021 |
| Jul-2021 |
| Jul-2021 |
| Jul-2021 |
| Jul-2021 |

| |
|---|
| Jul-2021 |
| Jul-2021 |
| Jul-2021 |
| Jul-2021 |
| Jul-2021 |
| Jul-2021 |
| Jul-2021 |
| Jul-2021 |
| Jul-2021 |
| Jul-2021 |
| Jul-2021 |
| Jul-2021 |
| Jul-2021 |
| Jul-2021 |
| Jul-2021 |
| 0 |
